# The clustering of spatially associated species unravels patterns in Bornean tree species distributions

**DOI:** 10.1101/2022.09.13.507725

**Authors:** Sean E. H. Pang, J. W. Ferry Slik, Damaris Zurell, Edward L. Webb

## Abstract

Complex distribution data can be summarised by grouping species with similar or overlapping distributions to unravel patterns in species distributions and separate trends (e.g., of habitat loss) among spatially unique groups. However, such classifications are often heuristic, lacking the transparency, objectivity, and data-driven rigour of quantitative methods, which limits their interpretability and utility. Here, we develop and illustrate the clustering of spatially associated species, a methodological framework aimed at statistically classifying species using explicit measures of interspecific spatial association. We investigate several association indices and clustering algorithms and show how these methodological choices engender substantial variations in clustering outcome and performance. To facilitate robust decision making, we provide guidance on choosing methods appropriate to the study objective(s). As a case study, we apply the framework to model tree distributions in Borneo to evaluate the impact of land-cover change on separate species groupings. We identified 11 distinct clusters that unravelled ecologically meaningful patterns in Bornean tree distributions. These clusters then enabled us to quantify trends of habitat loss tied to each of those specific clusters, allowing us to discern particularly vulnerable species clusters and their distributions. This study demonstrates the advantages of adopting quantitatively derived clusters of spatially associated species and elucidates the potential of resultant clusters as a spatially explicit framework for investigating distribution-related questions in ecology, biogeography, and conservation. By adopting our methodological framework and publicly available codes, practitioners can leverage the ever-growing abundance of distribution data to better understand complex spatial patterns among species distributions and the disparate effects of global changes on biodiversity.

**Statement of authorship:** SEHP and ELW conceived the idea and designed methodology. SEHP conducted all analyses and developed the methodological framework with key inputs from ELW, JWFS, and DZ. All authors contributed to the interpretation of the results. SEHP and ELW wrote the first draft of the manuscript. All authors provided feedback on the writing.

## Introduction

With recent advancements in species distribution modelling (SDM) and the greater availability of biodiversity and environmental data, detailed predictions on species distributions are becoming increasingly accessible (Elith and Leathwick 2009, Norberg et al. 2019, Wüest et al. 2020, GBIF 2022). This new wealth of high-resolution data opens avenues for spatial analyses involving large species pools, which can serve to provide greater insights into community ecology and conservation science (Wüest et al. 2020, Hannah et al. 2020, Santini et al. 2021, Pang et al. 2021). However, as the number of species increases, so does the inherent complexity of the biogeographical data. Without a way to decompose species distributions, large stacks of distribution data may instead encumber analyses and obscure patterns or trends in the results (Marquet et al. 2004, Kreft and Jetz 2010, Villalobos et al. 2013). To better summarise and interpret complex spatial datasets, there is a need for robust methods for classifying species based on their distribution (Jongman et al. 1995, Legendre and Legendre 2012).

Species with highly similar or overlapped distributions indicate shared environmental requirements, biotic requirements, or dispersal barriers, or direct interactions between species (e.g., mutualism or predation) (Keddy 1992, Peterson 2011). Conversely, dissimilar distributions indicate differences in those processes instead (e.g., cold versus warm temperature requirements or competitive exclusion). Classifying species into groups based on their distribution therefore reflect a combination of processes—like the hierarchical filters of community assembly theory—that have led to recurrent patterns of associations or disassociations across geographic space (Shipley and Keddy 1987, Keddy 1992, Calatayud et al. 2020). Investigating these processes provides the grounding theory and hypotheses for understanding biodiversity patterns and species co-existence (Clements 1936, Shipley and Keddy 1987, Roxburgh and Chesson 1998, HilleRisLambers et al. 2012). In other words, by decomposing species into relatively homogeneous subsets, with shared geographic distributions, we can make apparent the spatial structure of species communities and their driving processes.

Classifying species with similar distributions is the “sister analysis” of classifying sites with similar compositions, i.e., the R-mode versus Q-mode analysis, respectively (Legendre and Legendre 2012). Although classifying sites is the more prevalent method in biogeography and spatial ecology (Kreft and Jetz 2010), classifying species offers an alternative view of spatial patterns. Site classifications are useful when the focus is on understanding compositional relationships among sites, e.g., how differences in species composition among areas might reflect historical biogeography and evolution or events of vicariance and geodispersal (Kreft and Jetz 2010, Holt et al. 2013, Hazzi et al. 2018, Leroy et al. 2019). Conversely, species classifications are useful when the focus is on understanding spatial relationships among species. For instance, consider the impact of deforestation and climate change on species distributions. As spatially heterogenous threats, their impact varies substantially depending on the species’ initial distribution (Bellard et al. 2014, Newbold 2018, Trisos et al. 2020, Pang et al. 2021). Such variations are difficult to separate and investigate in across species summaries that only reveal the most prevailing trend, whereas species-specific interpretations are impractical for studies involving hundreds or thousands of species (Torres et al. 2014, Marshall et al. 2018, Velazco et al. 2019). However, by grouping species and conducting group-specific summaries instead, we might uncover differing trends of loss and gain linked to each group’s distributional pattern and better understand their vulnerability to a given threat (e.g., lowland versus montane vulnerability to deforestation) (Manchego et al. 2017, Yanahan and Moore 2019, Pang et al. 2021). The classification of species with similar distributions therefore functions as concise summaries of species distribution data, which can provide a spatially explicit framework for investigating distribution-related questions in ecology, biogeography, and conservation.

Despite the potential usefulness of grouping spatially associated species, few studies have adopted replicable quantitative methods for doing so. Instead, studies often adopt a heuristic approach towards grouping species, using classifiers based on a putative understanding or description of species associations (Pompe et al. 2010, Manchego et al. 2017, Baatar 2019, Yanahan and Moore 2019). Such qualitative approaches lack the transparency, objectivity, and data-driven rigour of more quantitative methods, which limits their interpretability and utility (Kahneman and Tversky 1972, Jongman et al. 1995, Marquet et al. 2004, Legendre and Legendre 2012). In this regard, multivariate methods—based on explicit measures of interspecific spatial association—hold immense potential as a quantitative approach for unravelling patterns in species distributions and delineating species groupings (Jongman et al. 1995, Roxburgh and Chesson 1998, Legendre and Legendre 2012, Keil et al. 2021).

Multivariate methods can reduce the inherent complexity of biogeographical data, and their strength lies in their ability to generate statistically derived species groupings, with within- and between-cluster variances that are quantifiable and testable. The reproducibility, and therefore transparency, of such groupings is especially relevant given the prevalent use of SDMs for evaluating species vulnerabilities and informing management decisions (Guisan et al. 2013, Titeux et al. 2017, Feng et al. 2019, Zurell et al. 2020). The challenge with clustering spatially associated species, however, is that there is no definitive measure of spatial association; multiple interpretations and indices exist (Cramér 1924, Hubálek 1982, Roxburgh and Chesson 1998, Keil et al. 2021). Likewise, there are multiple strategies and techniques for unsupervised clustering but no single universal approach (Jongman et al. 1995, Legendre and Legendre 2012, Guerra et al. 2012, Erman et al. 2015, Seif 2018). Furthermore, such a technique has not been applied to detailed distribution data before (i.e., modelled distributions) and a methodological framework does not yet exist. Thus, practitioners face a suite of methodological options that can lead to highly varied clustering outcomes, but lack clear guidance on how to choose between outcomes or what the implications are. If clustering of spatially associated species is to be taken up more broadly, there is a need to understand how users’ methodological choices affect variations in clustering results and their subsequent ecological interpretations.

In this study, we develop and illustrate a methodological framework for the clustering of spatially associated species, which aims at helping practitioners navigate the steps involved with forming and applying species clusters to leverage the abundance of distribution data. To further guide user decision-making, we test several association indices and clustering algorithms and investigate resultant variations in clustering outcomes. As a case study, we apply the framework onto the modelled distribution of 390 tree species in Borneo. We then demonstrate the use of resultant clusters to separate trends of habitat loss due to land-cover change and further discuss other applications of spatially associated species-clusters.

## Methods

### Framework

We developed our framework based on those made by Kreft and Jetz (2010) and Dufrêne and Legendre (1997) for classifying sites—the “sister analysis” of classifying species. We also incorporated key considerations noted by Legendre and Legendre (2012) and Keil et al. (2021) for quantifying interspecific spatial associations and clustering species based on those associations. Our methodological framework for clustering spatially associated species involves six main steps (Fig. 1).

**Figure 1.**
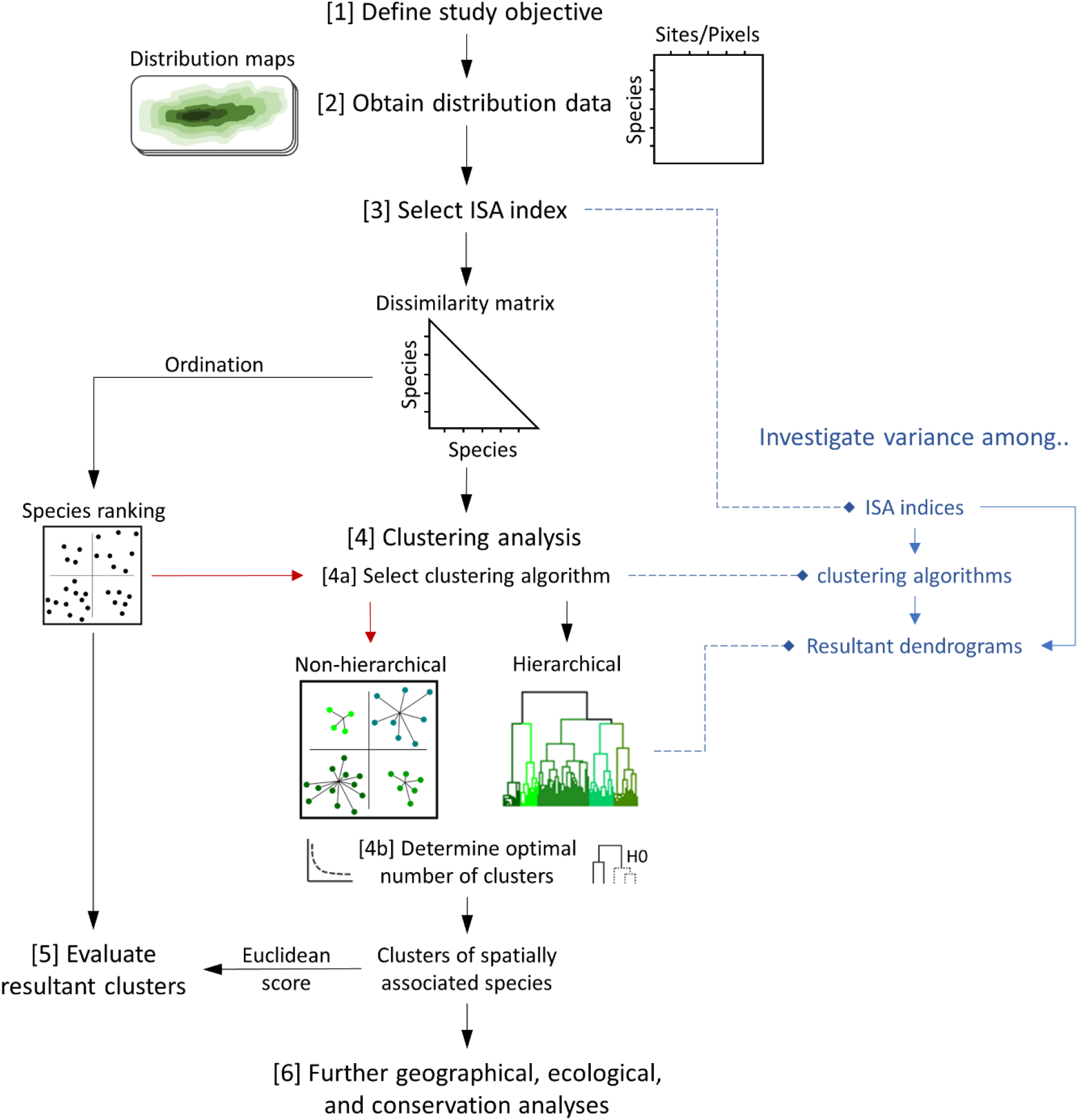
Methodological framework for the clustering of spatially associated species that consists of six main steps. The analyses of variance at different steps are included in blue. The red lines indicate the alternate route of applying non-hierarchical clustering algorithms, where the dissimilarity matrix is first ordinated.

1. Define the objective and purpose of the study. This sets the context and premise of the entire analysis and influences how subsequent steps are taken.
2. Obtain distribution data. The data type determines the spatial scale and extent at which associations are quantified (e.g., 50 m^2^ plot data vs. 10 km^2^ raster maps) and what further spatial analyses are possible (e.g., raster maps allow visualisations of each cluster’s distributional pattern); whereas the species list determines what patterns of associations can be found (e.g., montane patterns if only montane species are included).
3. Select an association index. A relevant index is selected to quantify interspecific spatial associations and produce a pairwise dissimilarity matrix as required for clustering.
4. Clustering analysis. A chosen clustering algorithm is applied to the dissimilarity matrix to cluster species. A stopping rule can be implemented to determine the optimal number of clusters, which often uses information on within-and between-cluster variances.
5. Evaluate resultant clusters. Clustering results can be evaluated quantitatively using a variety of metrics, each measuring a different aspect of clustering performance. Resultant clusters and their underlying dissimilarity matrix can also be visualised using ordination techniques for more qualitative comparisons of data structure and cluster performance.
6. Further geographical, ecological, and conservation analyses. Resultant clusters can be directly analysed (e.g., highlighting distinct patterns of spatial distributions) or used as a spatially explicit framework for further analyses (e.g., disentangling trends of habitat loss).

In the following, we elaborate on each of these steps and describe how we implemented them within the context of our case study. At steps (3) and (4), we also included analyses for investigating variations due to the choice of association index and clustering algorithm.

#### (1) Define the objective and purpose of the study

The objective of the study directly shapes how each subsequent step is taken. In selecting clustering strategies, for instance, one may choose between hierarchical or non-hierarchical clustering algorithms. If the only requirement of the clustering analysis is to form a given number of clusters for comparison with existing classifications (e.g., morphological adaptations; Boyce & Wong, 2019), discrete non-hierarchical clustering algorithms like K-means may be a good choice. On the other hand, if the focus is on investigating patterns of interspecific spatial associations and their relatedness to other interspecific relationships (e.g., phylogenetic or functional dissimilarity; Villalobos et al. 2017, Rüger et al. 2020), a hierarchical clustering algorithm is likely more appropriate. In our case study of Bornean tree species, we are interested in uncovering spatial relationships and identifying discrete clusters of similarly distributed species for further spatial analyses (i.e., habitat loss). Therefore, we focus on clustering algorithms that produce discrete hierarchical groupings of species (Kaufman and Rousseeuw 2005).

#### (2) Obtain distribution data

Species distribution data are the primary data required. The scale and extent of species distribution data, and the target species involved, all determine what can be inferred from the spatial analyses (Allen and Hoekstra 1990, Hurlbert and Jetz 2007, Kreft and Jetz 2010, Owen-Smith et al. 2015). However, more fundamentally, the type of distribution data itself affects how we interpret resultant clusters. There are three general types of distribution data: sampled, extent-of-occurrence, and modelled.

i. Sampled presence or abundance community data (e.g., plot data) give empirical information on species distributions. Their advantage is that they indicate directly observed patterns of associations across varying spatial and temporal scales (Owen-Smith et al. 2015, Ledo 2015). However, sampled data are often spatially biased and provide an incomplete representation of species distributions and their associations (Boakes et al. 2010, Stolar and Nielsen 2015).
ii. Extent-of-occurrence maps describe the maximum geographical extent of species and are typically hand drawn by experts using occurrence records, a heuristic understanding of species’ habitat requirements, or both (Gaston 1996, Lomolino et al. 2006). Extent-of-occurrence maps thus represent scale-dependent abstractions of species’ ranges (Gaston 2003, Hurlbert and Jetz 2007). Although extent-of-occurrence maps are highly qualitative and overestimate fine-scale occurrences (Graham and Hijmans 2006, Jetz et al. 2008), they are useful for broad-scale analyses where such data problems are less consequential (e.g., Kreft & Jetz, 2010; Trisos et al., 2020).
iii. Modelled data typically represent statistical inferences of species distributions derived from modelling occurrence records against prevailing environmental conditions (Soberon and Peterson 2005, Peterson and Soberón 2012) and are the data type of interest for this study. Modelled data are advantageous in that they can be used to examine, measure, and predict changes in species distributions across space and time and are frequently used to support biodiversity assessments and conservation prioritisations (Guisan and Thuiller 2005, Peterson 2011, Guisan et al. 2013). One inevitable limitation of this method is that the processes modelled to produce the data dictate what can be inferred. For instance, SDMs based on climate predictors alone cannot reflect variations in species distributions due to varying soil requirements (Corlett and Tomlinson 2020); and cannot be used to explicitly infer biotic interactions (i.e., modelled associations only reflect shared environmental requirements) (Peterson et al. 2020, Blanchet et al. 2020), although this can potentially be addressed if data on such interactions are incorporated (Tikhonov et al. 2017, Ovaskainen et al. 2017).

Our case study is of a regional scale, encompassing the island of Borneo, and aims to measure associations among tree species’ natural distributions (i.e., distributions predating anthropogenic disturbances). These distributions will be used to identify clusters of similarly distributed species and evaluate habitat loss due to land-cover change for each cluster. Modelled distributions were therefore the most appropriate choice as they allow fine-scale, statistical, and empirically based estimates of species distribution prior to anthropogenic disturbances like deforestation.

As our study focuses primarily on the methods for clustering spatially associated species, we present only a brief summary of the SDM methods (for full details, see Supporting Information S1-S4):

a. We obtained 19 bioclimatic (30 arcsec; Karger et al. 2017), 5 soil-water (30 arcsec; Trabucco and Zomer 2010, 2018), and 9 soil property (250 m^2^; Hengl et al. 2017) variables, resampled to 30 arcsec (~1 km^2^) and reduced using a principal component analysis (PCA) to their first 10 principal component (PC) axes (87% cumulative variance) via the ‘stats’ and ‘raster’ packages in R (Hijmans and Etten 2012, R Core Team 2013) (Supporting Information S2 and Table S1). A land-cover map (300 m^2^) from the year 1992 (earliest available) was obtained from the European Space Agency (ESA), reclassified as forested and non-forested following IPCC (Intergovernmental Panel on Climate Change) land categories and resampled to 30 arcsec (ESA 2017) (Supporting Information Table S1). A binary land-cover categorisation was adopted to provide a conservative approach towards identifying intact habitats; non-forested pixels were considered unsuitable. This was a reasonable assumption given our focus on tree species and that deforestation by definition entails clearing the land of all or most trees.
b. Occurrence data of vascular (Tracheophyta) species in Borneo were obtained from the Global Biodiversity Information Facility (GBIF) (Supporting Information S3). Spelling errors and synonyms were corrected using the ‘Taxonstand’ package in R (Cayuela et al. 2012, R Core Team 2013), and occurrences with low accuracy or precision were removed (Gueta and Carmel 2016) (Supporting Information S3). Tree species were identified using the GlobalTreeSearch database (Beech et al. 2017). To prevent mismatches between occurrences and contemporary land-cover data and account for anthropogenic niche truncations (Faurby and Araújo 2018, Milanesi et al. 2020, Pang et al. 2022), occurrences within forested and non-forested areas were separated following recommendations in Pang et al. (2022). While occurrences within forested areas were used for model training and cross-validation, occurrences within non-forested areas were used exclusively to validate historical distributions (i.e., projections onto a manually calibrated zero anthropogenic disturbance land-cover scenario). Occurrences were then thinned using a 10 km buffer (Aiello-Lammens et al. 2015), separately for each split. Species with fewer than 10 or 5 occurrences within forested or non-forested areas, respectively, were excluded.
c. All species were individually modelled and tuned using the MaxEnt (3.4.1) algorithm via the ‘ENMeval’ (2.0.0) package in R (Phillips et al. 2006, 2017, R Core Team 2013, Kass et al. 2021). To contrast occurrence data, 10,000 background points were sampled from pixels of ≥10 km distance from species’ occurrence points, where sampling probability was derived from a kernel density estimate representing the geographical sampling bias (VanDerWal et al. 2009, Kramer-Schadt et al. 2013, Vollering et al. 2019). For model tuning, 50 candidate models based on five combinations of feature classes and 10 regularisation multipliers were considered (Boria et al. 2017, Morales et al. 2017), each trained using occurrences within forested areas and the 10 environmental PC axes and reclassified land-cover map (year 1992) as predictors (Supporting Information S4 and Fig. S1). Candidate models were evaluated using a nested checkerboard cross-validation technique for species with >25 occurrences (else, a 10-fold cross-validation) and had their projections of historical distributions validated using occurrences within non-forested areas (testing for niche truncation; Pang et al. 2022). The best-performing candidate model was determined using a combination of Area Under the Curve (AUC), True Skill Statistics (TSS), and Omission Rates (OR), but accepted only if AUC > 0.7, TSS > 0.4 and OR < 0.2 (for cross-validation and previously excluded occurrences) (Supporting Information S4). Model projections of historical distributions as continuous estimates of habitat suitability (or probability of occurrence) were converted into binary range maps using the maximising the sum of sensitivity and specificity threshold (Liu et al. 2016).

#### (3) Select interspecific spatial association index

Clustering analyses require a distance/dissimilarity matrix. For clustering species distributions, we require a species-by-species matrix containing interspecific measures of distributional dissimilarity. To calculate the dissimilarity matrix, an appropriate association index must be selected. From a theoretical or conceptual standpoint, the choice of association index is arguably the most influential step, as the index used reflects the study’s mathematical and ecological definition of association (Box 1), and by extension, the clusters they inform (Hubálek 1982, Legendre and Legendre 2012, Keil et al. 2021).

##### Box 1. Significance of the choice of association index: the double-zero problem.

A simple example of how association indices can differ, relevant specifically to binary indices, is in their treatment of co-absences: this is known as the double-zero problem (Legendre and Legendre 2012). It is often difficult to sensibly define which sites of co-absence provide univocal or useful information for quantifying associations (Legendre and Legendre 2012). In extreme cases, the answer is clear, e.g., co-absences across Europe are not meaningful when measuring associations among Bornean species. But in many cases, the answer is less obvious. Binary indices are based on four quantities: number of sites uniquely occupied by species 1 (*b*) or species 2 (c), and number of sites occupied by both (*a*) or neither (*d*) species, where the total number of sites is *n* = *a* + *b* + *c* + *d*. As an example of the double-zero problem, compare the Jaccard (1901) index of association (*a*/(*a* + *b* + *c*)) against the Sokal & Michener (1958) Matching coefficient ((*a* + *d*)/*n*). As co-absences (*d*), and number of sites (*n*) by extension, becomes very large, the Jaccard index remains unchanged while the Matching coefficients in general approach a value of one. On the other hand, without co-absences (*d*), the index essentially ignores differences in species prevalence. Consider a map of 1000 pixels, two widespread species occupying 500 pixels each that co-occur in 250, and two range-restricted species occupying 50 pixels each that co-occur in 25. If we adopt the Jaccard index, associations between the two widespread and two range-restricted species are identical (widespread: 250/(250 + 250 + 250) = 0.33; range-restricted: 25/(25 + 25 + 25) = 0.33). However, it is generally considered “harder” for range-restricted than widespread species to co-occur, which the Matching coefficient reflects (widespread: (250 + 250)/1000 = 0.5; range-restricted: (25 + 925)/1000 = 0.95) (Legendre and Legendre 2012).

Keil et al. (2021) recently evaluated several association indices and found substantial variation in their performance and sensitivity, which provides vital information for selecting well-performing indices. However, within the context of clustering, we also need to consider how different indices affect the clustering outcome and distributional patterns identified. Moreover, it is unclear whether indices perform differently when using modelled distributions; Keil et al. (2021) used community matrices and simulated positions of individuals in a bound space. Thus, we selected and tested 12 indices from Keil et al. (2021)—six binary and six continuous—each representing various aspects of interspecific spatial association (Table 1). Broadly, the selected binary indices can be divided into two families: Jaccard index (jacc), Dice-Sorensen index (dice), and Alroy coefficient (alroy), which exclude co-absences; and Matching coefficient (match), Tetrachoric correlation (tetra), and Scaled C-score (scalec), which include co-absences. Correspondingly, continuous indices also vary in their mathematical properties and the dissimilarities they measure. Difference-based indices, Bray-Curtis (bray) and Ruzicka (ruz), which essentially measure the cumulative differences in occurrence probabilities across pixels; distance-based indices, Hellinger (hell) and Chi-squared (chi), which normalise probabilities by their sum before calculating differences; and correlation-based indices, Pearson (pears) and Spearman (spear), which examine the scaled covariances in probabilities (occurrence probabilities because of our use of modelled distribution data). Clustering analyses require non-negative distance or dissimilarity values, and so we transformed indices violating this requirement, e.g., (1 – pears) / 2 (see Table 1).

**Table 1.**
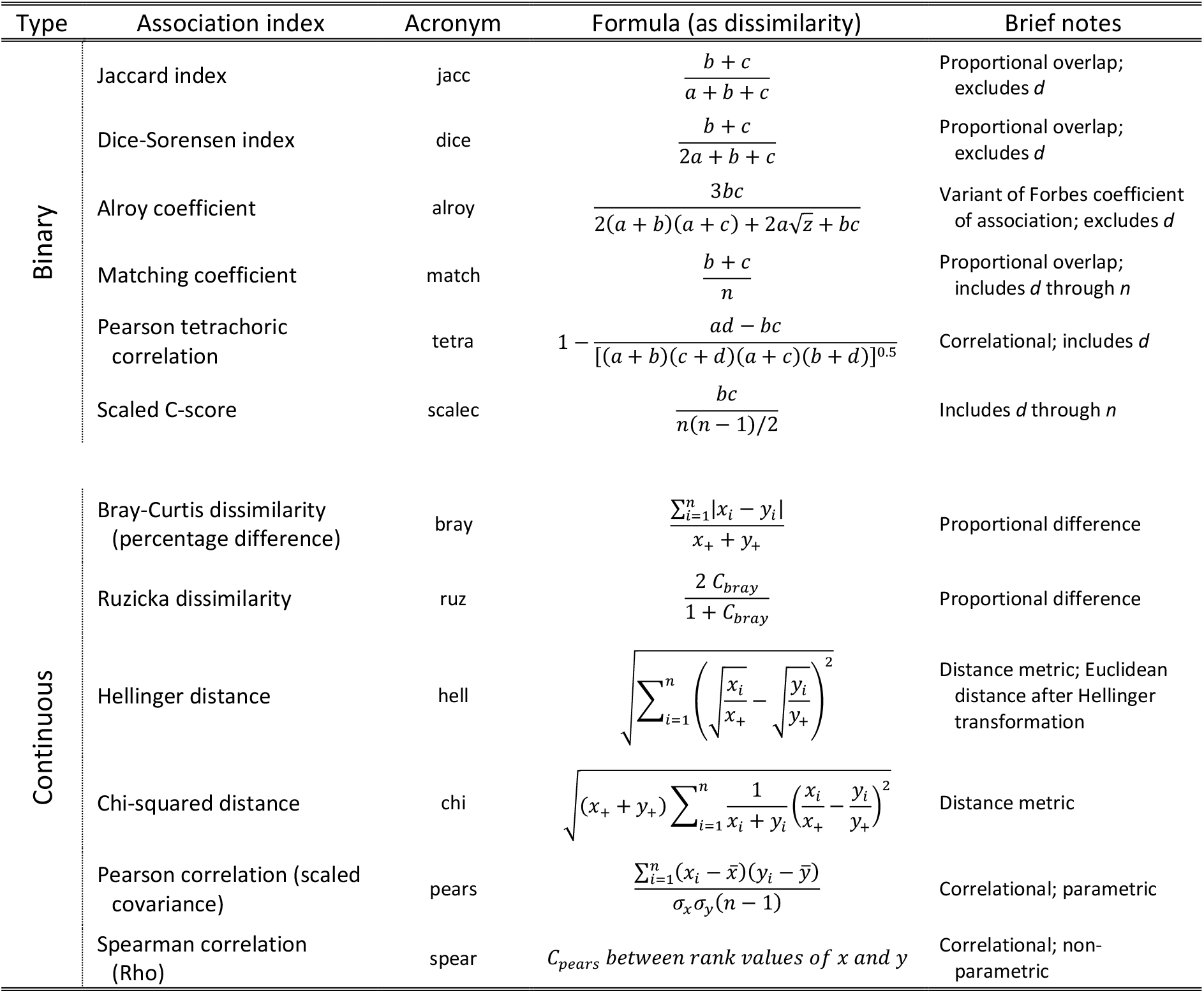
Continuous and binary indices of interspecific spatial associations tested for this study. Binary indices are based on four quantities: number of sites uniquely occupied by species 1 (*b*) or species 2 (*c*), and number of sites occupied by both (*a*) or neither (*d*) species, where the total number of sites is *n* = *a* + *b* + *c* + *d* and the total number of occupied sites is *z* = *a* + *b* + *c*. Continues indices are based on the vectors of continuous distribution data (occurrence probability or habitat suitability) of two species as represented by *x* and *y*, their means as 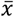 and 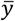, their sums as *x*_+_ and *y*_+_, and their standard deviations as *σ_x_* and *σ_y_*, where *x_i_* and *y_i_* are their values at site *i* and *n* equals the total number of sites.

These indices were applied to the modelled historical distribution of our 390 tree species, which resulted in 12 dissimilarity matrices of dimension 390 × 390, each corresponding to one index. While continuous indices used continuous distribution maps, binary indices used their binary equivalents. To examine variations in interspecific spatial association due to the choice of index, we calculated the Pearson correlation of association values among the 12 dissimilarity matrices. The correlation matrix was then passed into a PCA and visualised using a variable plot.

#### (4) Clustering analysis

Cluster analysis falls under the family of unsupervised learning methods in exploratory data analysis and its primary aim is to classify similar objects into groups while identifying boundaries between groups (Kaufman and Rousseeuw 2005). Before applying clustering, one needs to justify why discontinuities might exist or explain a practical need to divide a continuous set of objects into groups (Legendre and Legendre 2012). Justifying discontinuities among species distributions is complex. The community-unit model by Clements (1936) states that communities result from non-overlapping groups of species-response curves along an environmental gradient, which supports clearly discrete groupings of species distributions. In contrast, the continuum model of Whittaker (1951, 1953) and Curtis (1959) contends that communities vary gradually along complex environmental gradients, and that no distinct groupings exist. Other studies suggest that neither of these two views is correct, or some amalgamation of them (Westman 1985, Roberts 1987, Shipley and Keddy 1987), or that it depends on the scale (Allen and Hoekstra 1990, Hoekstra et al. 1991, Collins et al. 1993). However, evidence does support recurrent patterns of distribution along environmental gradients for Bornean flora (Slik et al. 2003, 2009, Raes et al. 2009), which clustering analyses may serve to uncover. More practically, clustering species distributions provides a spatially explicit framework for investigating distribution-related questions and applications in ecology, biogeography, and conservation. For this study, cluster-specific summaries of habitat loss would offer greater insights into trends of biodiversity loss in Borneo.

##### (4a) Select clustering algorithm

Two main families of clustering algorithms exist: non-hierarchical and hierarchical (Jain et al. 1999, Kaufman and Rousseeuw 2005). Non-hierarchical algorithms partition the data into a pre-determined number of clusters (*k*). Algorithms from this family include *k*-means and partitioning around medoids (Kaufman and Rousseeuw 2005). However, non-hierarchical algorithms are limited because they require the user to specify the number of clusters and they do not yield relationships among clusters (Legendre and Legendre 2012). Thus, we do not consider non-hierarchical algorithms further. By contrast, hierarchical algorithms construct a hierarchy of clusters, where a pre-determined number of clusters is not required and relationships among clusters are depicted through a dendrogram. Hierarchical relationships are especially relevant for spatially associated ecological communities (Clements 1936, Keddy 1992, Collins et al. 1993).

Hierarchical algorithms fall into two main categories: agglomerative and divisive (Jain et al. 1999, Kaufman and Rousseeuw 2005). Agglomerative algorithms start with each object as its own group and gradually merge the most similar objects (groups)—according to the linkage function—until all objects form one group. Divisive algorithms take the opposite approach and start with all objects as one group and continuously partition groups into two least-similar subsets until each object forms its own group. Although divisive algorithms are typically more efficient and accurate, they are also more complex and harder to interpret, whereas agglomerative algorithms rely on simpler merging steps and are also among the most popular (Rajalingam and Ranjini 2011, Singh and Singh 2012, Erman et al. 2015, Roux 2018). Thus, we focus on seven easy-to-implement and frequently used agglomerative clustering algorithms (Table 2): unweighted pair-group method using arithmetic averages (UPGMA), weighted pair-group method using arithmetic averages (WPGMA), complete linkage (CL), single linkage (SL), unweighted pair-group method using centroids (UPGMC), weighted pair-group method using centroids (WPGMC), and Ward’s method (WARD). Clustering algorithms were applied to each dissimilarity matrix using the ‘linkage’ function from the *mdendro* package in R, generating 84 candidate clustering outcomes (12 dissimilarity indices by seven clustering algorithms) and their dendrograms (R Core Team 2013, Fernández and Gómez 2020).

**Table 2.**
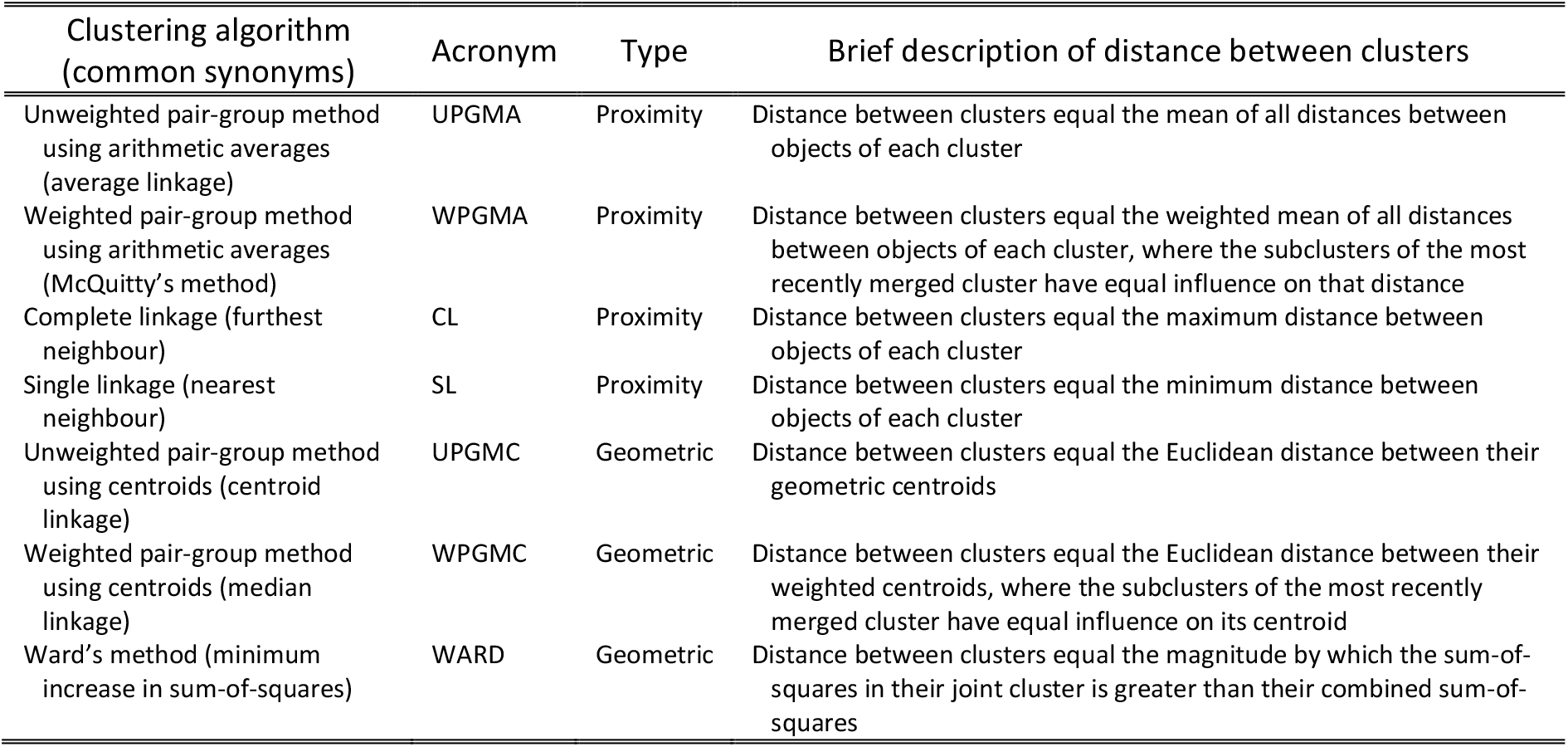
The seven hierarchical agglomerative clustering algorithms tested for this study.

To investigate variations among the 84 candidate dendrograms, we measured variations in dendrogram structure using Baker’s Gamma. Baker’s Gamma essentially compares the relative position of nodes between dendrograms as the Spearman correlation in lowest common branches, where the lowest common branch is the highest possible number of clusters for which two species belong to the same cluster (i.e., merging node in relation to other nodes) (Baker 1974). The correlation values between dendrograms were then passed into a PCA and visualised using variable plots. We also compared dendrograms using co-phenetic correlation, common nodes, and Full-text index in Minute space (FM-index) (Supporting Information Fig. S5-S12). However, we focused on Baker’s Gamma as it assessed the general structure of each dendrogram and is unaffected by the height of each branch, because branch heights may be distorted for dendrograms resulting from space-dilating clustering algorithms like WARD (Fernández and Gómez 2020).

##### (4b) Determine optimal number of clusters

Determining the optimal number of clusters is an age-old challenge of clustering analysis (Milligan and Cooper 1985, Chouikhi et al. 2015). While taxonomists often seek naturally formed clusters with small distances between member objects and large distances between objects from different clusters, ecologists seek to understand a world that often exists along a continuum and must usually contend with somewhat arbitrary clusters (Gauch and Whittaker 1981, Legendre and Legendre 2012). Clusters may be arbitrary when objects are evenly spread through dissimilarity space. In such cases, groupings are partially imposed by the clustering algorithm and are less intrinsic to the data, and a range of optimal number of clusters may be more appropriate than a single value (Gauch and Whittaker 1981). This does not imply, however, that a stopping rule or an internal validation criterion for determining cluster boundaries is unnecessary. On the contrary, a more rigorous selection procedure is required to ensure transparency, objectivity, and replicability in determining and justifying the (range of) optimal number of clusters (Legendre and Legendre 2012, Guerra et al. 2012).

Quantitative inspections of diagnostic graphs (i.e., an evaluation metric plotted against the number of clusters) offers a rigorous and data-driven procedure towards determining a meaningful and useful number of clusters (Milligan and Cooper 1985, Salvador and Chan 2004, e.g., Kreft and Jetz 2010). We assessed the diagnostic graph of three evaluation metrics: merging height, within-cluster variance, and between-cluster variance. Plotting these metrics against the number of clusters produced a scree-like evaluation plot (Supporting Information Fig. S2). The L-method of Salvador & Chan (2004) was used to quantitatively identify the knee or elbow in these evaluation plots, that is, the maximum curvature of the graph (for details, see Salvador and Chan 2004). Although other advanced stopping rules certainly exist, many of them are difficult to implement for ecological data where clusters are largely arbitrary and filled with outliers and noise (Milligan and Cooper 1985, Legendre and Legendre 2012, Guerra et al. 2012, Chouikhi et al. 2015). The Caliński & Harabasz (1974) index, Duda & Hart (1973) ratio criteria, Hubert & Levin (1976) *C*-index, and Rousseeuw (1987) silhouettes identified optimal number of clusters that were less meaningful and useful (see Supporting Information Table S3). Hence, to complement inspections of diagnostic graphs, we developed and employed a bifurcation paired t-test stopping rule. Moving down the dendrogram, we tested for a significant decrease in within-cluster variance using a paired t-test at each passing bifurcation (i.e., cluster partitioning). The bifurcation for which no significant decrease was observed determined the stopping point for partitioning the data, and therefore the optimal number of clusters (for details, see Supporting Information Fig. S3). For stopping rules involving within- and between-cluster variance, the centres used to calculate variances were either the aggregated (centroid; mean for continuous data and mode for binary data) or indicator species distribution of each cluster (medoid; object with the lowest sum of within-cluster dissimilarities).

#### (5) Evaluate resultant clusters

We propose a set of quantitative and qualitative evaluations for assessing clustering performance. First, we quantitatively assessed the dendrogram of each candidate clustering outcome using three dendrogram performance metrics:

1. Co-phenetic correlation measures faithfulness of the resultant co-phenetic distances (dendrogram branch heights) to the original dissimilarity matrix (Sokal and Rohlf 1962).
2. Agglomerative coefficient measures the strength of resulting clusters (Rousseeuw 1985).
3. Tree balance measures the equality in number of objects between clusters at each merger or partition (Fernández and Gómez 2020).

A min-max scaling was then applied to each metric and combined using the Euclidean formula, where the Euclidean score quantifies the dendrogram’s performance as a distance from an origin representing the worst performance possible. We used the Euclidean formula because of its flexibility, where metrics can be added, removed, or weighted, depending on the clustering characteristic(s) deemed most relevant to the study objective.

While the raw score of each metric covered specific aspects of a dendrogram’s performance, the combined score provides an overview of its performance, aiding the selection of the most appropriate dendrogram. Although internal validation criteria could also provide useful information on cluster performances, we focused on dendrogram performance metrics because they evaluated the performance of the entire clustering result rather than a defined set of clusters (Milligan and Cooper 1985, Legendre and Legendre 2012, Fernández and Gómez 2020). Dendrogram metrics were therefore consistent measures of clustering performance, indifferent to the number of clusters selected. This was vital because the relationship between clusters across the hierarchy was an important facet of the clustering result that needed to be assessed; and because exact cluster boundaries are less crucial when dealing with arbitrary clusters (Gauch and Whittaker 1981, Legendre and Legendre 2012).

Second, we performed non-metric multidimensional scaling (NMDS) on each dissimilarity matrix to visualise species distributions in ordination space. Ordination is a widely used tool for projecting multivariate data into low-dimensional space, where (in our case) species are arranged along reduced axes of geographic distributions (Legendre and Legendre 2012). NMDS is regarded as the most robust unconstrained method and most effective at reducing complex data (Minchin 1987, Legendre and Legendre 2012). Additionally, NMDS requires no underlying assumption about linearity or normality, in that any distance or dissimilarity matrix can be used (Ludwig et al. 1988). Ordination via NMDS therefore represents a useful approach for visualising distributional dissimilarities and investigating the spatial structure of community data. Paired alongside their respective dendrograms, NMDS ordinations also provide insight into the formation of cluster boundaries, thereby aiding interpretations of cluster memberships and hierarchical relationships. We performed the NMDS using the ‘metaMDS’ function from the *vegan* package in R, with 100 random starts based on a fixed initial seed (Minchin 1987, Oksanen et al. 2007, R Core Team 2013).

#### (6) Further geographical, ecological and conservation analyses

The representative distribution of each cluster was generated by summing the binary data of member species at each pixel and dividing values by the total number of member species (i.e., proportion of member species present). The distribution can be used to visually determine if clusters are ecologically meaningful and justifiable (Peterson 2011, Di Febbraro et al. 2018, Mainali et al. 2020), to identify consistencies among clustering outcomes, or as a spatially explicit framework for further spatial analyses (Keddy 1992, Roxburgh and Chesson 1998, Currie 2019). Binary distributions were used because variable sampling biases, species prevalence or rarity, and assumptions of occurrence probability across species meant that continuous distributions are debatably non-comparable and cannot be combined (Phillips et al. 2006, Elith and Leathwick 2009, Elith et al. 2011, Merow et al. 2013). Moreover, many applications of SDM require binary outputs, and binary distributions are easier to interpret than their continuous counterpart (Guisan and Thuiller 2005, Royle et al. 2012, Fithian and Hastie 2013, Liu et al. 2016).

As a demonstration of the application of distribution-based species clusters, representative distributions from the final clustering outcome were used to assess cluster-specific habitat loss due to land-cover change. Annual, 300 m^2^, land-cover maps for the years 1992 to 2020 (ESA 2017) were similarly reclassified to forested and non-forested and resampled to 30 arcsec as in step 2, and overlayed onto each representative distribution; habitat loss occurred when a pixel transitioned from forest to non-forest. For each cluster, habitat availability was quantified as the sum of representative distribution values (i.e., proportion of member species present) within forested pixels, such that pixels with higher proportion values were weighted higher and non-forested/deforested pixels were valued at zero. We then calculated and presented habitat loss for each cluster in three ways: (1) the percentage of historically available habitat lost by 1992, lost between 1992 to 2020, and remaining in the year 2020, (2) annual percentages of 1992 habitats remaining from 1992 to 2020, and (3) annual rates of habitat loss from 1993 to 2020 as a percentage of available habitats in the year before. Historical baselines assumed all pixels were forested and reforested pixels were not considered.

## Results

Of the 743 species with sufficient occurrence data, we accepted the SDM of 390. Accepted models had an average AUC of 0.76, TSS of 0.52, and OR of 0.10 and 0.04 for cross-validated occurrences and excluded occurrences (i.e., within non-forested pixels), respectively.

### Variance among association indices

We found association indices to generally capture one of three aspects of interspecific spatial association, as reflected by the three distinct groupings in our PCA of association values (Fig. 2a). The first group consisted of indices that measure association as differences (bray, ruz) or distances (hell, chi) in site values (i.e., occurrence probabilities), which also formed the tightest and most distinct group. The second group consisted of binary indices that exclude co-absences (jacc, dice, alroy). The last group consisted of two continuous indices that measure association as the correlation in site values (pears, spear) and three binary indices that include co-absences.

**Figure 2.**
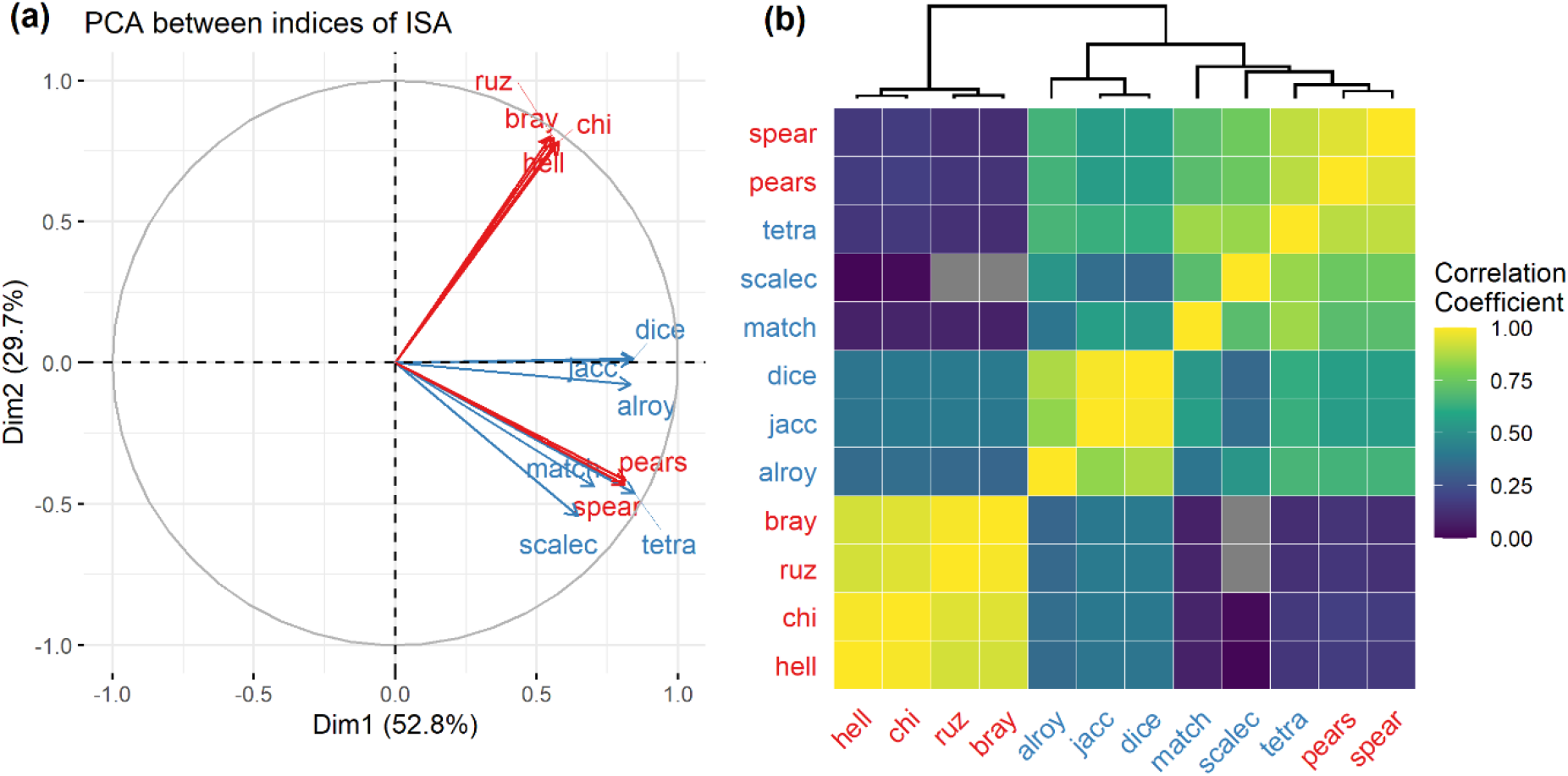
Comparison of interspecific spatial association values among 12 indices, which were subjected to a PCA. (a) A variable plot showing the first two PC axes and loadings of the 12 indices. (b) A correlation plot with a dendrogram showing the topological relationships between indices. For simplicity and ease of visualisation, correlation values were coloured from 0 to 1 only. Continuous indices were in red and binary indices were in blue.

We also observed comparatively higher correlation between pairs of indices with related mathematical properties, even within already tightly formed groups (Fig. 2b). For example, within the first group, the two difference-based indices were highly correlated, as were the two distance-based indices. Combined with the tight grouping of continuous and binary correlation-based indices (pears, spear, tetra) (Fig. 2a, 2b), our results indicate the underlying mathematical property of the index, or its interpretation of association, as the main factor driving differences in measurements of association.

### Variance among clustering algorithms and dendrograms

Although it was difficult to separate out variances in dendrogram outcomes due to the choice of clustering algorithm rather than association index, we observed some general trends. The most striking trend was the presence of reversals, or the upward branching of nodes, among dendrograms resulting from clustering algorithms UPGMC and WPGMC (Supporting Information Fig S4). Reversals greatly hindered the interpretation of hierarchical relationships and delineation of discrete clusters, often also resulting in statistically incomprehensible dendrogram structures (Wedley et al. 1993, Miyamoto 2012, Abe et al. 2017). Hence, we rejected clustering outcomes resulting from the UPGMC and WPGMC algorithms, regardless of their dendrogram performance (for variable plots with UPGMC and WPGMC, see Supporting Information Fig. S5).

Among the remaining five clustering algorithms, resultant variances in dendrogram structure depended on the underlying association index (for variable plots of all 12 association indices, see Supporting Information Fig. S5). Dendrograms based on Bray-Curtis dissimilarity were generally less varied, as seen through closely grouped vectors (small between arrow angles) and high loading scores (long arrow lengths) in the PCA variable plot (Fig. 3a). Comparatively, dendrograms based on Jaccard index were more varied; their vectors were more dispersed (Fig. 3b). Although vectors representing dendrograms based on Spearman correlation also grouped together, their loading scores were lower than those based on Bray-Curtis dissimilarity, which indicated weaker correlations and higher variances in dendrogram structure (Fig. 3c; for full correlation plots, see Supporting Information Fig. S9).

**Figure 3.**
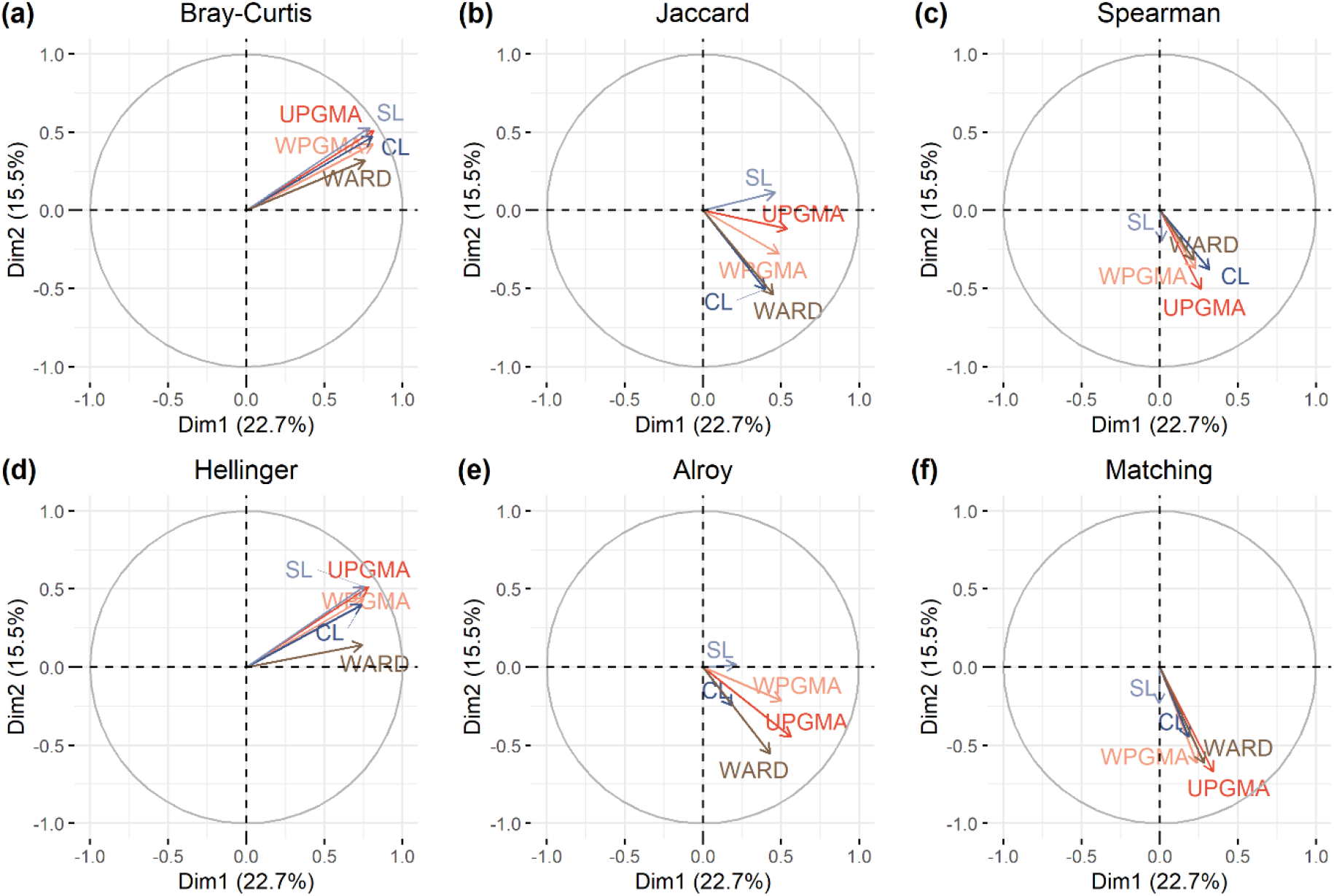
Comparison of dendrograms through a PCA (applied to all 84 candidate dendrograms). For clarity, results were separated and plotted for a subset of indices: (a) Bray-Curtis, (b) Jaccard, (c) Spearman, (d) Hellinger, (e) Alroy, and (f) Matching. Each variable plot showed the first two PC axes and loadings of dendrograms resulting from five clustering algorithms (excluding UPGMC and WPGMC because they led to reversals), which were based on a particular association index. Note, PC axes and loadings across panels were comparable as they were obtained from the same PCA.

Dendrogram outcomes indicated groups of association indices for which the resulting dissimilarity matrices exhibited low or high sensitivity to the choice of clustering algorithm. For example, variations among dendrograms based on Jaccard index and Alroy coefficient were moderately high, even among dendrograms resulting from the same clustering algorithm (Fig. 3b, 3e). By comparison, dendrograms based on Bray-Curtis dissimilarity and Hellinger distance were generally less varied (Fig. 3a, 3c). Dendrograms based on Spearman correlation and Matching coefficient were also quite similar to each other (Fig. 3c, 3f). Note that because we compared dendrograms using Baker’s Gamma, variance here pertained specifically to differences in dendrogram structure as defined by the relative positioning of their nodes (for comparisons using co-phenetic correlation, common nodes, or FM-index, see Supporting Information Fig. S6-S8 and S10-S12).

### Evaluations of dendrogram performance

Dendrogram performance varied substantially across association indices and clustering algorithms (Fig. 4; for performances of all 84 candidate dendrograms, see Supporting Information Fig. S13). We first examined performances among clustering algorithms. Dendrograms most faithful to the original dissimilarity matrix (i.e., co-phenetic correlation) were generally those resulting from UPGMA, while the least faithful resulted from CL, SL, and WARD (Fig. 4a). Cluster strength (i.e., agglomerative coefficient) and balance (i.e., tree balance) were clearly highest for WARD and second highest for CL, but lowest for SL. Hence, Euclidean scores were generally higher among dendrograms resulting from WARD or CL because they performed better on two out of the three metrics (Fig. 4b). Next, among dendrograms based on different association indices, dendrograms based on difference- and distance-based indices (bray, hell) had co-phenetic correlation scores that were clearly higher (Fig. 4a). Agglomerative coefficient scores were typically lower for dendrograms based on binary indices, except Alroy coefficient (Fig. 4a) and Scaled C-score (Supporting Information Fig. S13). However, we did not observe any clear differences in tree balance scores among indices. Hence, Euclidean scores were generally higher among dendrograms based on difference- and distance-based indices (bray, hell) because they performed much better in terms of co-phenetic correlation (Fig. 4b). Overall, dendrograms with the highest Euclidean score were those based on Bray-Curtis dissimilarity and clustered using either the UPGMA, CL, or WARD algorithm (Fig. 4b).

**Figure 4.**
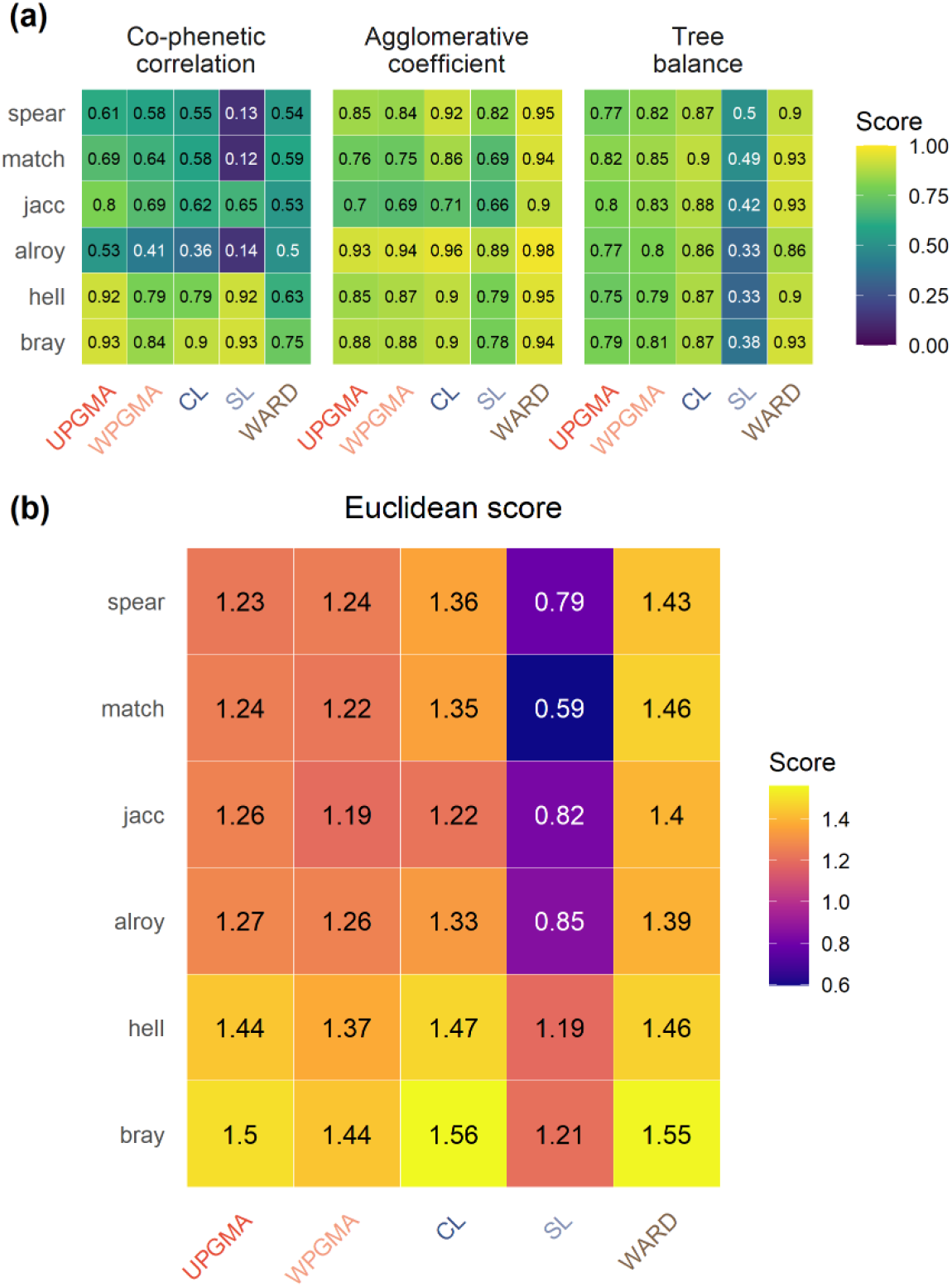
The performance of dendrograms resulting from five clustering algorithms (excluding UPGMC and WPGMC because they led to reversals) for a subset of association indices. (a) Dendrogram performance based on the three metrics: co-phenetic correlation, agglomerative coefficient, and tree balance. (b) Dendrogram overall performance as the Euclidean distance (score) across the three metrics after applying a min-max scaling.

### Evaluations of NMDS plots and dendrograms

The NMDS stress levels were lowest for difference-based indices (bray = 0.103; Fig. 5a–5c), low for distance-based indices (hell = 0.157; Fig. 5d, 5e), but extremely high for correlation-based indices (spear = 0.279; Fig. 5f, 5g). Stress levels were also extremely high for binary indices, except those of proportional overlap (match = 0.23 and jacc = 0.227; Fig. 5h–5k) (Fig. 5). Higher stress levels suggest those indices had captured spatial relationships that were too complex to accurate represent in low-dimensional space. Thus, separations among clusters in highly stressed NMDS space may not be visually apparent, while visually overlapping clusters may be artefacts of imperfectly reduced axes.

**Figure 5.**
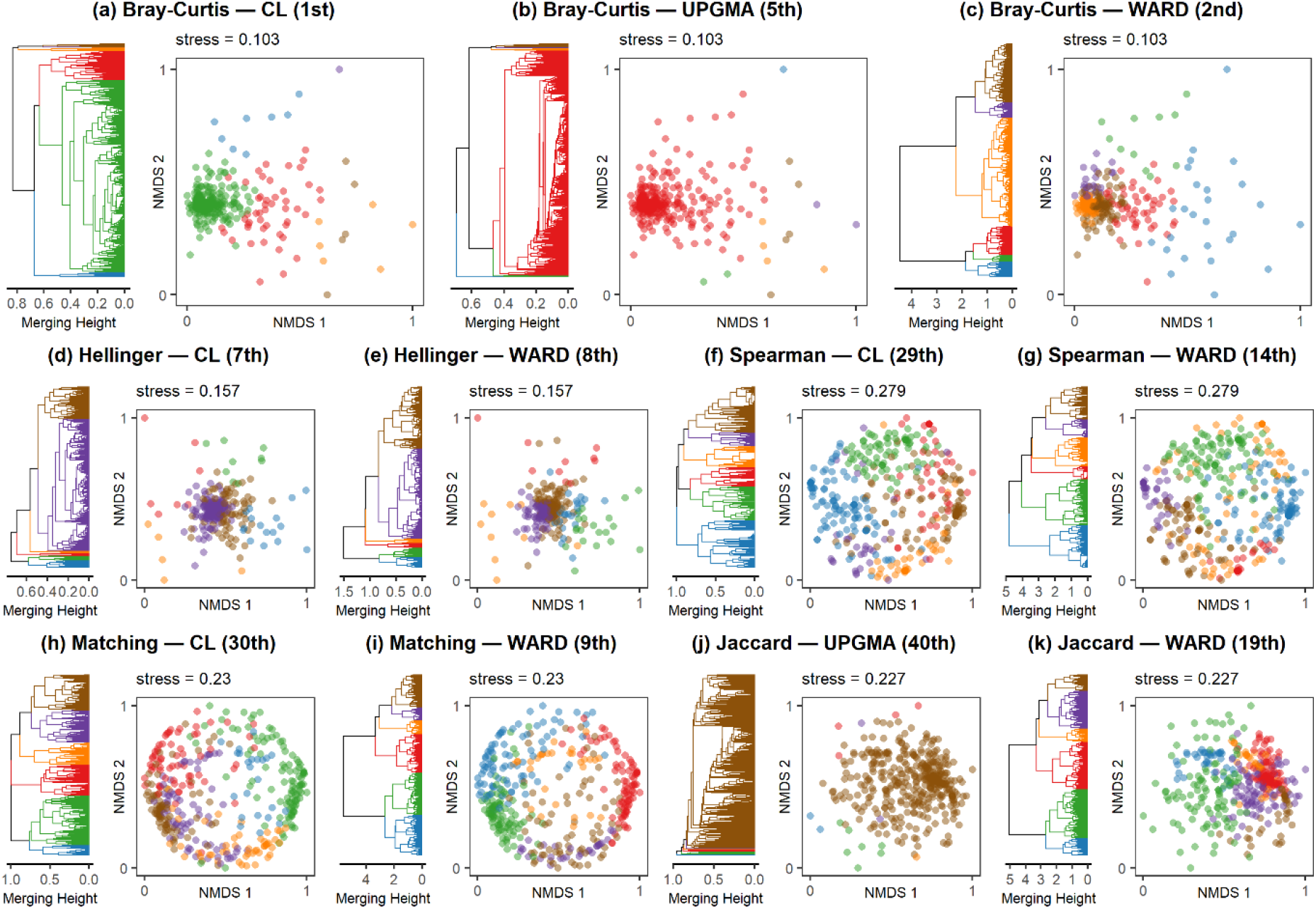
The dendrograms of 11 clustering outcomes and their underlying dissimilarity matrix visualised through an NMDS plot. Stress values were indicated above the NMDS plot. Each leaf of the dendrogram and each dot in the scatter plot represents a species’ distribution. For comparisons across clustering outcomes, the number of clusters was fixed at *k* = 6 and differentiated using colours. Colours were neither comparable nor relevant between panels. Panel titles indicated the association index and clustering algorithm used and its overall rank, i.e., association index — clustering algorithm (overall rank). Dendrograms and dissimilarity matrices were based on association indices Bray-Curtis dissimilarity (a, b, c), Hellinger distance (d, e), Spearman correlation (f, g), Matching coefficient (h, i), and Jaccard index (j, k), resulting from clustering algorithms CL (a, d, f, h), UPGMA (b, j), and WARD (c, e, g, I, k).

The NMDS plots revealed distinct differences in data structure that depended on the choice of association index. Difference- and distance-based indices resulted in points that formed a central aggregate with a scattering of outliers; the former’s outliers were more unidirectional (Fig. 5a–5c), and the latter’s more bidirectional (Fig. 5d, 5e). Comparatively, other indices resulted in points that were spread more evenly (Fig. 5f–5i). To a lesser extent, binary indices that exclude co-absences also resulted in a central aggregate of points (jacc, dice, alroy) (Fig. 5j, 5k).

Cluster memberships showed how the spread of points affected clustering outcomes under different clustering algorithms. Clustering algorithms sensitive to outliers, such as UPGMA, and CL to some extent, tended to classify the central aggregation of points as one large cluster and outliers as multiple smaller clusters. We observed this for difference- and distance-based indices (Fig. 5a, 5b, 5d), and binary indices that exclude co-absences (Fig. 5j). Although accompanying dendrograms indicated that the central cluster could be partitioned at higher values of *k* (number of clusters), it would also result in the excessive partitioning of outlier clusters and higher instances of one-species clusters. Comparatively, WARD was less sensitive to outliers in general, wherein the central aggregation of points was partitioned while outliers formed moderately sized clusters (Fig. 5c, 5e, 5k). As a result, WARD produced cluster memberships that were more balanced, and thus more meaningful for subsequent spatial analyses. Problems with unbalanced cluster memberships were less pertinent when the association index used resulted in evenly spread points (Fig., 5f–5i).

### Final clusters of spatially associated species

As the final clustering outcome, we selected clusters based on Bray-Curtis dissimilarity resulting from the WARD clustering algorithm, which had the second-best Euclidean score (Fig. 4b). We selected the second-best rather than best clustering outcome because it produced clusters that were more balanced, and thus more meaningful for subsequent analyses (Fig. 5a, 5c). Moreover, its Euclidean score was only 0.01 lower than the best score (Fig. 4b).

The optimal number of clusters *k* varied across stopping rules (Supporting Information Table S2). Among diagnostic graphs, the L-method identified *k* = 4 for merging height and *k* = 6 for within-cluster variance, regardless of the cluster centre used to quantify within-cluster variance. For between-cluster variance, the L-method identified *k* = 5 when aggregated distributions (centroids) were used as cluster centres and *k* = 4 when indicator distributions (medoids) were used instead. However, all diagnostic graphs showed a relatively smooth curvature (Supporting Information Fig. S2), suggesting the optimal number of clusters *k* to be above 6 as the L-method tends to underestimate *k* in such cases (Salvador and Chan 2004). The bifurcation paired t-test identified *k* = 34 and 11, for cluster centres using aggregated and indicator distributions, respectively. The bifurcation paired t-test tended to identify a large *k* when aggregated distributions were used, particularly for clusters resulting from WARD, but identified *k* closer to the other three stopping rules when indicator distributions were used instead (Supporting Information Table S2). This was likely because medoids typically represent image type datasets (i.e. raster maps) better than centroids and are less sensitive to outliers that might inflate changes in within-cluster variance (Van der Laan et al. 2003, Kaufman and Rousseeuw 2005). Hence, we set *k* as 11, as identified by the bifurcation paired t-test when using indicator distributions (medoids). Although only one value of *k* was selected, we acknowledged that a range of possible *k* values exists and explored other probable values of *k* in Supporting Information Fig. S14-S16 (Gauch and Whittaker 1981).

### Clusters of tree species distributions in Borneo

Resultant clusters and their representative distributions (i.e., proportion of member species present) yielded spatially meaningful patterns of tree species distributions in Borneo (Fig. 6a) (for aggregate and indicator species distributions, i.e., cluster centroids and medoids, see Supporting Information Fig. S17 and S18). Representative distributions delineated, to some extent, the geographical unit to which member species were endemic, and gradients indicate site suitability for supporting member species. The environmental conditions underlying each representative distribution were also extracted to characterise their habitats (Supporting Information Fig. S19). Interestingly, many of the representative distributions here were also observed in other well-performing dendrograms (Supporting Information Fig. S20), even when their dendrogram structure or cluster memberships differed greatly. This suggests that despite relatively varied clustering outcomes, well-performing dendrograms tended to identify clusters reflecting similar spatial patterns.

**Figure 6.**
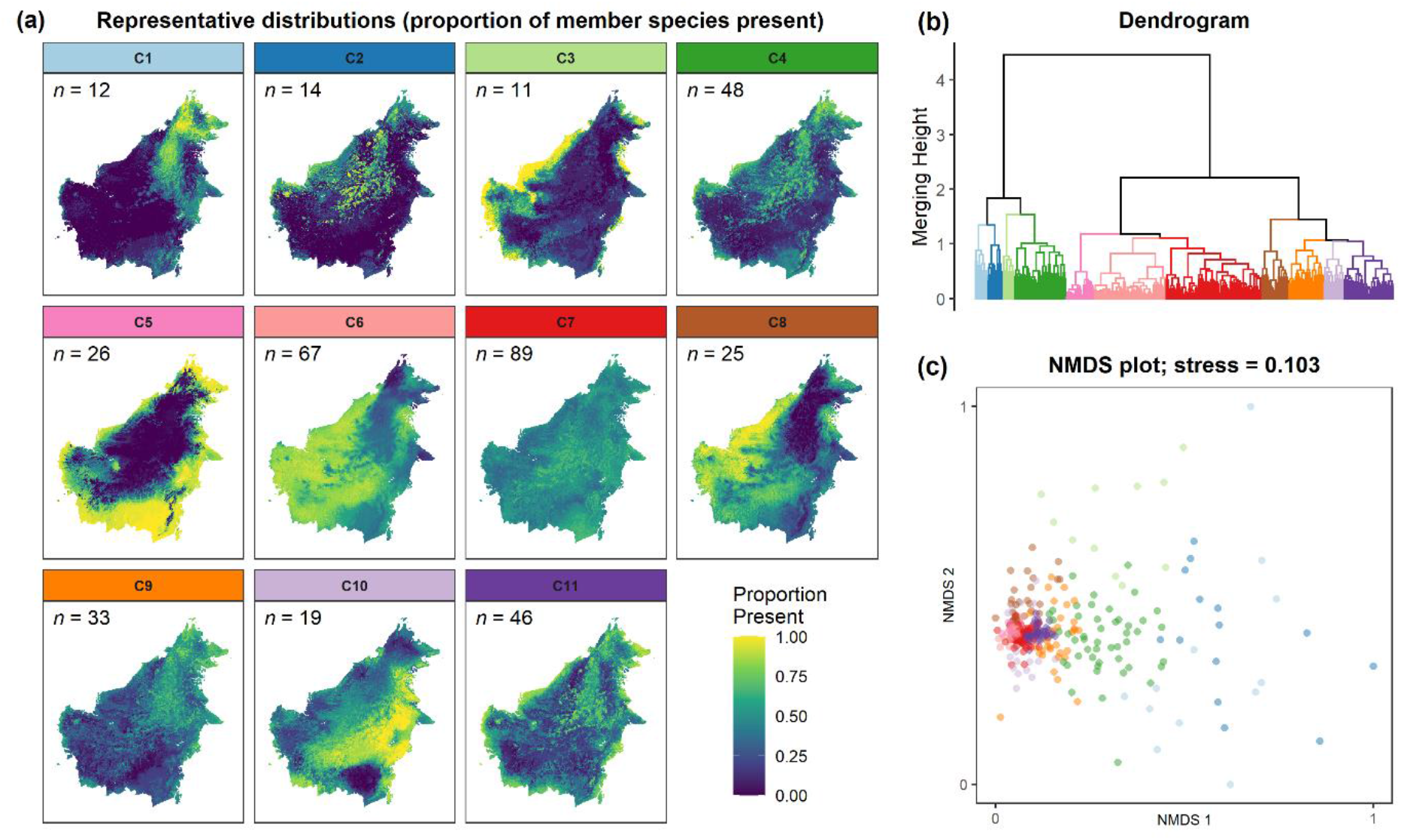
The visualisation of clusters obtained from the second-best performing clustering outcome (Bray-Curtis — WARD) for number of clusters *k* = 11. (a) The representative distribution of each cluster, where *n* equals the number of member species. Representative distributions were generated by summing the binary data of member species at each pixel and dividing values by *n* (i.e., proportion of member species present). (b) The dendrogram of the clustering outcome and (c) its underlying dissimilarity matrix visualised as an NMDS plot. Cluster memberships here (*k* = 11) differed from that in Figure 5c (*k* = 6). Clusters were differentiated by colour, which were consistent across panels and for Figures 6 and 7.

The first split among species distributions was between clusters 1 to 4 and clusters 5 to 11, and occurred early (i.e., high merging height in Fig. 6b), indicating high dissimilarity between partitions. This first split separated clusters with highly range-restricted distributions (clusters 1 – 4) from the rest (clusters 5 – 11) (Fig. 6a, 6c). Cluster 3 was distributed predominantly across Borneo’s western coastal/peatland regions, and clusters 1, 2, and 4 were restricted to separate parts of Borneo’s central montane region. Many peatland species are known, and were found here, to occur in montane habitats (e.g., *Litsea accedens* and *Timonius flavescens;* Slik 2009), which may explain the grouping of cluster 3 with clusters 1, 2, and 4 (but see Discussion: Methods implication for ecological interpretations).

The second split was between clusters 5 to 7 and clusters 8 to 11 (Fig. 6b, 6c). Among clusters 8 to 11, cluster 8 was the most distinct since that cluster split off relatively early in the dendrogram, and was distributed mainly across the western lowlands, like cluster 3, but more inland than coastal (Fig. 6a). Comparatively, clusters 9 to 11 split up much later and at near identical levels (merging height = 1.04 and 1.07; Fig. 6b), indicating low and even measures of between-cluster dissimilarities. While cluster 10 was distributed predominantly along the eastern lowland regions, clusters 9 and 11 were distributed across the mid-montane regions but over areas with vastly different underlying soil conditions (mainly available water and cation exchange capacity; Supporting Information Fig. S19).

Lastly, the remaining clusters, 5 to 7, split up late and at near identical levels (merging height = 1.11 and 1.18; Fig. 6b). Cluster 5 characterised the coastal/peatland forests of Indonesian Borneo (south and east Kalimantan), occupying areas south and east of Borneo’s central mountain range (Fig. 6a). Cluster 6 was broadly distributed across the lowland regions west and south of Borneo’s central mountain range. Cluster 7 contained the most species (*n* = 89) and had a generally widespread distribution, though with slightly higher proportion values along the mid-montane and south-eastern lowland regions of Borneo.

### Habitat loss due to land-cover changes

We found a substantial loss of habitat due to land-cover changes for all clusters (Fig. 7). By 1992, habitat loss among clusters averaged 30% (Fig. 7a)—highest for clusters 3 and 5 (38% and 44%, respectively) and lowest for cluster 2 (22%). Subsequent land-cover changes resulted in a cumulative mean habitat loss of 43% by 2020—again, habitat loss was highest for clusters 3 and 5 (56% and 61%, respectively) and lowest for cluster 2 (33%) (Fig. 7a).

**Figure 7.**
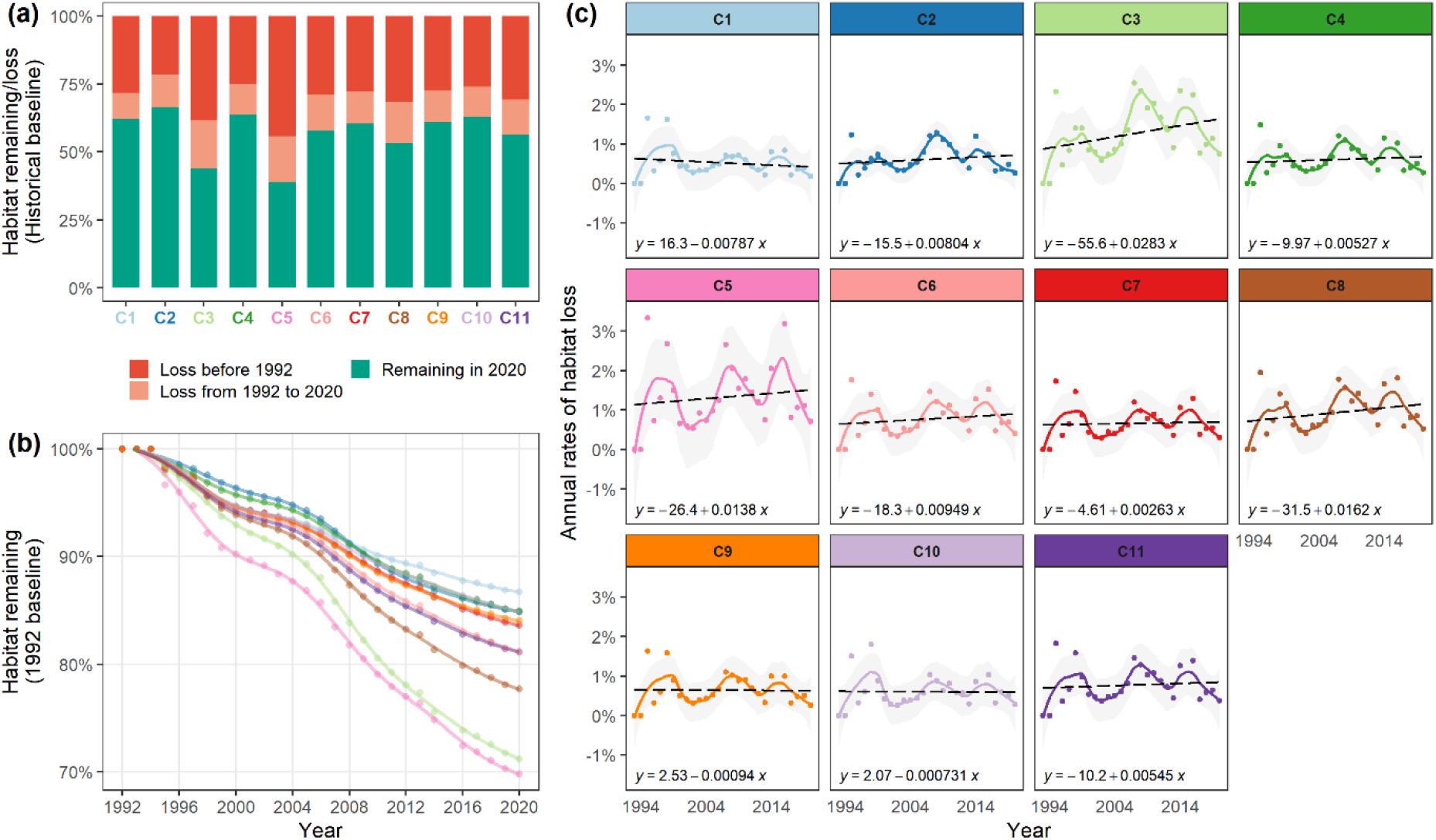
The impact of deforestation on each cluster’s representative distribution as changes in habitat availability. (a) Bar plots showing the percentage of historically available habitats lost before 1992, loss between 1992 to 2020, and remaining in the year 2020. (b) Annual percentages of 1992 habitats remaining from 1992 to 2020 fitted using a generalised additive model. (c) Annual rates of habitat loss from 1993 to 2020 fitted using a local polynomial regression with α = 0.75 (solid, coloured) and a linear regression (dashed, black). Clusters are differentiated by colour and are consistent across panels and for Figures 6 and 7.

Annual trends made apparent the differences in habitat loss among clusters. Most striking was the severe and continued loss of coastal/peatland habitats supporting clusters 3 and 5, and the western lowland habitats supporting cluster 8 (Fig. 7b). Rates of habitat loss from 1992 to 2020 increased for most clusters and was also highest for clusters 3, 5, and 8; only clusters 1, 9, and 10 experienced a decrease in rates of habitat loss (Fig. 7c). Annual trends also revealed cycles in habitat loss, oscillating from a slump to a peak rate of habitat loss (Fig. 7b, 7c). The first cycle started from a slump in 1994 to a peak in 1998, the second from 2003 to 2007, and the third from 2013 to 2016.

Differing rates of habitat loss between cycles indicates potential shifts in habitats targeted for land-cover change. Among montane distributed clusters 1, 2, and 4 (Fig. 6a), cluster 1 experienced higher rates of habitat loss than those of clusters 2 and 4 in the first cycle (1994 – 1998) (Fig. 7c). The subsequent cycle saw a switch, with rates of habitat loss being greater for clusters 2 and 4 instead (2002 – 2007). Although comparatively lower in the third cycle, rates of habitat loss were still higher for clusters 2 and 4. This was likely a recent shift, as prior to 1992, clusters 2 and 4 were the clusters least impacted by land-cover changes (Fig. 7a). Between coastal-/peatland-distributed clusters 3 and 5, rates of habitat loss surged for cluster 3 in the second cycle (2002 – 2007) but there was also an increase for cluster 5, albeit an increase of lower magnitude than for cluster 3. Moreover, the increased rate of habitat loss for cluster 3 in the second cycle was so great that its proportion of habitat remaining in 1992 that was lost by 2012 nearly matched that of cluster 5 (Fig. 7b). Although we did not observe a similar surge in rates of habitat loss for cluster 3 in the third cycle (2012 – 2016), its overall increase from 1992 to 2020 was the highest among clusters (Fig. 7c).

## Discussion

To capitalise on the emerging wealth of distribution data, researchers must first contend with a corresponding increase in data complexity. We demonstrate how the clustering of spatially associated species offers a way to simplify that complexity to unravel meaningful patterns in species distributions, where resultant clusters provide a spatially explicit framework for investigating distribution-related questions in ecology, biogeography, and conservation (Clements 1936, Marquet et al. 2004, Legendre and Legendre 2012, HilleRisLambers et al. 2012). Importantly, clusters are quantitatively derived and easily reproducible, while the methodological framework promotes transparency and peer-scrutiny of the methods and results. The primary advantage of the framework, however, is that it makes key steps of the clustering process more explicit, forcing practitioners to carefully consider their methodological choices in relation to their research objectives (Legendre and Legendre 2012). Combined with steps that support practitioners in investigating different methods and interpreting results, our methodological framework encourages more informed decision-making and rigorous selection of final clustering outcomes.

### Clusters of spatially associated species with variable trends of habitat loss

Our methods identified 11 distinct clusters of tree species in Borneo, based on their spatial distributions, which provided valuable ecological and conservation insights. Montane species clusters 1, 2, and 4 had the most distinct distributions because their ranges were highly restricted. Their distributions also matched previous predictions of endemicity by Raes et al. (2009): in particular, high endemicity in the northern montane region of Borneo coincided with an area of overlap among our three montane—and some sub-montane—clusters. With narrow ranges and lack of areas to migrate upwards in elevation under climate change (Bellard et al. 2014, Yanahan and Moore 2019, Pang et al. 2021), these montane-distributed clusters are of great conservation value (Villalobos et al. 2013, Guisan et al. 2013, Struebig et al. 2015). We also observed clusters restricted to western Borneo: the coastal distributed cluster 3, slightly more inland cluster 8, and more widespread cluster 6 that extended further south as well. Because our SDMs relied on environmental variables, this restriction stems from a unique set of environmental conditions (i.e., high precipitation and low clay content; Supporting Information Fig. S19). Indeed, western Borneo is home to many endemic and is noted for its unique floral composition, with present day environmental conditions previously also identified as probable drivers (Slik et al. 2003, 2011, Neo et al. 2021). Conversely, the more widely distributed clusters 5, 7, 9, and 10 seem to represent eastern Borneo, which is characterised by a combination of low precipitation, low available water capacity, and high soil clay content (Supporting Information Fig. S19).

Our general results of severe habitat loss corroborated previous assessments of deforestation in Borneo (Miettinen et al. 2011, Gaveau et al. 2014, 2016, 2019, Sloan et al. 2019, Wong et al. 2020), but our classification of species distributions allowed us to separate variable trends of habitat loss to discern especially threatened species groups. Clusters with coastal/peatland distributions (clusters 3 and 5) suffered the most severe loss of habitat, likely due to extensive oil palm expansions that primarily target coastal/peatland habitats (Miettinen et al. 2011, Gaveau et al. 2014, 2016).

However, the western-restricted, inland cluster 8 also suffered severe habitat losses, losses not observed for its eastern-restricted, inland counterpart, cluster 10. Water stress greatly limits oil palm yields, where greater rainfall in western than southern/eastern Borneo may be facilitating oil palm plantation expansions into the more inland habitats (Carr 2011, Sa’adi et al. 2021; Supporting Information Fig. S19). Although deforestation—in absolute terms—is less extensive in western Borneo (Miettinen et al. 2011, Gaveau et al. 2014, 2019), cluster 8 has a narrower distribution than most, resulting in it suffering the third most severe loss of habitat.

Similarly, because our method separated montane habitats into three distinct clusters, we were able to identify a switch in habitat loss severity. The northern montane-distributed cluster 1 suffered habitat losses much greater in the first cycle (peak in 1998) than those after (peaks in 2007 and 2016); likely due to protected area implementation and enforcement in the Kinabalu montane alpine meadows ecoregion that coincides most with cluster 1’s distribution (Olson et al. 2001, Phua et al. 2008). However, this protection seemed to not extend to the more central, western montane-distributed clusters 2 and 4, which suffered greater habitat losses in later cycles. Our results highlight the emerging threat of land-cover changes for tropical montane habitat that recent studies have also confirmed (Karger et al. 2021, Feng et al. 2021), but that there exists variability in those trends of loss. Our separation of montane species clusters and their variable loss of habitat can facilitate more targeted conservation planning to better protect those more threatened in recent decades (Ashcroft 2010, Guisan et al. 2013, Struebig et al. 2015).

The observed temporal oscillations in habitat losses were also of interest. Oscillations were temporally congruent across clusters, with peaks in 1995, 1998, 2007, and 2016 that coincided with notable El Niño periods of 1992-1995, 1997-1998, 2005-2007, and 2014-2016 (NOAA 2022). Moreover, studies have found El Niño effects to compound with deforestation to increase forest fire severity and frequency (Wooster et al. 2012, Huijnen et al. 2016, Sloan et al. 2019, Chapman et al. 2020; but see Langner and Siegert 2009, Gaveau et al. 2015), which may explain differing magnitudes of oscillation among clusters. Because a substantial proportion of habitat losses was tied to these oscillations, its exact driver warrants further investigation, for which our clusters provide the framework to.

### Further applications

The application of the clustering of spatially associated species goes beyond separating trends of habitat loss. We may use clustering outcomes to investigate structural changes in species associations. For instance, due to the spread of an invasive species or climate-induced range shifts, which is a particularly underappreciated facet of global change impacts on biodiversity (Early and Sax 2014, Krosby et al. 2015, Keil et al. 2021). Alternatively, we might compare the resultant dendrogram against a phylogenetic tree to investigate speciation events linked to present-day spatial patterns (Villalobos et al. 2017), or against dendrograms of functional similarity to uncover co-existence or competitive mechanisms that underlie co-occurrence patterns (Rüger et al. 2020).

The representative spatial distribution of species from each group is also useful for biodiversity monitoring and management (Webb 1989, Cousins 1991). Commonly used criteria for evaluating the conservation value of sites, like absolute species richness or beta-diversity, are prone to taxonomic and sampling biases (i.e., the so-called Linnean and Wallacean shortfalls) (Lomolino 2004, Whittaker et al. 2005, Possingham et al. 2007), which typically overrepresent widespread and easy-to-detect species (Prendergast 1993, Jetz and Rahbek 2002, Lennon et al. 2003, Boakes et al. 2010). However, in using species clusters and their representative distributions instead, site evaluations are unbiased by the relative number of species from each cluster (Roberge and Angelstam 2004, Possingham et al. 2007). Widespread species will also form their own groups and not affect evaluations tied to range-restricted species. Moreover, representative distributions indicate the geographical unit to which member species reside and are, to some extent, endemic. Representative distributions are thus highly informative when the goal is to detect specific ecosystems, protect species ranges, and assess extinction risks (Guisan et al. 2013, Hannah et al. 2020).

### On selecting association indices and clustering algorithms

Our findings highlight the value of our methodological framework, not just its individual steps but also the testing of multiple methods. Different association indices and clustering algorithms led to clustering outcomes that were highly varied in dendrogram structure, performance, and cluster memberships. Data-driven assessments and comparisons and ecological theory need to guide the selection of an appropriate method.

Exploring multiple association indices is especially crucial given the sequential nature of steps in our framework, in that the index influences how clustering algorithms work and perform (Jain et al. 1999, Legendre and Legendre 2012). In our study, we explored a wide range of binary and continuous indices with different inherent mathematical properties. Like previous studies, indices varied greatly in their measurement of associations but generally separated into three groups as determined by those mathematical properties: (1) difference- and distance-based indices; (2) binary indices that exclude co-absences; and (3) correlation-based indices and binary indices that include co-absences (Hubálek 1982, Keil et al. 2021). Given the computational intensity of quantifying associations across entire spatial extents (maps in lieu of plots), we advocate exploring at least one index from each group. Beyond the groups described, practitioners may consider other qualities when selecting (or excluding) indices to explore: ability to recover simulated magnitudes of spatial attraction and repulsion, into which Keil et al. (2021) offers insight; adherence to the triangle inequality rule (e.g., Hellinger distance), an important axiom of distance matrices for geometric clustering; non-parametric to allow measurements between non-comparable data without any prior transformation (e.g., Spearman correlation for occurrence probabilities between SDM algorithms, or count data between species with disparate raw abundances) (Warren et al. 2008, 2019); or popularity and simplicity to facilitate the interpretation of results (e.g., Jaccard index as proportional overlap).

Our framework emphasises testing multiple clustering algorithms because algorithms vary in their capacity to simultaneously minimise within-cluster and maximise between-cluster variances. This was observed in the variation across dendrogram performances, which offer great insight into selecting clustering algorithms with certain characteristics (Sokal and Rohlf 1962, Rousseeuw 1985, Legendre and Legendre 2012, Fernández and Gómez 2020). The relative importance of those characteristics, represented by each dendrogram performance metric, depends on the research objective. When examining changes in a community’s spatial structure over time (e.g., due to invasive species introduction or climate-induced range shifts; Early and Sax 2014, Krosby et al. 2015), it is important to minimise data distortions introduced by clustering. Thus, faithfulness to the original dissimilarity matrix may be the only priority, in which case UPGMA might be preferred. Alternatively, if the main objective is to decompose a large dataset into smaller but evenly sized subsets, cluster strength and balance may be more important, in which case WARD and CL might be selected instead. The flexibility of this step goes beyond our three metrics, where practitioners may incorporate others of importance, such as space distortion ratio, connectedness, or isolation (see

Estabrook 1966, Wirth et al. 1966, Legendre and Legendre 2012, Fernández and Gómez 2020). Practitioners must also consider the relevance of each candidate algorithm; selected algorithms should have linkage functions that align with the study’s theoretical expectation of how clusters form (see Erman et al. 2015, Seif 2018, Roux 2018). Here, we rejected dendrograms resulting from UPGMC and WPGMC to avoid reversals (Wedley et al. 1993, Miyamoto 2012, Abe et al. 2017).

Although for the case study of Borneo, we selected the dendrogram based on Bray-Curtis dissimilarity resulting from WARD because its performed (second) best, this does not suggest that practitioners should always base their selection on dendrogram performance scores. For example, Spearman correlation captured associations that were inherently more complex (i.e., highly stressed 2-dimension NMDS), and thus difficult to maintain during the clustering analysis (i.e., low co-phenetic correlation scores). However, this complexity may be a point of ecological interest rather than a basis for rejection, which reemphasises the need for ecological theory to guide comparisons of multiple indices. The resultant data structure of a chosen association index also determines if the clustering algorithm achieves meaningful clusters (Jain et al. 1999, Rajalingam and Ranjini 2011, Legendre and Legendre 2012, Roux 2018). In cases where it does not, practitioners might select an alternative—but comparably well-performing—algorithm instead, as we did for our case study. Indeed, dendrogram scores provide crucial information on each clustering outcome, but data visualising tools like NMDS are equally vital aids for selecting methods most appropriate to one’s study objective.

### Implication of methods for interpretating species clusters

Despite substantial variations among candidate clustering outcomes, broad patterns in tree species distributions remained relatively consistent as many of those observed in our chosen clustering outcome were observed among other well-performing clustering outcomes as well (i.e., high dendrogram performance and meaningful cluster sizes). This consistency lends credibility to the spatial patterns of our chosen clustering outcome and the robustness of employing species clusters. However, distributional patterns characterising widespread species were irregular, suggesting the classification of those species to be partly contingent on the association index and clustering algorithm used (Jongman et al. 1995, Dufrêne and Legendre 1997, Legendre and Legendre 2012). Although most clustering outcomes were still able to discern some general patterns in widespread species distributions, our findings highlight that special attention is needed when clustering widely distributed species; more so because such clusters may represent hyper-abundant species that comprise the bulk of stem density and aboveground carbon (Fauset et al. 2015), where an understanding of their spatial patterns may improve the protection of high-carbon forests (Sullivan et al. 2017, 2020, Siman et al. 2021).

When interpreting clusters, we must also recognise the limits of the underlying methods and how that might affect ecological interpretations. In selecting Bray-Curtis dissimilarity, we define associations as absolute differences in occurrence probability. Hence, dissimilarities were low between species with probabilities close in value, even when their binary distributions were non-overlapping or when their probabilities were uncorrelated (see clusters 10 and 11 representative distributions versus their relatedness) (Fig. 6). Because many lowland species exhibited this characteristic, we found many species with relatively low dissimilarities (large aggregate of points in the NMDS plot), which did not apply to other groups of associations indices (e.g., correlation-based indices). The inverse was also true for species with highly skewed probabilities (i.e., high dissimilarities due to sharp probability gradients), which was probably why highly range-restricted species were predominantly further dispersed in the NMDS plot. Thus, we acknowledge that dissimilarities among range-restricted species and their clusters might have been exaggerated to a degree, while more nuanced patterns among lowland widespread distributions might have been overlooked.

In selecting the WARD clustering algorithm, resultant cluster boundaries are typically oddly shaped and evenly sized (i.e., number of objects per cluster) (Ward 1963, Legendre and Legendre 2012, Erman et al. 2015, Seif 2018). As a result, WARD was able to partition the large aggregate of points and form meaningfully clusters. The trade-off, however, is that WARD often distorts the dissimilarity space, especially for ecological data where objects often exist along a continuum (Kreft and Jetz 2010, Legendre and Legendre 2012, Holt et al. 2013, Fernández and Gómez 2020). Hence, resultant hierarchical relationships were likely inaccurate in representing original dissimilarities and must be carefully interpreted. In summary, we emphasise that the most appropriate clustering outcome is context- and need-dependent. This need for a flexible combination of ecological theory and data-driven assessments—to guide practitioners in considering the benefits and drawbacks of each method—is embedded in our framework.

## References

Abe, R., S. Miyamoto, Y. Endo, and Y. Hamasuna. 2017. Hierarchical clustering algorithms with automatic estimation of the number of clusters. Pages 1–5 2017 Joint 17th World Congress of International Fuzzy Systems Association and 9th International Conference on Soft Computing and Intelligent Systems (IFSA-SCIS).

Aiello-Lammens, M. E., R. A. Boria, A. Radosavljevic, B. Vilela, and R. P. Anderson. 2015. spThin: an R package for spatial thinning of species occurrence records for use in ecological niche models. Ecography 38:541–545.

Allen, T. F. H., and T. W. Hoekstra. 1990. The confusion between scale-defined levels and conventional levels of organization in ecology. Journal of Vegetation Science 1:5–12.

Ashcroft, M. B. 2010. Identifying refugia from climate change. Journal of Biogeography:1407–1413.

Baatar, U.-O. 2019. Evaluating climatic threats to habitat types based on co-occurrence patterns of characteristic species. Basic and Applied Ecology 38:13.

Baker, F. B. 1974. Stability of two hierarchical grouping techniques case I: sensitivity to data errors. Journal of the American Statistical Association 69:440–445.

Beech, E., M. Rivers, S. Oldfield, and P. P. Smith. 2017. GlobalTreeSearch: The first complete global database of tree species and country distributions. Journal of Sustainable Forestry 36:454–489.

Bellard, C., C. Leclerc, B. Leroy, M. Bakkenes, S. Veloz, W. Thuiller, and F. Courchamp. 2014. Vulnerability of biodiversity hotspots to global change. Global Ecology and Biogeography 23:1376–1386.

Blanchet, F. G., K. Cazelles, and D. Gravel. 2020. Co-occurrence is not evidence of ecological interactions. Ecology Letters 23:1050–1063.

Boakes, E. H., P. J. K. McGowan, R. A. Fuller, D. Chang-qing, N. E. Clark, K. O’Connor, and G. M. Mace. 2010. Distorted Views of Biodiversity: Spatial and Temporal Bias in Species Occurrence Data. PLoS Biology 8:e1000385.

Boria, R. A., L. E. Olson, S. M. Goodman, and R. P. Anderson. 2017. A single-algorithm ensemble approach to estimating suitability and uncertainty: cross-time projections for four Malagasy tenrecs. Diversity and Distributions 23:196–208.

Boyce, P. C., and S. Y. Wong. 2019. Borneo and its disproportionately large rheophytic aroid flora. Gardens’ Bulletin Singapore 71:497–524.

Calatayud, J., E. Andivia, A. Escudero, C. J. Melián, R. Bernardo-Madrid, M. Stoffel, C. Aponte, N. G. Medina, R. Molina-Venegas, X. Arnan, M. Rosvall, M. Neuman, J. A. Noriega, F. Alves-Martins, I. Draper, A. Luzuriaga, J. A. Ballesteros-Cánovas, C. Morales-Molino, P. Ferrandis, A. Herrero, L. Pataro, L. Juen, A. Cea, and J. Madrigal-González. 2020. Positive associations among rare species and their persistence in ecological assemblages. Nature Ecology & Evolution 4:40–45.

Caliński, T., and J. Harabasz. 1974. A dendrite method for cluster analysis. Communications in Statistics-theory and Methods 3:1–27.

Carr, M. K. V. 2011. THE WATER RELATIONS AND IRRIGATION REQUIREMENTS OF OIL PALM (*ELAEIS GUINEENSIS):* A REVIEW. Experimental Agriculture 47:629–652.

Cayuela, L., Í. Granzow-de la Cerda, F. S. Albuquerque, and D. J. Golicher. 2012. Taxonstand: An R package for species names standardisation in vegetation databases. Methods in Ecology and Evolution 3:1078–1083.

Chapman, S., J. Syktus, R. Trancoso, A. Salazar, M. Thatcher, J. E. M. Watson, E. Meijaard, D. Sheil, P. Dargusch, and C. A. McAlpine. 2020. Compounding impact of deforestation on Borneo’s climate during El Niño events. Environmental Research Letters 15:084006.

Chouikhi, H., M. Charrad, and N. Ghazzali. 2015. A comparison study of clustering validity indices. Pages 1–4 2015 Global Summit on Computer & Information Technology (GSCIT). IEEE, Sousse, Tunisia.

Clements, F. E. 1936. Nature and Structure of the Climax. The Journal of Ecology 24:252.

Collins, S. L., S. M. Glenn, and D. W. Roberts. 1993. The hierarchical continuum concept. Journal of Vegetation Science 4:149–156.

Corlett, R. T., and K. W. Tomlinson. 2020. Climate Change and Edaphic Specialists: Irresistible Force Meets Immovable Object? Trends in Ecology & Evolution 35:367–376.

Cousins, S. H. 1991. Species diversity measurement: choosing the right index. Trends in Ecology & Evolution 6:190–192.

Cramér, H. 1924. Remarks on correlation. Scandinavian Actuarial Journal 1924:220–240.

Currie, D. J. 2019. Where Newton might have taken ecology. Global Ecology and Biogeography 28:18–27.

Curtis, J. T. 1959. The vegetation of Wisconsin: an ordination of plant communities. University of Wisconsin Pres.

Di Febbraro, M., L. Sallustio, M. Vizzarri, D. De Rosa, L. De Lisio, A. Loy, B. A. Eichelberger, and M. Marchetti. 2018. Expert-based and correlative models to map habitat quality: Which gives better support to conservation planning? Global Ecology and Conservation 16:e00513.

Duda, R. O., and P. E. Hart. 1973. Pattern classification and scene analysis. Wiley New York.

Dufrêne, M., and P. Legendre. 1997. Species assemblages and indicator species: the need for a flexible asymmetrical approach. Ecological Monographs 67:345–366.

Early, R., and D. F. Sax. 2014. Climatic niche shifts between species’ native and naturalized ranges raise concern for ecological forecasts during invasions and climate change. Global Ecology and Biogeography 23:1356–1365.

Elith, J., and J. R. Leathwick. 2009. Species Distribution Models: Ecological Explanation and Prediction Across Space and Time. Annual Review of Ecology, Evolution, and Systematics 40:677–697.

Elith, J., S. J. Phillips, T. Hastie, M. Dudík, Y. E. Chee, and C. J. Yates. 2011. A statistical explanation of MaxEnt for ecologists. Diversity and Distributions 17:43–57.

Erman, N., A. Korosec, and J. Suklan. 2015. Performance of selected agglomerative hierarchical clustering methods. Innovative Issues and Approaches in Social Sciences 8:180–204.

ESA. 2017. Land Cover CCI Product User Guide Version 2. Tech. Rep.

Estabrook, G. F. 1966. A mathematical model in graph theory for biological classification. Journal of Theoretical Biology 12:297–310.

Faurby, S., and M. B. Araújo. 2018. Anthropogenic range contractions bias species climate change forecasts. Nature Climate Change 8:252–256.

Fauset, S., M. O. Johnson, M. Gloor, T. R. Baker, A. Monteagudo M., R. J. W. Brienen, T. R. Feldpausch, G. Lopez-Gonzalez, Y. Malhi, H. ter Steege, N. C. A. Pitman, C. Baraloto, J. Engel, P. Pétronelli, A. Andrade, J. L. C. Camargo, S. G. W. Laurance, W. F. Laurance, J. Chave, E. Allie, P. N. Vargas, J. W. Terborgh, K. Ruokolainen, M. Silveira, G. A. Aymard C., L. Arroyo, D. Bonal, H. Ramirez-Angulo, A. Araujo-Murakami, D. Neill, B. Hérault, A. Dourdain, A. Torres-Lezama, B. S. Marimon, R. P. Salomão, J. A. Comiskey, M. Réjou-Méchain, M. Toledo, J. C. Licona, A. Alarcón, A. Prieto, A. Rudas, P. J. van der Meer, T. J. Killeen, B.-H. Marimon Junior, L. Poorter, R. G. A. Boot, B. Stergios, E. V. Torre, F. R. C. Costa, C. Levis, J. Schietti, P. Souza, N. Groot, E. Arets, V. C. Moscoso, W. Castro, E. N. H. Coronado, M. Peña-Claros, C. Stahl, J. Barroso, J. Talbot, I. C. G. Vieira, G. van der Heijden, R. Thomas, V. A. Vos, E. C. Almeida, E. Á. Davila, L. E. O. C. Aragão, T. L. Erwin, P. S. Morandi, E. A. de Oliveira, M. B. X. Valadão, R. J. Zagt, P. van der Hout, P. A. Loayza, J. J. Pipoly, O. Wang, M. Alexiades, C. E. Cerón, I. Huamantupa-Chuquimaco, A. Di Fiore, J. Peacock, N. C. P. Camacho, R. K. Umetsu, P. B. de Camargo, R. J. Burnham, R. Herrera, C. A. Quesada, J. Stropp, S. A. Vieira, M. Steininger, C. R. Rodríguez, Z. Restrepo, A. E. Muelbert, S. L. Lewis, G. C. Pickavance, and O. L. Phillips. 2015. Hyperdominance in Amazonian forest carbon cycling. Nature Communications 6:6857.

Feng, X., D. S. Park, C. Walker, A. T. Peterson, C. Merow, and M. Papeş. 2019. A checklist for maximizing reproducibility of ecological niche models. Nature Ecology & Evolution 3:1382–1395.

Feng, Y., A. D. Ziegler, P. R. Elsen, Y. Liu, X. He, D. V. Spracklen, J. Holden, X. Jiang, C. Zheng, and Z. Zeng. 2021. Upward expansion and acceleration of forest clearance in the mountains of Southeast Asia. Nature Sustainability 4:892–899.

Fernández, A., and S. Gómez. 2020. Versatile Linkage: a Family of Space-Conserving Strategies for Agglomerative Hierarchical Clustering. Journal of Classification 37:584–597.

Fithian, W., and T. Hastie. 2013. Finite-sample equivalence in statistical models for presence-only data. The annals of applied statistics 7:1917.

Gaston, K. J. 1996. Species-range-size distributions: patterns, mechanisms and implications. Trends in Ecology & Evolution 11:197–201.

Gaston, K. J. 2003. The structure and dynamics of geographic ranges. Oxford University Press on Demand.

Gauch, H. G., and R. H. Whittaker. 1981. Hierarchical Classification of Community Data. The Journal of Ecology 69:537.

Gaveau, D. L. A., B. Locatelli, M. A. Salim, H. Yaen, P. Pacheco, and D. Sheil. 2019. Rise and fall of forest loss and industrial plantations in Borneo (2000–2017). Conservation Letters 12.

Gaveau, D. L. A., M. A. Salim, K. Hergoualc’h, B. Locatelli, S. Sloan, M. Wooster, M. E. Marlier, E. Molidena, H. Yaen, R. DeFries, L. Verchot, D. Murdiyarso, R. Nasi, P. Holmgren, and D. Sheil. 2015. Major atmospheric emissions from peat fires in Southeast Asia during non-drought years: evidence from the 2013 Sumatran fires. Scientific Reports 4:6112.

Gaveau, D. L. A., D. Sheil, Husnayaen, M. A. Salim, S. Arjasakusuma, M. Ancrenaz, P. Pacheco, and E. Meijaard. 2016. Rapid conversions and avoided deforestation: examining four decades of industrial plantation expansion in Borneo. Scientific Reports 6:32017.

Gaveau, D. L. A., S. Sloan, E. Molidena, H. Yaen, D. Sheil, N. K. Abram, M. Ancrenaz, R. Nasi, M. Quinones, N. Wielaard, and E. Meijaard. 2014. Four Decades of Forest Persistence, Clearance and Logging on Borneo. PLoS ONE 9:e101654.

GBIF. 2022. Global Data Trends.

Graham, C. H., and R. J. Hijmans. 2006. A comparison of methods for mapping species ranges and species richness. Global Ecology and Biogeography 0:060831043455001–???

Guerra, L., V. Robles, C. Bielza, and P. Larrañaga. 2012. A comparison of clustering quality indices using outliers and noise. Intelligent Data Analysis 16:703–715.

Gueta, T., and Y. Carmel. 2016. Quantifying the value of user-level data cleaning for big data: A case study using mammal distribution models. Ecological Informatics 34:139–145.

Guisan, A., and W. Thuiller. 2005. Predicting species distribution: offering more than simple habitat models. Ecology Letters 8:993–1009.

Guisan, A., R. Tingley, J. B. Baumgartner, I. Naujokaitis-Lewis, P. R. Sutcliffe, A. I. T. Tulloch, T. J. Regan, L. Brotons, E. McDonald-Madden, C. Mantyka-Pringle, T. G. Martin, J. R. Rhodes, R. Maggini, S. A. Setterfield, J. Elith, M. W. Schwartz, B. A. Wintle, O. Broennimann, M. Austin, S. Ferrier, M. R. Kearney, H. P. Possingham, and Y. M. Buckley. 2013. Predicting species distributions for conservation decisions. Ecology Letters 16:1424–1435.

Hannah, L., P. R. Roehrdanz, P. A. Marquet, B. J. Enquist, G. Midgley, W. Foden, J. C. Lovett, R. T. Corlett, D. Corcoran, S. H. M. Butchart, B. Boyle, X. Feng, B. Maitner, J. Fajardo, B. J. McGill, C. Merow, N. Morueta-Holme, E. A. Newman, D. S. Park, N. Raes, and J. Svenning. 2020. 30% land conservation and climate action reduces tropical extinction risk by more than 50%. Ecography:ecog.05166.

Hazzi, N. A., J. S. Moreno, C. Ortiz-Movliav, and R. D. Palacio. 2018. Biogeographic regions and events of isolation and diversification of the endemic biota of the tropical Andes. Proceedings of the National Academy of Sciences 115:7985–7990.

Hengl, T., J. Mendes de Jesus, G. B. M. Heuvelink, M. Ruiperez Gonzalez, M. Kilibarda, A. Blagotić, W. Shangguan, M. N. Wright, X. Geng, B. Bauer-Marschallinger, M. A. Guevara, R. Vargas, R. A. MacMillan, N. H. Batjes, J. G. B. Leenaars, E. Ribeiro, I. Wheeler, S. Mantel, and B. Kempen. 2017. SoilGrids250m: Global gridded soil information based on machine learning. PLOS ONE 12:e0169748.

Hijmans, R. J., and J. V. Etten. 2012. Geographic analysis and modeling with raster data. R Package version 2:1–25.

HilleRisLambers, J., P. B. Adler, W. S. Harpole, J. M. Levine, and M. M. Mayfield. 2012. Rethinking Community Assembly through the Lens of Coexistence Theory. Annual Review of Ecology, Evolution, and Systematics 43:227–248.

Hoekstra, T. W., T. F. H. Allen, and C. H. Flather. 1991. Implicit Scaling in Ecological Research. BioScience 41:148–154.

Holt, B. G., J.-P. Lessard, M. K. Borregaard, S. A. Fritz, M. B. Araújo, D. Dimitrov, P.-H. Fabre, C. H. Graham, G. R. Graves, K. A. Jønsson, D. Nogués-Bravo, Z. Wang, R. J. Whittaker, J. Fjeldså, and C. Rahbek. 2013. An Update of Wallace’s Zoogeographic Regions of the World 339:6.

Hubálek, Z. 1982. Coefficients of association and similarity, based on binary (presence-absence) data: an evaluation. Biological Reviews 57:669–689.

Hubert, L. J., and J. R. Levin. 1976. A general statistical framework for assessing categorical clustering in free recall. Psychological bulletin 83:1072.

Huijnen, V., M. J. Wooster, J. W. Kaiser, D. L. A. Gaveau, J. Flemming, M. Parrington, A. Inness, D. Murdiyarso, B. Main, and M. van Weele. 2016. Fire carbon emissions over maritime southeast Asia in 2015 largest since 1997. Scientific Reports 6:26886.

Hurlbert, A. H., and W. Jetz. 2007. Species richness, hotspots, and the scale dependence of range maps in ecology and conservation. Proceedings of the National Academy of Sciences 104:13384–13389.

Jaccard, P. 1901. Étude comparative de la distribution florale dans une portion des Alpes et des Jura. Bull Soc Vaudoise Sci Nat 37:547–579.

Jain, A. K., M. N. Murty, and P. J. Flynn. 1999. Data clustering: a review. ACM Computing Surveys 31:264–323.

Jetz, W., and C. Rahbek. 2002. Geographic Range Size and Determinants of Avian Species Richness. Science 297:1548–1551.

Jetz, W., C. H. Sekercioglu, and J. E. M. Watson. 2008. Ecological Correlates and Conservation Implications of Overestimating Species Geographic Ranges: *Overestimation of Species Ranges*. Conservation Biology 22:110–119.

Jongman, R. H., C. J. F. ter Braak, and O. F. R. van Tongeren. 1995. Data analysis in community and landscape ecology. Cambridge university press.

Kahneman, D., and A. Tversky. 1972. Subjective probability: A judgment of representativeness. Cognitive Psychology 3:430–454.

Karger, D. N., O. Conrad, J. Böhner, T. Kawohl, H. Kreft, R. W. Soria-Auza, N. E. Zimmermann, H. P. Linder, and M. Kessler. 2017. Climatologies at high resolution for the earth’s land surface areas. Scientific Data 4:170122.

Karger, D. N., M. Kessler, M. Lehnert, and W. Jetz. 2021. Limited protection and ongoing loss of tropical cloud forest biodiversity and ecosystems worldwide. Nature Ecology & Evolution 5:854–862.

Kass, J. M., R. Muscarella, P. J. Galante, C. L. Bohl, G. E. Pinilla-Buitrago, R. A. Boria, M. Soley-Guardia, and R. P. Anderson. 2021. ENMeval 2.0: Redesigned for customizable and reproducible modeling of species’ niches and distributions. Methods in Ecology and Evolution 12:1602–1608.

Kaufman, L., and P. J. Rousseeuw. 2005. Finding groups in data: an introduction to cluster analysis. Wiley, Hoboken, N.J.

Keddy, P. A. 1992. Assembly and response rules: two goals for predictive community ecology. Journal of Vegetation Science 3:157–164.

Keil, P., T. Wiegand, A. B. Tóth, D. J. McGlinn, and J. M. Chase. 2021. Measurement and analysis of interspecific spatial associations as a facet of biodiversity. Ecological Monographs.

Kramer-Schadt, S., J. Niedballa, J. D. Pilgrim, B. Schröder, J. Lindenborn, V. Reinfelder, M. Stillfried, I. Heckmann, A. K. Scharf, D. M. Augeri, S. M. Cheyne, A. J. Hearn, J. Ross, D. W. Macdonald, J. Mathai, J. Eaton, A. J. Marshall, G. Semiadi, R. Rustam, H. Bernard, R. Alfred, H. Samejima, J. W. Duckworth, C. Breitenmoser-Wuersten, J. L. Belant, H. Hofer, and A. Wilting. 2013. The importance of correcting for sampling bias in MaxEnt species distribution models. Diversity and Distributions 19:1366–1379.

Kreft, H., and W. Jetz. 2010. A framework for delineating biogeographical regions based on species distributions: Global quantitative biogeographical regionalizations. Journal of Biogeography 37:2029–2053.

Krosby, M., C. B. Wilsey, J. L. McGuire, J. M. Duggan, T. M. Nogeire, J. A. Heinrichs, J. J. Tewksbury, and J. J. Lawler. 2015. Climate-induced range overlap among closely related species. Nature Climate Change 5:883–886.

Langner, A., and F. Siegert. 2009. Spatiotemporal fire occurrence in Borneo over a period of 10 years. Global Change Biology 15:48–62.

Ledo, A. 2015. Nature and Age of Neighbours Matter: Interspecific Associations among Tree Species Exist and Vary across Life Stages in Tropical Forests. PLOS ONE 10:e0141387.

Legendre, P., and L. Legendre. 2012. Numerical ecology. Elsevier.

Lennon, J. J., P. Koleff, J. J. D. Greenwood, and K. J. Gaston. 2003. Contribution of rarity and commonness to patterns of species richness: Richness patterns and rarity/commonness. Ecology Letters 7:81–87.

Leroy, B., M. S. Dias, E. Giraud, B. Hugueny, C. Jézéquel, F. Leprieur, T. Oberdorff, and P. A. Tedesco. 2019. Global biogeographical regions of freshwater fish species. Journal of Biogeography 46:2407–2419.

Liu, C., G. Newell, and M. White. 2016. On the selection of thresholds for predicting species occurrence with presence-only data. Ecology and Evolution 6:337–348.

Lomolino, M. V. 2004. Conservation biogeography. Frontiers of Biogeography: new directions in the geography of nature 293.

Lomolino, M. V., B. R. C. Riddle, and J. H. C. Brown. 2006. Biogeography. Sinauer Associates, Inc,.

Ludwig, J. A., J. F. Reynolds, L. Quartet, and J. F. Reynolds. 1988. Statistical ecology: a primer in methods and computing. John Wiley & Sons.

Mainali, K., T. Hefley, L. Ries, and W. F. Fagan. 2020. Matching expert range maps with species distribution model predictions. Conservation Biology 34:1292–1304.

Manchego, C. E., P. Hildebrandt, J. Cueva, C. I. Espinosa, B. Stimm, and S. Günter. 2017. Climate change versus deforestation: Implications for tree species distribution in the dry forests of southern Ecuador. PloS one 13:e0190092.

Marquet, P. A., M. Fernandez, S. A. Navarrete, and C. Valdovinos. 2004. Diversity emerging: Toward a deconstruction of biodiversity patterns. Page Frontiers of biogeography: new directions in the geography of nature.

Marshall, L., J. C. Biesmeijer, P. Rasmont, N. J. Vereecken, L. Dvorak, U. Fitzpatrick, F. Francis, J. Neumayer, F. Ødegaard, J. P. T. Paukkunen, T. Pawlikowski, M. Reemer, S. P. M. Roberts, J. Straka, S. Vray, and N. Dendoncker. 2018. The interplay of climate and land use change affects the distribution of EU bumblebees. Global Change Biology 24:101–116.

Merow, C., M. J. Smith, and J. A. Silander. 2013. A practical guide to MaxEnt for modeling species’ distributions: what it does, and why inputs and settings matter. Ecography 36:1058–1069.

Miettinen, J., C. Shi, and S. C. Liew. 2011. Deforestation rates in insular Southeast Asia between 2000 and 2010: DEFORESTATION IN INSULAR SOUTHEAST ASIA 2000-2010. Global Change Biology 17:2261–2270.

Milanesi, P., F. Della Rocca, and R. A. Robinson. 2020. Integrating dynamic environmental predictors and species occurrences: Toward true dynamic species distribution models. Ecology and Evolution 10:1087–1092.

Milligan, G. W., and M. C. Cooper. 1985. An examination of procedures for determining the number of clusters in a data set. Psychometrika 50:159–179.

Minchin, P. R. 1987. An evaluation of the relative robustness of techniques for ecological ordination. Vegetatio 69:89–107.

Miyamoto, S. 2012. An Overview of Hierarchical and Non-hierarchical Algorithms of Clustering for Semi-supervised Classification. Pages 1–10 in V. Torra, Y. Narukawa, B. López, and M. Villaret, editors. Modeling Decisions for Artificial Intelligence. Springer, Berlin, Heidelberg.

Morales, N. S., I. C. Fernández, and V. Baca-González. 2017. MaxEnt’s parameter configuration and small samples: are we paying attention to recommendations? A systematic review. PeerJ 5:e3093.

Neo, L., H. T. W. Tan, and K. M. Wong. 2021. Centres of endemism in Borneo and their environmental correlates revealed by endemic plant genera. Flora 285:151966.

Newbold, T. 2018. Future effects of climate and land-use change on terrestrial vertebrate community diversity under different scenarios. Proceedings of the Royal Society B: Biological Sciences 285:20180792.

NOAA. 2022. Southern Oscillation Index (SOI) | El Niño/Southern Oscillation (ENSO). Monitoring. https://www.ncei.noaa.gov/access/monitoring/enso/soi.

Norberg, A., N. Abrego, F. G. Blanchet, F. R. Adler, B. J. Anderson, J. Anttila, M. B. Araújo, T. Dallas, D. Dunson, J. Elith, S.D. Foster, R. Fox, J. Franklin, W. Godsoe, A. Guisan, B. O’Hara, N. A. Hill, R. D. Holt, F. K. C. Hui, M. Husby, J. A. Kålås, A. Lehikoinen, M. Luoto, H. K. Mod, G. Newell, I. Renner, T. Roslin, J. Soininen, W. Thuiller, J. Vanhatalo, D. Warton, M. White, N. E. Zimmermann, D. Gravel, and O. Ovaskainen. 2019. A comprehensive evaluation of predictive performance of 33 species distribution models at species and community levels. Ecological Monographs 89:1–24.

Oksanen, J., R. Kindt, P. Legendre, B. O’Hara, M. H. H. Stevens, M. J. Oksanen, and M. Suggests. 2007. The vegan package. Community ecology package 10:719.

Olson, D. M., E. Dinerstein, E. D. Wikramanayake, N. D. Burgess, G. V. Powell, E. C. Underwood, J. A. D’amico, I. Itoua, H. E. Strand, and J. C. Morrison. 2001. Terrestrial Ecoregions of the World: A New Map of Life on Earth. BioScience 51:933–938.

Ovaskainen, O., G. Tikhonov, A. Norberg, F. Guillaume Blanchet, L. Duan, D. Dunson, T. Roslin, and N. Abrego. 2017. How to make more out of community data? A conceptual framework and its implementation as models and software. Ecology Letters 20:561–576.

Owen-Smith, N., J. Martin, and K. Yoganand. 2015. Spatially nested niche partitioning between syntopic grazers at foraging arena scale within overlapping home ranges. Ecosphere 6:art152.

Pang, S. E. H., J. D. T. De Alban, and E. L. Webb. 2021. Effects of climate change and land cover on the distributions of a critical tree family in the Philippines. Scientific Reports 11:276.

Pang, S. E. H., Y. Zeng, J. D. T. De Alban, and E. L. Webb. 2022. Occurrence–habitat mismatching and niche truncation when modelling distributions affected by anthropogenic range contractions. Diversity and Distributions:ddi.13544.

Peterson, A. T., editor. 2011. Ecological niches and geographic distributions. Princeton University Press, Princeton, N.J.

Peterson, A. T., and J. Soberón. 2012. Species Distribution Modeling and Ecological Niche Modeling: Getting the Concepts Right. Natureza & Conservação 10:102–107.

Peterson, A. T., J. Soberón, J. Ramsey, and L. Osorio-Olvera. 2020. Co-occurrence Networks do not Support Identification of Biotic Interactions. Biodiversity Informatics 15:1–10.

Phillips, S. J., R. P. Anderson, M. Dudík, R. E. Schapire, and M. E. Blair. 2017. Opening the black box: an open-source release of Maxent. Ecography 40:887–893.

Phillips, S. J., R. P. Anderson, and R. E. Schapire. 2006. Maximum entropy modeling of species geographic distributions. Ecological Modelling 190:231–259.

Phua, M.-H., S. Tsuyuki, N. Furuya, and J. S. Lee. 2008. Detecting deforestation with a spectral change detection approach using multitemporal Landsat data: A case study of Kinabalu Park, Sabah, Malaysia. Journal of Environmental Management 88:784–795.

Pompe, S., J. Hanspach, F.-W. Badeck, S. Klotz, H. Bruelheide, and I. Kühn. 2010. Investigating habitat-specific plant species pools under climate change. Basic and Applied Ecology 11:603–611.

Possingham, H. P., H. Grantham, and C. Rondinini. 2007. How can you conserve species that haven’t been found?: Commentary. Journal of Biogeography 34:758–759.

Prendergast, R. 1993. Rare species, the coincidence of diversity hotspots and conservation strategies. Nature 365:335–337.

R Core Team. 2013. R: A language and environment for statistical computing. R Foundation for Statistical Computing, Vienna, Austria 55:275–286.

Raes, N., M. C. Roos, J. W. F. Slik, E. E. Van Loon, and H. ter Steege. 2009. Botanical richness and endemicity patterns of Borneo derived from species distribution models. Ecography 32:180–192.

Rajalingam, D. N., and K. Ranjini. 2011. Hierarchical Clustering Algorithm - A Comparative Study. International Journal of Computer Applications 19:42–46.

Roberge, J.-M., and P. Angelstam. 2004. Usefulness of the Umbrella Species Concept as a Conservation Tool. Conservation Biology 18:76–85.

Roberts, D. W. 1987. A Dynamical Systems Perspective on Vegetation Theory. Vegetatio 69:27–33.

Rousseeuw, P. J. 1985. A visual display for hierarchical classification.

Rousseeuw, P. J. 1987. Silhouettes: a graphical aid to the interpretation and validation of cluster analysis. Journal of computational and applied mathematics 20:53–65.

Roux, M. 2018. A Comparative Study of Divisive and Agglomerative Hierarchical Clustering Algorithms. Journal of Classification 35:345–366.

Roxburgh, S. H., and P. Chesson. 1998. A New Method for Detecting Species Associations with Spatially Autocorrelated Data. Ecology 79:2180–2192.

Royle, J. A., R. B. Chandler, C. Yackulic, and J. D. Nichols. 2012. Likelihood analysis of species occurrence probability from presence-only data for modelling species distributions. Methods in Ecology and Evolution 3:545–554.

Rüger, N., R. Condit, D. H. Dent, S. J. DeWalt, S. P. Hubbell, J. W. Lichstein, O. R. Lopez, C. Wirth, and C. E. Farrior. 2020. Demographic trade-offs predict tropical forest dynamics. Science 368:165–168.

Sa’adi, Z., S. Shahid, and M. S. Shiru. 2021. Defining climate zone of Borneo based on cluster analysis. Theoretical and Applied Climatology 145:1467–1484.

Salvador, S., and P. Chan. 2004. Determining the number of clusters/segments in hierarchical clustering/segmentation algorithms. Pages 576–584 16th IEEE International Conference on Tools with Artificial Intelligence. IEEE Comput. Soc, Boca Raton, FL, USA.

Santini, L., L. H. Antão, M. Jung, A. Benítez-López, G. Rapacciuolo, M. Di Marco, F. A. M. Jones, J. M. Haghkerdar, and M. González-Suárez. 2021. The interface between Macroecology and Conservation: existing links and untapped opportunities. Frontiers of Biogeography 0.

Seif, G. 2018. The 5 clustering algorithms data scientists need to know. https://towardsdatascience.com/the-5-clustering-algorithms-data-scientists-need-to-know-a36d136ef68.

Shipley, B., and P. A. Keddy. 1987. The individualistic and community-unit concepts as falsifiable hypotheses. Pages 47–55 Theory and models in vegetation science. Springer.

Siman, K., D. A. Friess, M. Huxham, S. McGowan, J. Drewer, L. P. Koh, Y. Zeng, A. M. Lechner, J. S. H. Lee, C. D. Evans, S. Evers, A. Jonay, H. Varkkey, G. Anshari, A. Jaya, K. Y. Chong, S. Page, S. Mishra, and S. Sjögersten. 2021. Nature-based Solutions for Climate Change Mitigation: Challenges and Opportunities for the ASEAN Region. British High Commission and the COP26 Universities Network:1–36.

Singh, N., and D. Singh. 2012. Performance Evaluation of K-Means and Heirarichal Clustering in Terms of Accuracy and Running Time 3:3.

Slik, J. W. F. 2009. Plants of Southeast Asia. https://asianplant.net/.

Slik, J. W. F., S.-I. Aiba, M. Bastian, F. Q. Brearley, C. H. Cannon, K. A. O. Eichhorn, G. Fredriksson, K. Kartawinata, Y. Laumonier, A. Mansor, A. Marjokorpi, E. Meijaard, R. J. Morley, H. Nagamasu, R. Nilus, E. Nurtjahya, J. Payne, A. Permana, A. D. Poulsen, N. Raes, S. Riswan, C. P. van Schaik, D. Sheil, K. Sidiyasa, E. Suzuki, J. L. C. H. van Valkenburg, C. O. Webb, S. Wich, T. Yoneda, R. Zakaria, and N. Zweifel. 2011. Soils on exposed Sunda Shelf shaped biogeographic patterns in the equatorial forests of Southeast Asia. Proceedings of the National Academy of Sciences 108:12343–12347.

Slik, J. W. F., A. D. Poulsen, P. S. Ashton, C. H. Cannon, K. A. O. Eichhorn, K. Kartawinata, I. Lanniari, H. Nagamasu, M. Nakagawa, M. G. L. van Nieuwstadt, J. Payne, Purwaningsih, A. Saridan, K. Sidiyasa, R. W. Verburg, C. O. Webb, and P. Wilkie. 2003. A floristic analysis of the lowland dipterocarp forests of Borneo. Journal of Biogeography 30:1517–1531.

Slik, J. W. F., N. Raes, S.-I. Aiba, F. Q. Brearley, C. H. Cannon, E. Meijaard, H. Nagamasu, R. Nilus, G. Paoli, A. D. Poulsen, D. Sheil, E. Suzuki, J. L. C. H. van Valkenburg, C. O. Webb, P. Wilkie, and S. Wulffraat. 2009. Environmental correlates for tropical tree diversity and distribution patterns in Borneo. Diversity and Distributions 15:523–532.

Sloan, S., P. Meyfroidt, T. K. Rudel, F. Bongers, and R. Chazdon. 2019. The forest transformation: Planted tree cover and regional dynamics of tree gains and losses. Global Environmental Change 59:101988.

Soberon, J., and A. T. Peterson. 2005. Interpretation of Models of Fundamental Ecological Niches and Species’ Distributional Areas. Biodiversity Informatics 2:1–10.

Sokal, R. R., and C. D. Michener. 1958. A statistical method for evaluating systematic relationships. Univ. Kansas, Sci. Bull. 38:1409–1438.

Sokal, R. R., and F. J. Rohlf. 1962. The comparison of dendrograms by objective methods. TAXON 11:33–40.

Stolar, J., and S. E. Nielsen. 2015. Accounting for spatially biased sampling effort in presence-only species distribution modelling. Diversity and Distributions 21:595–608.

Struebig, M. J., A. Wilting, D. L. A. Gaveau, E. Meijaard, R. J. Smith, M. Fischer, K. Metcalfe, S. Kramer-Schadt, T. Abdullah, N. Abram, R. Alfred, M. Ancrenaz, D. M. Augeri, J. L. Belant, H. Bernard, M. Bezuijen, A. Boonman, R. Boonratana, T. Boorsma, C. Breitenmoser-Würsten, J. Brodie, S. M. Cheyne, C. Devens, J. W. Duckworth, N. Duplaix, J. Eaton, C. Francis, G. Fredriksson, A. J. Giordano, C. Gonner, J. Hall, M. E. Harrison, A. J. Hearn, I. Heckmann, M. Heydon, H. Hofer, J. Hon, S. Husson, F. A. Anwarali Khan, T. Kingston, D. Kreb, M. Lammertink, D. Lane, F. Lasmana, L. B. Liat, N. T.-L. Lim, J. Lindenborn, B. Loken, D. W. Macdonald, A. J. Marshall, I. Maryanto, J. Mathai, W. J. McShea, A. Mohamed, M. Nakabayashi, Y. Nakashima, J. Niedballa, S. Noerfahmy, S. Persey, A. Peter, S. Pieterse, J. D. Pilgrim, E. Pollard, S. Purnama, A. Rafiastanto, V. Reinfelder, C. Reusch, C. Robson, J. Ross, R. Rustam, L. Sadikin, H. Samejima, E. Santosa, I. Sapari, H. Sasaki, A. K. Scharf, G. Semiadi, C. R. Shepherd, R. Sykes, T. van Berkel, K. Wells, B. Wielstra, and A. Wong. 2015. Targeted Conservation to Safeguard a Biodiversity Hotspot from Climate and Land-Cover Change. Current Biology 25:372–378.

Sullivan, M. J. P., S. L. Lewis, K. Affum-Baffoe, C. Castilho, F. Costa, A. C. Sanchez, C. E. N. Ewango, W. Hubau, B. Marimon, A. Monteagudo-Mendoza, L. Qie, B. Sonké, R. V. Martinez, T. R. Baker, R. J. W. Brienen, T. R. Feldpausch, D. Galbraith, M. Gloor, Y. Malhi, S.-I. Aiba, M. N. Alexiades, E. C. Almeida, E. A. de Oliveira, E. Á. Dávila, P. A. Loayza, A. Andrade, S. A. Vieira, L. E. O. C. Aragão, A. Araujo-Murakami, E. J. M. M. Arets, L. Arroyo, P. Ashton, G. Aymard C., F. B. Baccaro, L. F. Banin, C. Baraloto, P. B. Camargo, J. Barlow, J. Barroso, J.-F. Bastin, S. A. Batterman, H. Beeckman, S. K. Begne, A. C. Bennett, E. Berenguer, N. Berry, L. Blanc, P. Boeckx, J. Bogaert, D. Bonal, F. Bongers, M. Bradford, F. Q. Brearley, T. Brncic, F. Brown, B. Burban, J. L. Camargo, W. Castro, C. Céron, S. C. Ribeiro, V. C. Moscoso, J. Chave, E. Chezeaux, C. J. Clark, F. C. de Souza, M. Collins, J. A. Comiskey, F. C. Valverde, M. C. Medina, L. da Costa, M. Dančák, G. C. Dargie, S. Davies, N. D. Cardozo, T. de Haulleville, M. B. de Medeiros, J. del Aguila Pasquel, G. Derroire, A. Di Fiore, J.-L. Doucet, A. Dourdain, V. Droissart, L. F. Duque, R. Ekoungoulou, F. Elias, T. Erwin, A. Esquivel-Muelbert, S. Fauset, J. Ferreira, G. F. Llampazo, E. Foli, A. Ford, M. Gilpin, J. S. Hall, K. C. Hamer, A. C. Hamilton, D. J. Harris, T. B. Hart, R. Hédl, B. Herault, R. Herrera, N. Higuchi, A. Hladik, E. H. Coronado, I. Huamantupa-Chuquimaco, W. H. Huasco, K. J. Jeffery, E. Jimenez-Rojas, M. Kalamandeen, M. N. K. Djuikouo, E. Kearsley, R. K. Umetsu, L. K. Kho, T. Killeen, K. Kitayama, B. Klitgaard, A. Koch, N. Labrière, W. Laurance, S. Laurance, M. E. Leal, A. Levesley, A. J. N. Lima, J. Lisingo, A. P. Lopes, G. Lopez-Gonzalez, T. Lovejoy, J. C. Lovett, R. Lowe, W. E. Magnusson, J. Malumbres-Olarte, Â. G. Manzatto, B. H. Marimon, A. R. Marshall, T. Marthews, S. M. de Almeida Reis, C. Maycock, K. Melgaço, C. Mendoza, F. Metali, V. Mihindou, W. Milliken, E. T. A. Mitchard, P. S. Morandi, H. L. Mossman, L. Nagy, H. Nascimento, D. Neill, R. Nilus, P. N. Vargas, W. Palacios, N. P. Camacho, J. Peacock, C. Pendry, M. C. Peñuela Mora, G. C. Pickavance, J. Pipoly, N. Pitman, M. Playfair, L. Poorter, J. R. Poulsen, A. D. Poulsen, R. Preziosi, A. Prieto, R. B. Primack, H. Ramírez-Angulo, J. Reitsma, M. Réjou-Méchain, Z. R. Correa, T. R. de Sousa, L. R. Bayona, A. Roopsind, A. Rudas, E. Rutishauser, K. Abu Salim, R. P. Salomão, J. Schietti, D. Sheil, R. C. Silva, J. S. Espejo, C. S. Valeria, M. Silveira, M. Simo-Droissart, M. F. Simon, J. Singh, Y. C. Soto Shareva, C. Stahl, J. Stropp, R. Sukri, T. Sunderland, M. Svátek, M. D. Swaine, V. Swamy, H. Taedoumg, J. Talbot, J. Taplin, D. Taylor, H. ter Steege, J. Terborgh, R. Thomas, S. C. Thomas, A. Torres-Lezama, P. Umunay, L. V. Gamarra, G. van der Heijden, P. van der Hout, P. van der Meer, M. van Nieuwstadt, H. Verbeeck, R. Vernimmen, A. Vicentini, I. C. G. Vieira, E. V. Torre, J. Vleminckx, V. Vos, O. Wang, L. J. T. White, S. Willcock, J. T. Woods, V. Wortel, K. Young, R. Zagt, L. Zemagho, P. A. Zuidema, J. A. Zwerts, and O. L. Phillips. 2020. Long-term thermal sensitivity of Earth’s tropical forests. Science 368:869–874.

Sullivan, M. J. P., J. Talbot, S. L. Lewis, O. L. Phillips, L. Qie, S. K. Begne, J. Chave, A. Cuni-Sanchez, W. Hubau, G. Lopez-Gonzalez, L. Miles, A. Monteagudo-Mendoza, B. Sonké, T. Sunderland, H. ter Steege, L. J. T. White, K. Affum-Baffoe, S. Aiba, E. C. de Almeida, E. A. de Oliveira, P. Alvarez-Loayza, E. Á. Dávila, A. Andrade, L. E. O. C. Aragão, P. Ashton, G. A. Aymard C., T. R. Baker, M. Balinga, L. F. Banin, C. Baraloto, J.-F. Bastin, N. Berry, J. Bogaert, D. Bonal, F. Bongers, R. Brienen, J. L. C. Camargo, C. Cerón, V. C. Moscoso, E. Chezeaux, C. J. Clark, Á. C. Pacheco, J. A. Comiskey, F. C. Valverde, E. N. H. Coronado, G. Dargie, S. J. Davies, C. De Canniere, M. N. Djuikouo K., J.-L. Doucet, T. L. Erwin, J. S. Espejo, C. E. N. Ewango, S. Fauset, T. R. Feldpausch, R. Herrera, M. Gilpin, E. Gloor, J. S. Hall, D. J. Harris, T. B. Hart, K. Kartawinata, L. K. Kho, K. Kitayama, S. G. W. Laurance, W. F. Laurance, M. E. Leal, T. Lovejoy, J. C. Lovett, F. M. Lukasu, J.-R. Makana, Y. Malhi, L. Maracahipes, B. S. Marimon, B. H. M. Junior, A. R. Marshall, P. S. Morandi, J. T. Mukendi, J. Mukinzi, R. Nilus, P. N. Vargas, N. C. P. Camacho, G. Pardo, M. Peña-Claros, P. Pétronelli, G. C. Pickavance, A. D. Poulsen, J. R. Poulsen, R. B. Primack, H. Priyadi, C. A. Quesada, J. Reitsma, M. Réjou-Méchain, Z. Restrepo, E. Rutishauser, K. A. Salim, R. P. Salomão, I. Samsoedin, D. Sheil, R. Sierra, M. Silveira, J. W. F. Slik, L. Steel, H. Taedoumg, S. Tan, J. W. Terborgh, S. C. Thomas, M. Toledo, P. M. Umunay, L. V. Gamarra, I. C. G. Vieira, V. A. Vos, O. Wang, S. Willcock, and L. Zemagho. 2017. Diversity and carbon storage across the tropical forest biome. Scientific Reports 7:39102.

Tikhonov, G., N. Abrego, D. Dunson, and O. Ovaskainen. 2017. Using joint species distribution models for evaluating how species-to-species associations depend on the environmental context. Methods in Ecology and Evolution 8:443–452.

Titeux, N., K. Henle, J.-B. Mihoub, A. Regos, I. R. Geijzendorffer, W. Cramer, P. H. Verburg, and L. Brotons. 2017. Global scenarios for biodiversity need to better integrate climate and land use change. Diversity and Distributions 23:1231–1234.

Torres, R., N. I. Gasparri, P. G. Blendinger, and H. R. Grau. 2014. Land-use and land-cover effects on regional biodiversity distribution in a subtropical dry forest: a hierarchical integrative multi-taxa study. Regional Environmental Change 14:1549–1561.

Trabucco, A., and R. J. Zomer. 2010. Global Soil Water Balance Geospatial Database. CGIAR Consortium for Spatial Information.

Trabucco, A., and R. J. Zomer. 2018. Global aridity index and potential evapotranspiration (ET0) climate database v2. CGIAR Consortium for Spatial Information 10.

Trisos, C. H., C. Merow, and A. L. Pigot. 2020. The projected timing of abrupt ecological disruption from climate change. Nature.

Van der Laan, M., K. Pollard, and J. Bryan. 2003. A new partitioning around medoids algorithm. Journal of Statistical Computation and Simulation 73:575–584.

VanDerWal, J., L. P. Shoo, C. Graham, and S. E. Williams. 2009. Selecting pseudo-absence data for presence-only distribution modeling: How far should you stray from what you know? Ecological Modelling 220:589–594.

Velazco, S. J. E., F. Villalobos, F. Galvão, and P. De Marco Júnior. 2019. A dark scenario for Cerrado plant species: Effects of future climate, land use and protected areas ineffectiveness. Diversity and Distributions.

Villalobos, F., A. Lira-Noriega, J. Soberón, and H. T. Arita. 2013. Range–diversity plots for conservation assessments: Using richness and rarity in priority setting. Biological Conservation 158:313–320.

Villalobos, F., M. Á. Olalla-Tárraga, M. V. Cianciaruso, T. F. Rangel, and J. A. F. Diniz-Filho. 2017. Global patterns of mammalian co-occurrence: phylogenetic and body size structure within species ranges. Journal of Biogeography 44:136–146.

Vollering, J., R. Halvorsen, I. Auestad, and K. Rydgren. 2019. Bunching up the background betters bias in species distribution models. Ecography 42:1717–1727.

Ward, J. H. 1963. Hierarchical grouping to optimize an objective function. Journal of the American statistical association 58:236–244.

Warren, D. L., L. J. Beaumont, R. Dinnage, and J. B. Baumgartner. 2019. New methods for measuring ENM breadth and overlap in environmental space. Ecography 42:444–446.

Warren, D. L., R. E. Glor, and M. Turelli. 2008. Environmental niche equivalency versus conservatism: quantitative approaches to niche evolution. Evolution 62:2868–2883.

Webb, N. R. 1989. Studies on the invertebrate fauna of fragmented heathland in Dorset, UK, and the implications for conservation. Biological Conservation 47:153–165.

Wedley, W. C., B. Schoner, and E. U. Choo. 1993. Clustering, dependence and ratio scales in AHP: Rank reversals and incorrect priorities with a single criterion. Journal of Multi-Criteria Decision Analysis 2:145–158.

Westman, W. E. 1985. Xeric Mediterranean-type shrubland associations of Alta and Baja California and the community/continuum debate. Pages 79–95 in R. K. Peet, editor. Plant community ecology: Papers in honor of Robert H. Whittaker. Springer Netherlands, Dordrecht.

Whittaker, R. H. 1951. A criticism of the plant association and climatic climax concepts. Northwest Scientist 25:17–31.

Whittaker, R. H. 1953. A Consideration of Climax Theory: The Climax as a Population and Pattern. Ecological Monographs 23:41–78.

Whittaker, R. J., M. B. Araújo, P. Jepson, R. J. Ladle, J. E. M. Watson, and K. J. Willis. 2005. Conservation Biogeography: assessment and prospect. Diversity and Distributions 11:3–23.

Wirth, M., G. F. Estabrook, and D. J. Rogers. 1966. A graph theory model for systematic biology, with an example for the Oncidiinae (Orchidaceae). Systematic Zoology 15:59–69.

Wong, C. J., D. James, N. A. Besar, K. U. Kamlun, J. Tangah, S. Tsuyuki, and M.-H. Phua. 2020. Estimating Mangrove Above-Ground Biomass Loss Due to Deforestation in Malaysian Northern Borneo between 2000 and 2015 Using SRTM and Landsat Images. Forests 11:1018.

Wooster, M. J., G. L. W. Perry, and A. Zoumas. 2012. Fire, drought and El Niño relationships on Borneo (Southeast Asia) in the pre-MODIS era (1980–2000). Biogeosciences 9:317–340.

Wüest, R. O., N. E. Zimmermann, D. Zurell, J. M. Alexander, S. A. Fritz, C. Hof, H. Kreft, S. Normand, J. S. Cabral, E. Szekely, W. Thuiller, M. Wikelski, and D. N. Karger. 2020. Macroecology in the age of Big Data – Where to go from here? Journal of Biogeography 47:1–12.

Yanahan, A. D., and W. Moore. 2019. Impacts of 21st-century climate change on montane habitat in the Madrean Sky Island Archipelago. Diversity and Distributions 25:1625–1638.

Zurell, D., J. Franklin, C. König, P. J. Bouchet, C. F. Dormann, J. Elith, G. Fandos, X. Feng, G. Guillera-Arroita, A. Guisan, J. J. Lahoz-Monfort, P. J. Leitão, D. S. Park, A. T. Peterson, G. Rapacciuolo, D. R. Schmatz, B. Schröder, J. M. Serra-Diaz, W. Thuiller, K. L. Yates, N. E. Zimmermann, and C. Merow. 2020. A standard protocol for reporting species distribution models. Ecography 43:1261–1277.

