## Supporting Information for "The clustering of spatially associated species unravels patterns in Bornean tree species distributions"

#### Extended Methods

More detailed descriptions of the methods used for developing the species distribution models (see Fig. S1 for methods flowchart). The methods were divided into four parts: (1) study area; (2) environment and land-cover data; (3) occurrence data; and (4) species distribution modelling.

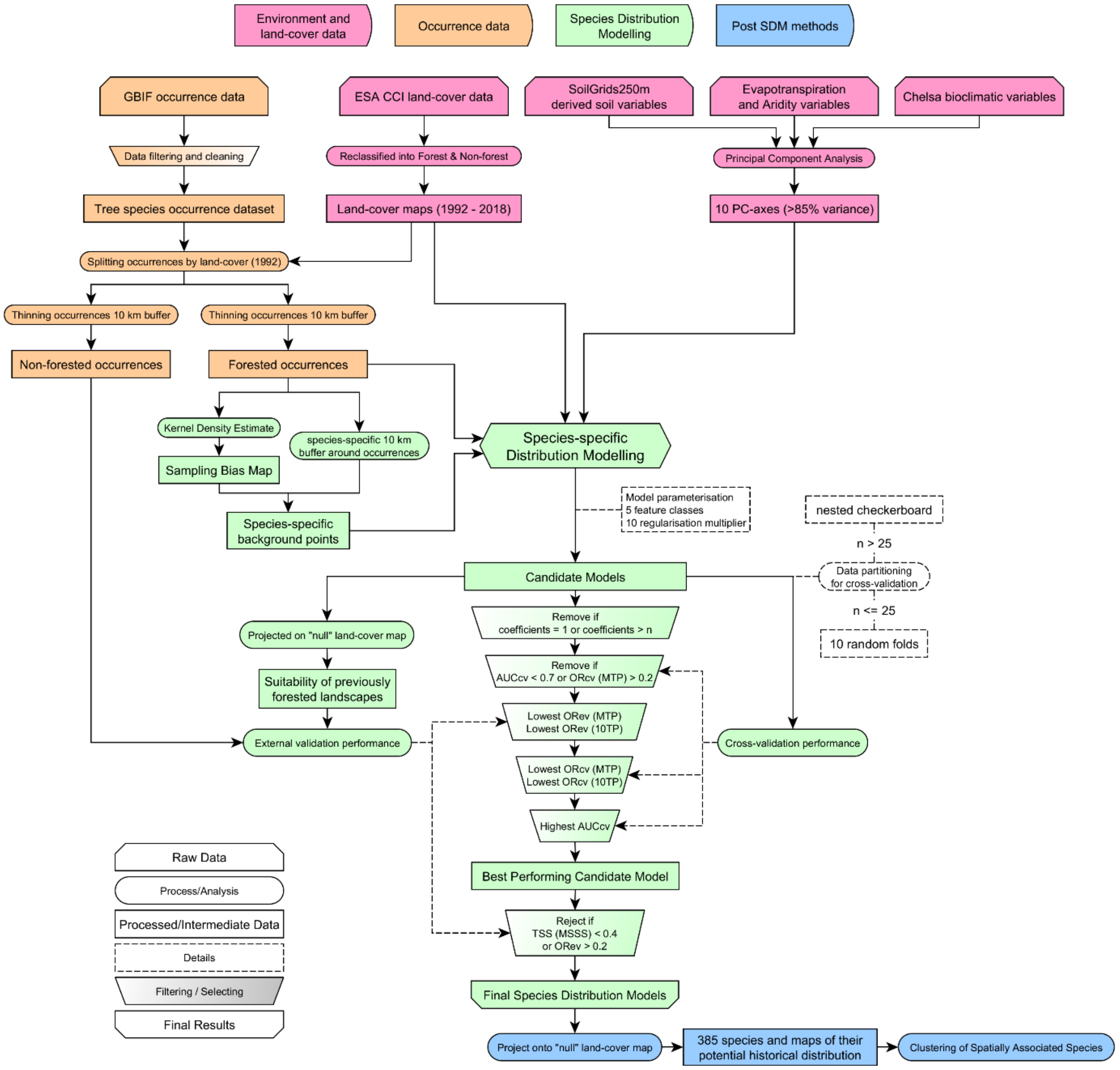


**Figure S1.** Methods flowchart for species distribution modelling.

### S1. Study Area

For our case study, we used the island of Borneo as our study area. Borneo is the third largest island in the world and the largest in Asia. It is located north of Java, west of Sulawesi, and east of peninsular Malaysia. The island is politically divided among Malaysia, Indonesia, and Brunei.

The island was formed via Mesozoic accretion of microcontinental fragment, ophiolite terranes and island arc crust onto a Paleozoic continental core. Biogeographically, Borneo is part of Sundaland, a large landmass that was exposed throughout the last 2.6 million years during periods of lower sea levels. Sundaland consists of what is known today as peninsular Malaysia and the larger islands of Borneo, Java, and Sumatra and their surrounding islands.

Borneo is located on the equator, with warm temperatures and high precipitation all year round. It consists of lowland tropical rainforests, peat swamp forests, Kerangas (heath forests), mangroves, montane rainforests and alpine meadows. The highest point of Borneo is Gunung Kinabalu at (4,095 m).

Borneo is plagued by a multitude of conservation challenges: extensive logging for lumber, large scale deforestation for oil palm plantations, and out of control human induced forest fires.

**S2. Environment and land-cover data**

In total, we obtained 44 environmental variables: 19 bioclimatic variables, 14 soil-water variables, 1 aridity index, 1 potential evapotranspiration variable, and 9 soil property variables (Table 1). Variables were cropped to the study region of Borneo, and resampled (bilinear) to 30 arc sec (~1 km). Areas with missing data for any variable were removed.

The variables were scaled and reduced using a Principal Component Analysis (PCA). The first 10 principal component (PC) axes were selected, which explained 87% of variance (selection threshold of above 85% cumulative proportion of variance).

Annual land-cover maps (year 1992 – 2020) were obtained from the European Space Agency (ESA) (Table 1). The land-cover maps were first reclassified following IPCC (Intergovernmental Panel on Climate Change) land categories into forest and non-forest, where non-forest consists of IPCC classes agriculture, grassland, wetland, settlement, shrubland, sparse vegetation, bare area, and water. The reclassified land-cover maps were then aggregated (mode) to 30 arc sec and resampled (nearest neighbour) to match the projection of environmental variables.

We used a binary categorisation of land-cover because of the study's focus on tree species, where a conservative approach towards identifying intact habitats was adopted. Areas with >50% forest cover was considered as intact and therefore suitable for tree species, or vice versa. Although natural non-forest habitats can exist in Borneo (e.g., Kinabalu alpine meadow), these areas were few and should not adversely affect models of tree species.

**Table S1.** List of environmental and land-cover data.

| Source & access data | Name | Full description | Native resolution |
|---|---|---|---|
| | Bio 01 | Annual mean temperature | |
| | Bio 02 | Mean Diurnal Range | |
| | Bio 03 | Isothermality | |
| | Bio 04 | Temperature Seasonality | |
| | Bio 05 | Max Temperature of Warmest Month | |
| | Bio 06 | Min Temperature of Coldest Month | |
| CHELSA v1.2, bioclimatic variables | Bio 07 | Temperature Annual Range | |
| for the time period 1973 – 2013 | Bio 08 | Mean Temperature of Wettest Quarter | 30 arcsec |
| [accessed 02/02/2020] | Bio 09 | Mean Temperature of Driest Quarter | (~ 1 km) |
| | Bio 10 | Mean Temperature of Warmest Quarter | |
| | Bio 11 | Mean Temperature of Coldest Quarter | |
| | Bio 12 | Annual Precipitation | |
| | Bio 13 | Precipitation of Wettest Month | |
| | Bio 14 | Precipitation of Driest Month | |
| | Bio 15 | Precipitation Seasonality | |
| | Bio 16 | Precipitation of Wettest Quarter | |

| | | | |
|---|---|---|---|
| | Bio 17 | Precipitation of Driest Quarter | |
| | Bio 18 | Precipitation of Warmest Quarter | |
| | Bio 19 | Precipitation of Coldest Quarter | |
| Global High-Resolution Soil-Water Balance v3 (Trabucco & Zomer, 2019) [accessed 18/11/2019] | AET | Mean Annual Actual Evapotranspiration rate | 30 arcsec (~ 1 km) |
| | ALPHA | Priestley-Talor Alpha Coefficient | |
| | SWC | Monthly Soil Water Content | |
| Global Aridity Index and Potential Evapo-Transpiration Climate Database v2 (Trabucco & Zomer, 2018) [accessed 19/11/2019] | AI | Aridity Index | 30 arcsec (~ 1 km) |
| | PET | Annual Potential Evapotranspiration | |
| SoilGrids250m v1 [accessed 30/12/2019] | AWC | Derived Available Water Capacity, volumetric fraction | 250 m |
| | BDRICM | Depth to bedrock (R horizon), up to 200 cm | |
| | BLDFIE | Bulk Density of Fine Earth, kg/m$^3$ | |
| | CECSOL | Cation Exchange Capacity of Soil in cmolc/kg | |
| | CLYPPT | Clay content, mass fraction | |
| | CRFVOL | Coarse Fragments, volumetric fraction | |
| | SNDPPT | Sand content, mass fraction | |
| | PHIHOX | Soil pH x 10 in $H_2O$ | |
| | PHIKCL | Soil pH x 10 in KCL | |
| European Space Agency, Climate Change Initiative [accessed 13/03/2017] | ESA_YEAR | Land Cover Maps v2.0.7, 1992 – 2015 | 300 m |
| European Space Agency, Climate Change Initiative [accessed 15/11/2021] | ESA_YEAR | Land Cover Maps v2.1.1, 2016 – 2020 | |

### S3. Occurrence data

The occurrence data was obtained from GBIF [https://doi.org/10.15468/dl.3sqcf4; accessed 18/11/2019]. Occurrence data was restricted to records from Malaysia, Indonesia and Brunei, from the year 1000 to 2019, with coordinates, without geospatial issues, and from Tracheophyta (vascular plants).

The occurrence data was then filtered to remove potentially ambiguous, erroneous or inaccurate records. First, occurrence data with an empty species tag (blank or "UNKNOWN" basis of record), coordinate uncertainty in meters exceeding 1 km, or locations outside the study area of Borneo, were removed. To further improve the precision and certainty in the coordinates of each occurrence point, records with coordinates (longitude and latitude) of less than two decimal places were considered "low accuracy" and removed. Species names were then checked for spelling errors and synonyms using the 'Taxonstand' package in R, and corrected. To restrict occurrence data to tree species, only the occurrence data of species identified in the GlobalTreeSearch database (Beech et al., 2017) were kept.

Our previous study, Pang et al. (2022), we found that SDMs developed using temporally dynamic anthropogenic predictors performed best (i.e., occurrence data associated with the anthropogenic predictor from the year it was sampled). However, the ESA land-cover data extended to 1992 only. If a minimum occurrence record of 10 is imposed, only 325 species would remain. Moreover, this was before any data thinning was applied; and with further model calibration and validation to remove underperforming models, even fewer species (ca. $n$ = 50) would remain. The main objective of our

SDMs was to identify the historical distribution of suitable habitats (i.e., before human intrusion), whereby calculations of interspecific spatial associations would be based upon. Therefore, the main requisite of our SDMs was to be able to correctly identify historically suitable habitats. Following the findings and recommendations in Pang et al., (2022), the use of a contemporary anthropogenic predictor can also allow accurate predictions of historical distributions, even with occurrence-habitat mismatching. Problems of unaccounted niche truncation due to anthropogenic range contractions, however, may still persist. As such, the occurrence data was further split before any further data cleaning was done.

The occurrence data was split into two datasets: occurrences that occurred within forested areas, and occurrences that occurred within non-forested areas. The within forested/non-forested areas designator was based on the ESA land-cover map for 1992, the most historical land-cover data available from that database. While occurrences within forested area were used as per usual (i.e., model calibration, training, and cross-validation), occurrences within non-forested areas were used exclusively to validate model projections of suitable habitat onto a manually calibrated “null” land-cover map. The null land-cover map was essentially the land-cover of Borneo under the assumption of no human intrusion, where the entire region was considered as forested. While excluding occurrences within non-forest areas from model training would remove any potential occurrence-habitat mismatching (e.g., occurrence sampled in 1967 from a forest plot being associated with the non-forested land-cover class in 1992), using those occurrences to validate model projections would test for potential truncations of the species’ pre-disturbance niche (i.e., omission rates would be high for truncated niche estimates).

The occurrence data of each species was first thinned using a 10 km buffer. This was done separately for each dataset, so that points from one dataset would not affect the thinning of the other. After thinning, species with fewer than 10 or 5 occurrences within forested or non-forested areas, respectively, were excluded. This was to ensure a sufficient number of occurrences for developing and validating species distribution models. In total, 743 tree species remained.

**S4. Species distribution modelling**

All species were individually modelled using the MaxEnt (3.4.1) algorithm via the ENMeval (2.0.0) package in R (Phillips et al. 2006, R Core Team 2013, Kass et al. 2021). A species-specific approach was adopted to maximise the reliability and performance of each SDM. The 10 PC-axes and the reclassified land-cover map for 1992 were used as predictors.

First, 10,000 background points were selected. Background points were restricted to areas that were at least 10 km away from any of the species’ occurrence points. To account for geographical sampling biases, the sampling probability of background points across the study area was based on an across species sampling bias map. To create the sampling bias map, occurrence data were combined to generate a spatial kernel density estimate with a Gaussian kernel bandwidth of 1°; using the *sp.kde* function from the ‘spatialEco’ package in R. This served to represent and account for the relative geographical sampling bias of occurrence points, i.e., areas with higher sampling bias were more likely to be sampled as background points.

For model tuning, candidate models were assessed based on the two datasets: cross-validation scores obtained from using the occurrences within forested areas (subscripted CV), and external validation scores obtained from using the occurrences within non-forested areas (subscripted EV). $Scores_{CV}$ were obtained by cross-validating models built from partitioned occurrence and background points. To reduce spatial autocorrelation and improve spatial independence between partitions, a nested checkerboard partitioning approach with an aggregation factor of 5 for both checkerboards was used for species with occurrence points greater than 25 (see *get.checkerboard2* function from the 'ENMeval' package in R). A 10-fold cross-validation approach was adopted for species with occurrence points less than or equal to 25, an approach that better identifies better performing and less overfitted models despite being developed with fewer samples.

$Scores_{EV}$ were obtained by projecting models onto a manually calibrated "null" land-cover map, and validating the resultant predictions of habitat suitability with the occurrence points from non-forested areas. The null land-cover map was essentially the land-cover of Borneo under the assumption of no human intrusion, where the entire region was considered as forested. Meaning, this validation tested the ability of the model to identify the suitability of previously forested landscapes. Additionally, model predictions of habitat suitability onto the null land-cover map was considered to represent the potential historical distribution of species, before any anthropogenic range contractions occurred. These projections, from the accepted final models, were the distribution maps later used to calculate interspecific spatial associations.

For the selection of MaxEnt parameters, an extensive round of species-specific tuning was performed for the model's feature class and regularisation multiplier. Feature classes tested were L, LQ, LQP, LQH, LQHP (Linear, Quadratic, Hinge and Product); regularisation multipliers tested were 0.1, 0.25, 0.5, 1, 1.5, 2, 3, 4, 6 and 8. To remove overly simplified or complex models, candidate models with number of coefficients equal to 1 or greater than the number of occurrences were removed. Candidate models with Area Under the Curve, $AUC_{CV} < 0.7$, or Omission Rates, $OR_{CV}$ (MTP—minimum training presence—threshold) > 0.2 were considered to have performed poorly and removed. If no candidate models remained at this point, the species was rejected. The best performing candidate model was selected based on five sequential criterions, where subsequent criterions were only applied when ties in best performing candidate models occurred. In sequence, the criterions for selecting the best performing model were: 1) lowest $OR_{EV}$, for MTP threshold; 2) lowest $OR_{EV}$, for 10TP—10% training presence—threshold; 3) lowest $OR_{CV}$, for MTP threshold; 4) lowest $OR_{CV}$, for 10TP threshold; and 5) highest $AUC_{CV}$. Finally, species whose best performing candidate model had True Skills Statistic (MSSS—maximum sum of sensitivity and specificity—threshold), TSS < 0.4, or $OR_{EV} > 0.2$, were also rejected. In total, 385 species distribution models were accepted.

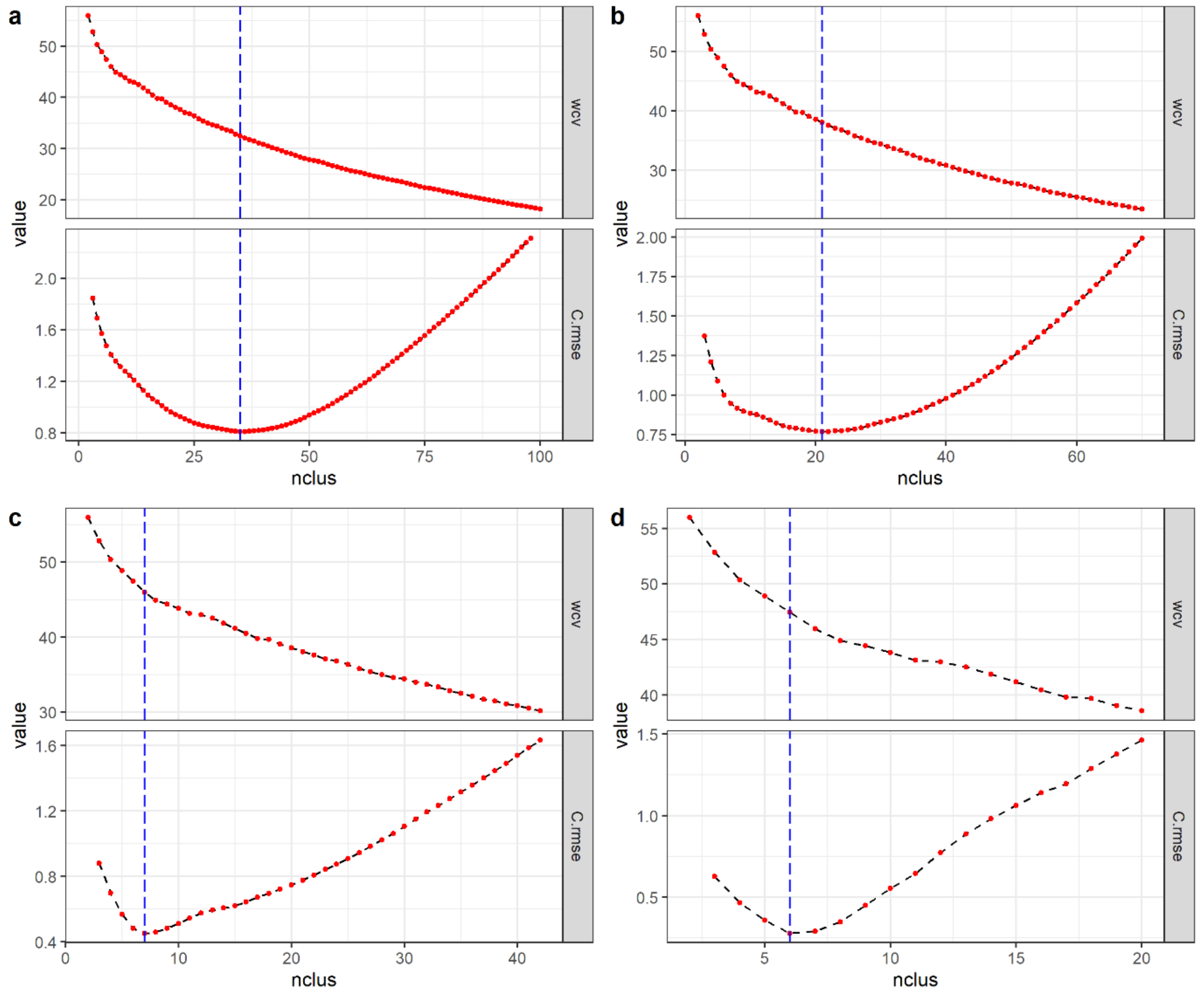


**Figure S2.** The diagnostic graph of within-cluster variance (wcv) across *k* number of clusters (nclus) for the clustering outcome resulting from WARD clustering algorithm based on Bray-Curtis dissimilarity. The vertical blue dash line indicates *k* at which the combine root mean square error (C.rmse) was the lowest; C.rmse was the combined RMSE of the linear regression of points left of *k* and right of *k* (for details, see Salvador and Chan 2004). The L-method was iterative, where the values of nclus were "cut-off" until the determined number of clusters *k* recurred; a-d shows this iteration where the nclus examined were 2-100, 2-70, 2-42, and 2-20.

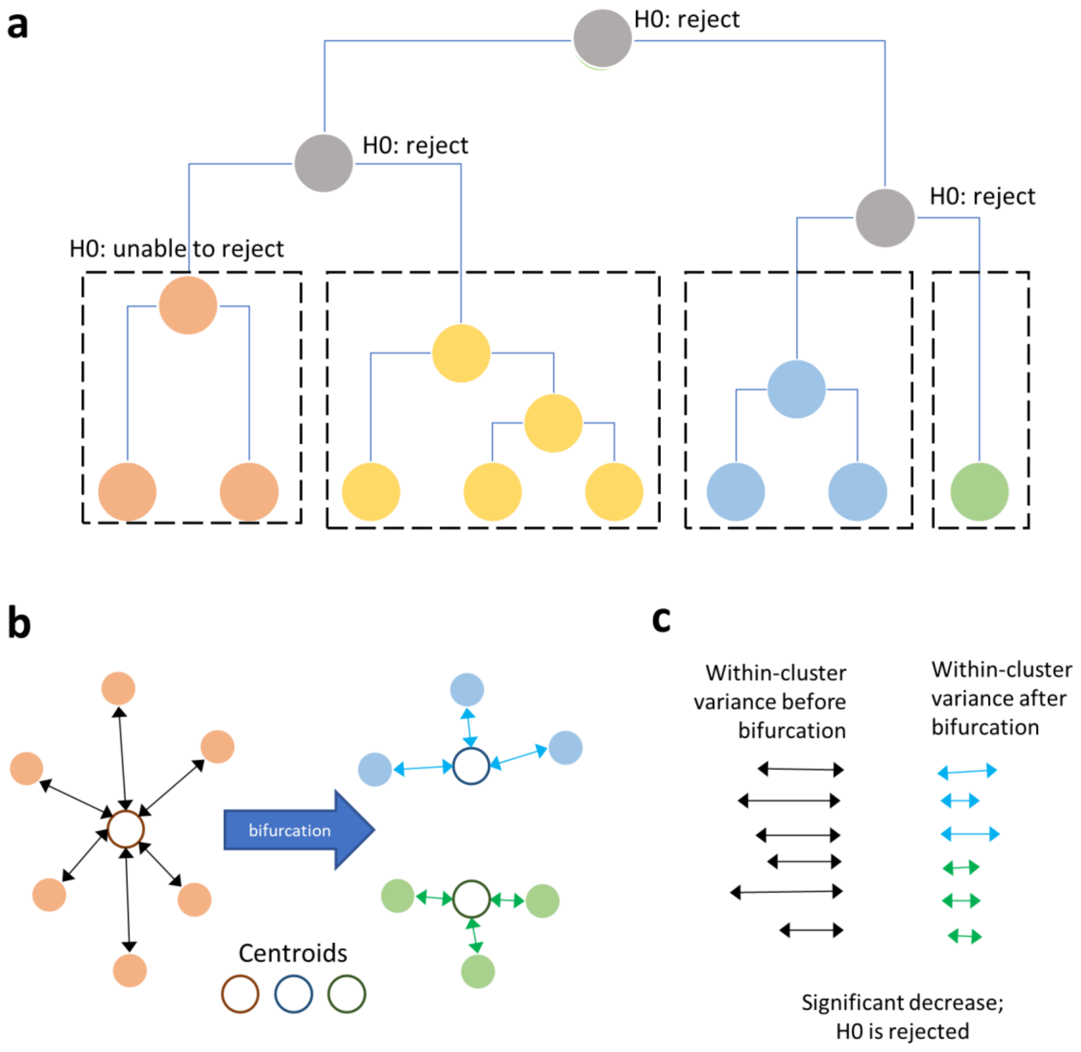


**Figure S3.** Conceptualisation of the Bifurcation Paired T-test. (a) At each bifurcation of the dendrogram, in increasing number of *k*, we tested for a significant change in a given evaluation metric. (b) Here, we examine within-cluster variance from the cluster's centroid. Showing an example bifurcation, a 6-object cluster is split into two 3-object clusters. The double-headed arrows indicate the distance/dissimilarity of each object from its cluster's centroid (i.e., the within-cluster variance). (c) Within-cluster variance before and after the bifurcation is then compared using a paired T-test. If there is a significant decrease in within-cluster variance, the bifurcation led to a significant improvement in within-cluster variance; thus, the 6-object cluster is less optimal than the two 3-object clusters and further bifurcation is required. However, if there is a non-significant decrease in within-cluster variance, the bifurcation led to a non-significant improvement; thus, objects are already optimally clustered, and no further bifurcation is required.

### Extended Results

**Table S2.** The optimal number of clusters for each dendrogram determined using different stopping rules. The number of clusters *k* assessed were from 2 to 100, i.e., *k* = 2 and *k* = 100 were the lowest and highest possible number of clusters, respectively. Stopping rules here were those considered within the main analysis.

| Data Type | Association index | Clustering Algorithm | Aggregated distribution (centroids) | | | Indicator distribution (medoids) | | | Object dissimilarity |
|---|---|---|---|---|---|---|---|---|---|
| | | | Bifurcation Pairwise T-test | L-method (BCV) | L-method (WCV) | Bifurcation Pairwise T-test | L-method (BCV) | L-method (WCV) | L-method (merging height) |
| Continuous | Spearman | UPGMA | 7 | 6 | 6 | 7 | 6 | 6 | 8 |
| | Spearman | CL | 17 | 8 | 6 | 9 | 4 | 6 | 9 |
| | Spearman | WARD | 37 | 3 | 5 | 20 | 4 | 5 | 7 |
| | Pearsons | UPGMA | 6 | 9 | 9 | 6 | 12 | 6 | 28 |
| | Pearsons | CL | 30 | 7 | 5 | 9 | 5 | 5 | 8 |
| | Pearsons | WARD | 38 | 8 | 7 | 19 | 11 | 8 | 9 |
| | Ruzicka | UPGMA | 4 | 9 | 9 | 4 | 9 | 9 | 4 |
| | Ruzicka | CL | 5 | 4 | 4 | 5 | 4 | 4 | 7 |
| | Ruzicka | WARD | 37 | 4 | 8 | 10 | 7 | 6 | 4 |
| | Bray-Curtis | UPGMA | 4 | 9 | 9 | 4 | 9 | 9 | 4 |
| | Bray-Curtis | CL | 5 | 4 | 4 | 5 | 4 | 4 | 7 |
| | Bray-Curtis | WARD | 34 | 5 | 6 | 11 | 4 | 6 | 4 |
| | Hellinger | UPGMA | 4 | 14 | 14 | 3 | 28 | 14 | 4 |
| | Hellinger | CL | 9 | 6 | 5 | 6 | 31 | 8 | 9 |
| | Hellinger | WARD | 10 | 6 | 7 | 7 | 11 | 9 | 8 |
| | Chi-squared | UPGMA | 3 | 7 | 40 | 3 | 28 | 7 | 4 |
| | Chi-squared | CL | 3 | 5 | 5 | 3 | 5 | 6 | 7 |
| | Chi-squared | WARD | 9 | 5 | 6 | 8 | 11 | 5 | 7 |
| Binary | Alroy | UPGMA | 7 | 4 | 3 | 4 | 38 | 3 | 6 |
| | Alroy | CL | 14 | 14 | 8 | 7 | 38 | 6 | 18 |
| | Alroy | WARD | 5 | 5 | 9 | 3 | 43 | 17 | 4 |
| | Tetrachloric | UPGMA | 9 | 11 | 8 | 4 | 8 | 8 | 7 |
| | Tetrachloric | CL | 27 | 4 | 5 | 15 | 4 | 5 | 4 |
| | Tetrachloric | WARD | 48 | 4 | 7 | 26 | 4 | 7 | 9 |
| | Jaccard | UPGMA | 8 | 15 | 15 | 4 | 15 | 15 | 24 |
| | Jaccard | CL | 10 | 4 | 7 | 10 | 7 | 8 | 22 |
| | Jaccard | WARD | 36 | 12 | 6 | 17 | 3 | 6 | 4 |
| | Dice-Sorensen | UPGMA | 3 | 13 | 13 | 3 | 19 | 13 | 24 |
| | Dice-Sorensen | CL | 10 | 5 | 5 | 3 | 7 | 9 | 22 |
| | Dice-Sorensen | WARD | 30 | 8 | 8 | 17 | 8 | 8 | 10 |
| | Scale C-score | UPGMA | 5 | 47 | 11 | 18 | 11 | 26 | 40 |
| | Scale C-score | CL | 8 | 37 | 6 | 6 | 3 | 16 | 6 |
| | Scale C-score | WARD | 12 | 3 | 8 | 6 | 5 | 5 | 10 |
| | Matching | UPGMA | 3 | 8 | 23 | 3 | 63 | 23 | 43 |
| | Matching | CL | 24 | 4 | 7 | 8 | 4 | 7 | 5 |
| | Matching | WARD | 29 | 3 | 5 | 7 | 3 | 5 | 6 |

**Table S3.** The optimal number of clusters for each dendrogram determined using different stopping rules. The number of clusters *k* assessed were from 2 to 100, i.e., *k* = 2 and *k* = 100 were the lowest and highest possible number of clusters, respectively. Stopping rules here were those not considered within the main analysis. Values of NA occurred when a significance test was required for the stopping rule, but all values of *k* were insignificant; applicable only to Duda & Hart ratio criterion.

| Data Type | Association index | Clustering Algorithm | Aggregated distribution (centroids) | | Indicator distribution (medoids) | | Object dissimilarity | |
|---|---|---|---|---|---|---|---|---|
| | | | Calinski & Harabasz – trace between/within | Duda & Hart – ratio criterion | Calinski & Harabasz – trace between/within | Duda & Hart – ratio criterion | Average Sillhoute | C-index |
| Continuous | Spearman | UPGMA | 2 | NA | 2 | NA | 35 | 100 |
| | Spearman | CL | 100 | NA | 2 | 52 | 92 | 96 |
| | Spearman | WARD | 3 | NA | 2 | NA | 100 | 100 |
| | Pearsons | UPGMA | 2 | NA | 2 | NA | 99 | 100 |
| | Pearsons | CL | 98 | NA | 2 | NA | 98 | 100 |
| | Pearsons | WARD | 2 | NA | 2 | NA | 100 | 80 |
| | Ruzicka | UPGMA | 10 | NA | 10 | NA | 2 | 82 |
| | Ruzicka | CL | 5 | NA | 5 | 73 | 2 | 11 |
| | Ruzicka | WARD | 2 | NA | 2 | NA | 2 | 2 |
| | Bray-Curtis | UPGMA | 3 | NA | 10 | NA | 2 | 80 |
| | Bray-Curtis | CL | 2 | NA | 5 | NA | 2 | 31 |
| | Bray-Curtis | WARD | 2 | NA | 2 | NA | 2 | 89 |
| | Hellinger | UPGMA | 3 | NA | 3 | NA | 2 | 78 |
| | Hellinger | CL | 2 | NA | 2 | 47 | 2 | 100 |
| | Hellinger | WARD | 2 | NA | 2 | NA | 4 | 100 |
| | Chi-squared | UPGMA | 3 | NA | 3 | NA | 2 | 80 |
| | Chi-squared | CL | 2 | NA | 2 | 76 | 2 | 100 |
| | Chi-squared | WARD | 2 | NA | 2 | NA | 2 | 100 |
| Binary | Alroy | UPGMA | 99 | NA | 96 | 5 | 95 | 100 |
| | Alroy | CL | 100 | 14 | 98 | 2 | 82 | 96 |
| | Alroy | WARD | 2 | 33 | 2 | 3 | 86 | 94 |
| | Tetrachloric | UPGMA | 2 | NA | 2 | NA | 91 | 100 |
| | Tetrachloric | CL | 2 | NA | 2 | 15 | 100 | 98 |
| | Tetrachloric | WARD | 2 | NA | 2 | NA | 98 | 78 |
| | Jaccard | UPGMA | 16 | NA | 16 | 34 | 2 | 2 |
| | Jaccard | CL | 2 | NA | 2 | NA | 80 | 100 |
| | Jaccard | WARD | 2 | 2 | 2 | 2 | 100 | 69 |
| | Dice-Sorensen | UPGMA | 14 | NA | 14 | NA | 2 | 100 |
| | Dice-Sorensen | CL | 2 | NA | 2 | 3 | 80 | 100 |
| | Dice-Sorensen | WARD | 2 | NA | 2 | NA | 99 | 100 |
| | Scale C-score | UPGMA | 100 | 49 | 7 | 4 | 36 | 100 |
| | Scale C-score | CL | 100 | 16 | 3 | 3 | 100 | 100 |
| | Scale C-score | WARD | 2 | NA | 2 | 6 | 100 | 100 |
| | Matching | UPGMA | 2 | NA | 2 | NA | 2 | 100 |
| | Matching | CL | 2 | NA | 2 | 71 | 2 | 95 |
| | Matching | WARD | 2 | NA | 2 | NA | 2 | 91 |

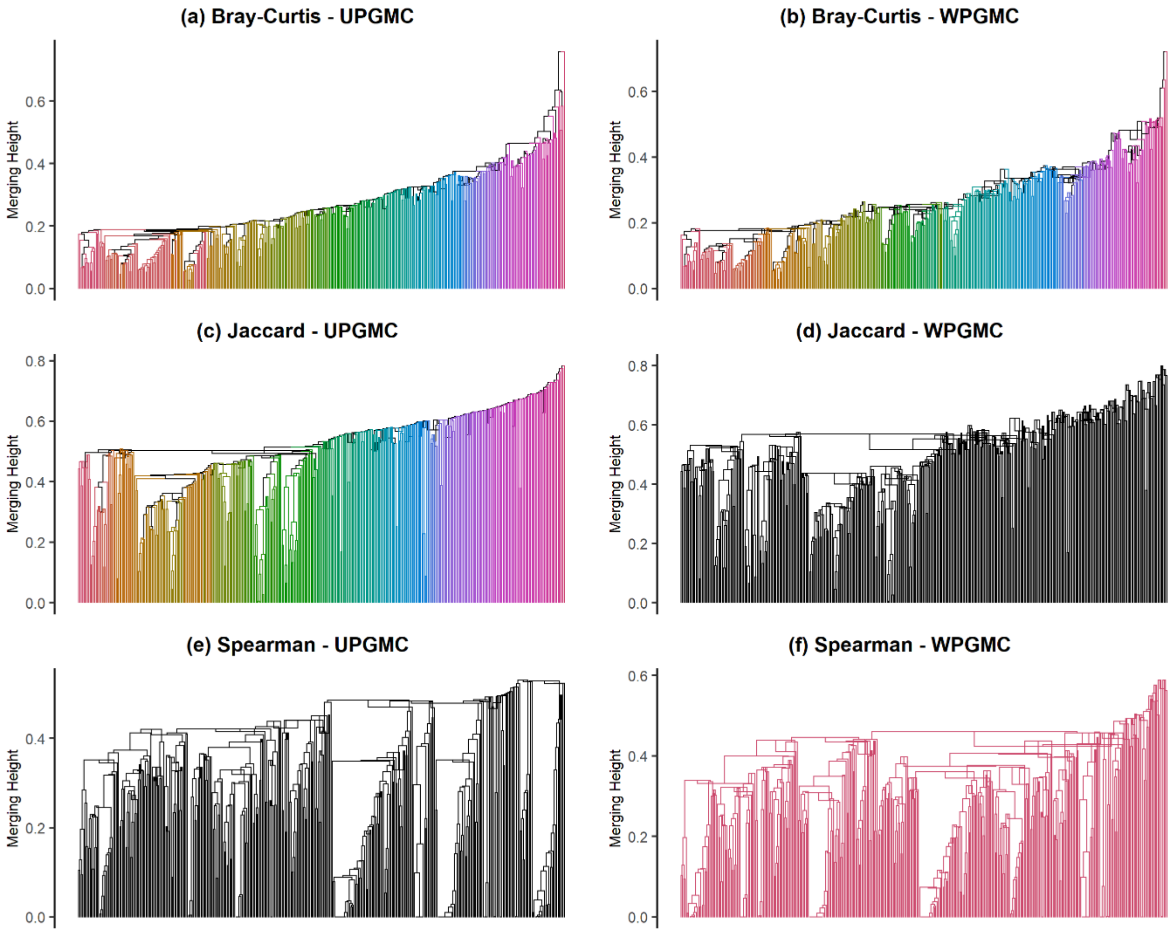


**Figure S4.** The presence of reversals, or the upward branching of nodes, among dendrograms resulting from UPGMC (a, c, e) and WPGMC (b, d, f). Example dendrograms here were based on Bray-Curtis dissimilarity (a, b), Jaccard index (c, d), and Spearman correlation (e, f). Reversals refer to branches that branched upwards, where the subsequent branching occurs at a merging height greater than the previous branching (i.e., the dendrogram structure is reversed). The number of clusters was arbitrarily set at $k = 6$, but clusters could not be properly delineated as a result of reversal. This forced $k > 6$ for the dendrogram in panels a, b, and c, $k = 0$ in panels d and e, and $k = 1$ in panel f.

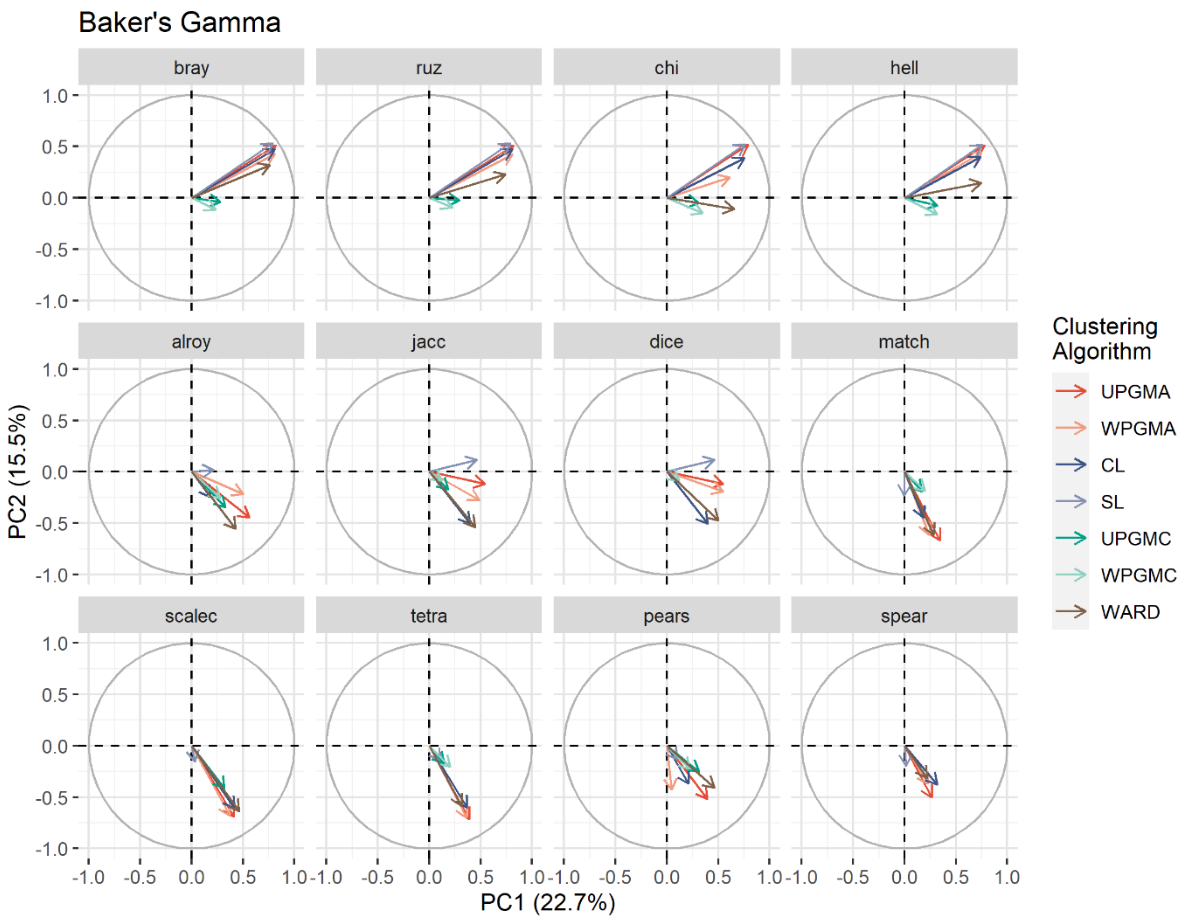


**Figure S5.** Dendrogram comparisons using Baker's Gamma to assess similarity and analysed using a PCA. Note, all facets represent vectors within the same PCA space, same dimensions, or same PC-axes, vectors among dendrograms based on different association indices were separated for visual clarity only.

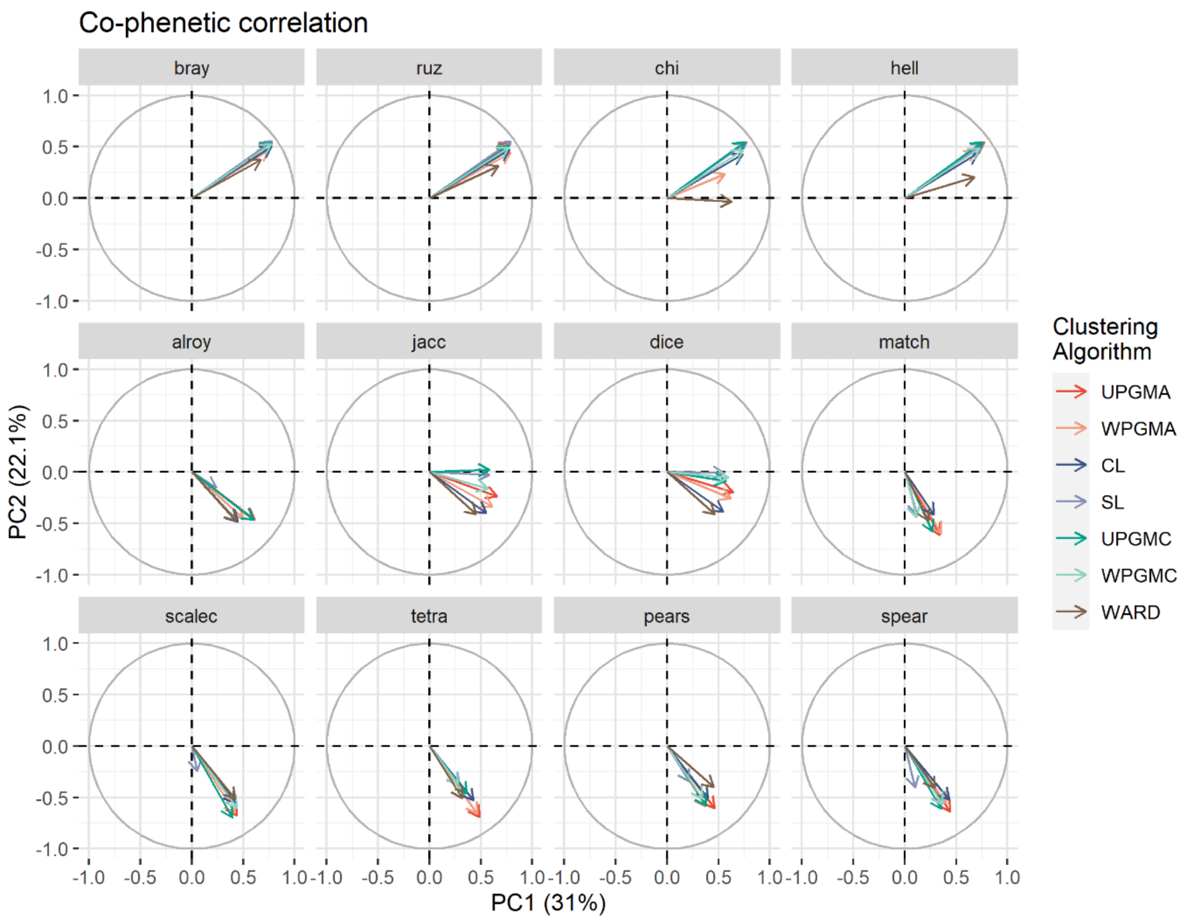


**Figure S6.** Dendrogram comparisons using Co-phenetic correlation to assess similarity and analysed using a PCA. Note, all facets represent vectors within the same PCA space, same dimensions, or same PC-axes, vectors among dendrograms based on different association indices were separated for visual clarity only.

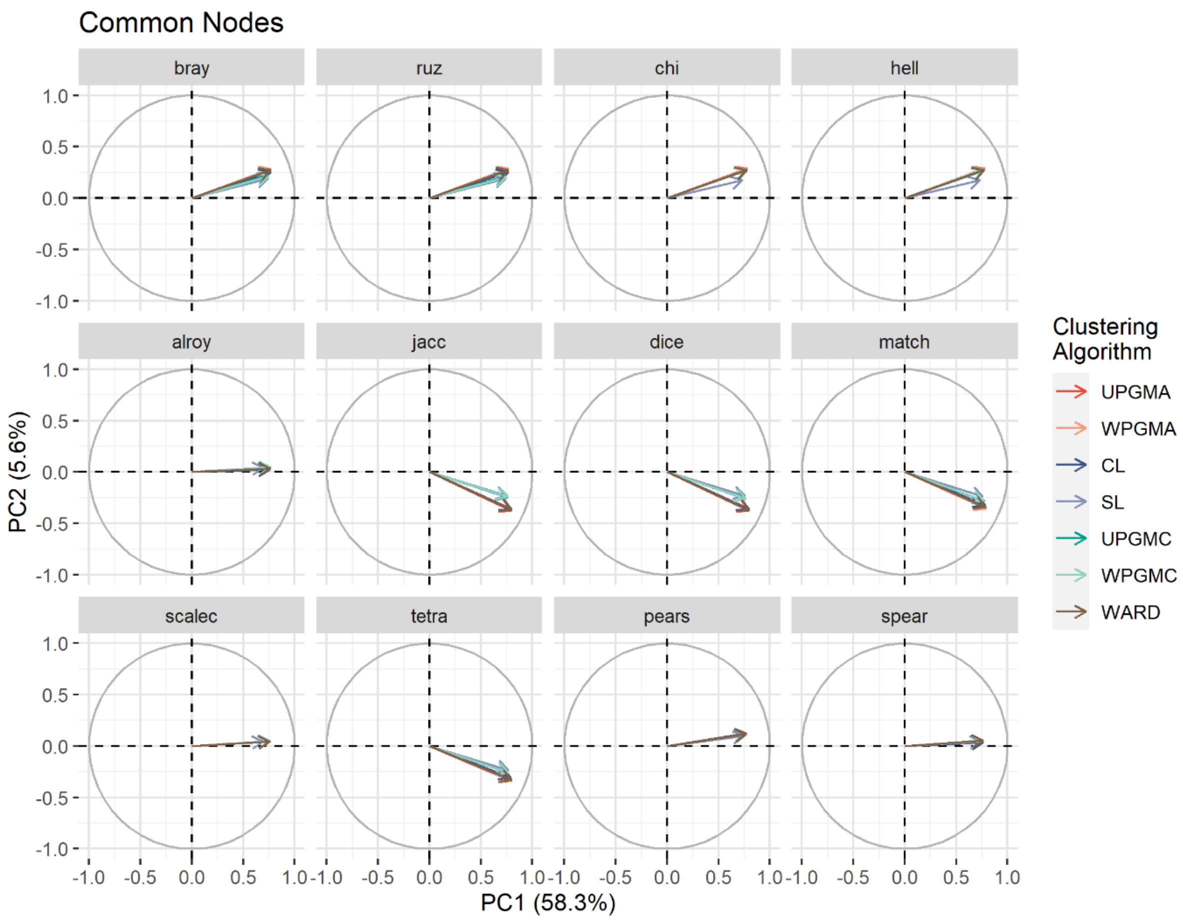


**Figure S7.** Dendrogram comparisons using Common Nodes to assess similarity and analysed using a PCA. Note, all facets represent vectors within the same PCA space, same dimensions, or same PC-axes, vectors among dendrograms based on different association indices were separated for visual clarity only.

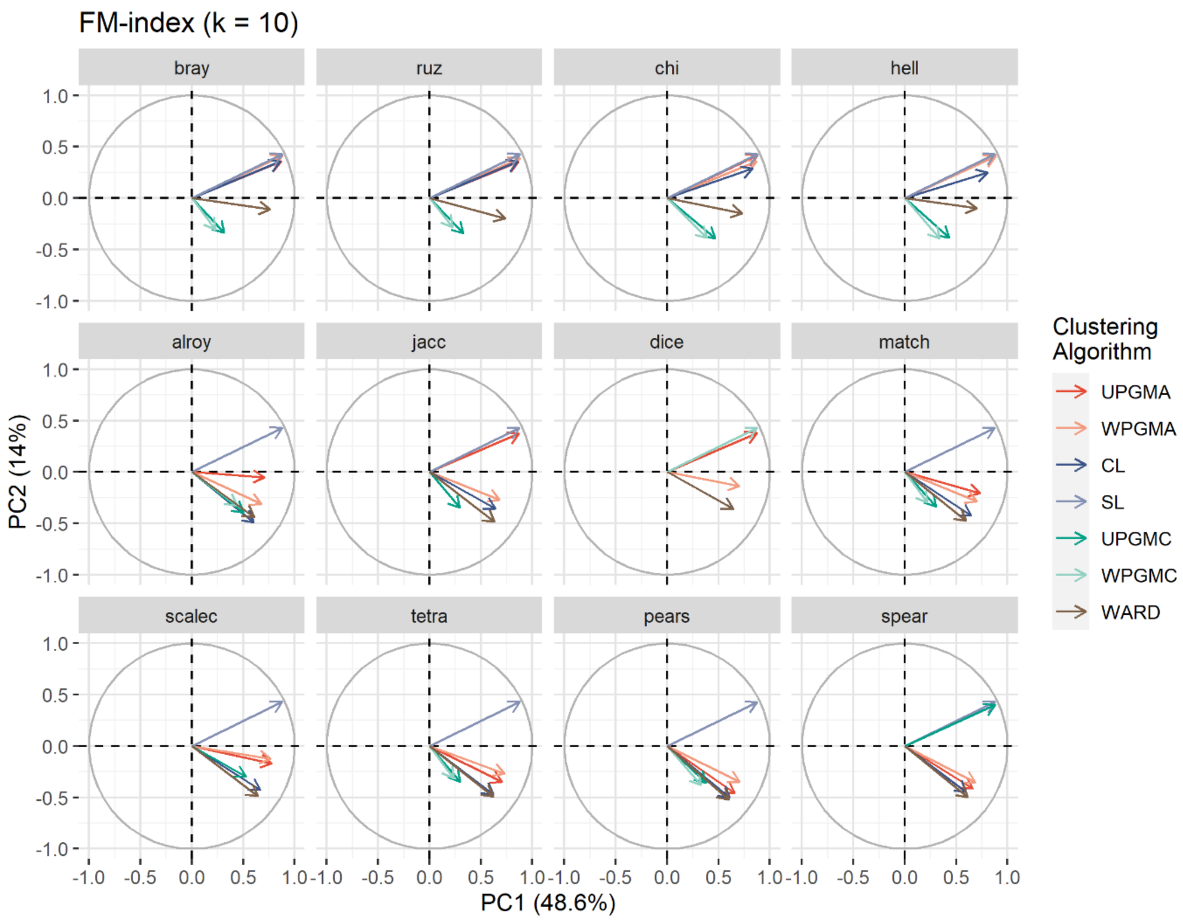


**Figure S8.** Dendrogram comparisons using FM-index (*k* = 10) to assess similarity and analysed using a PCA. Note, all facets represent vectors within the same PCA space, same dimensions, or same PC-axes, vectors among dendrograms based on different association indices were separated for visual clarity only.

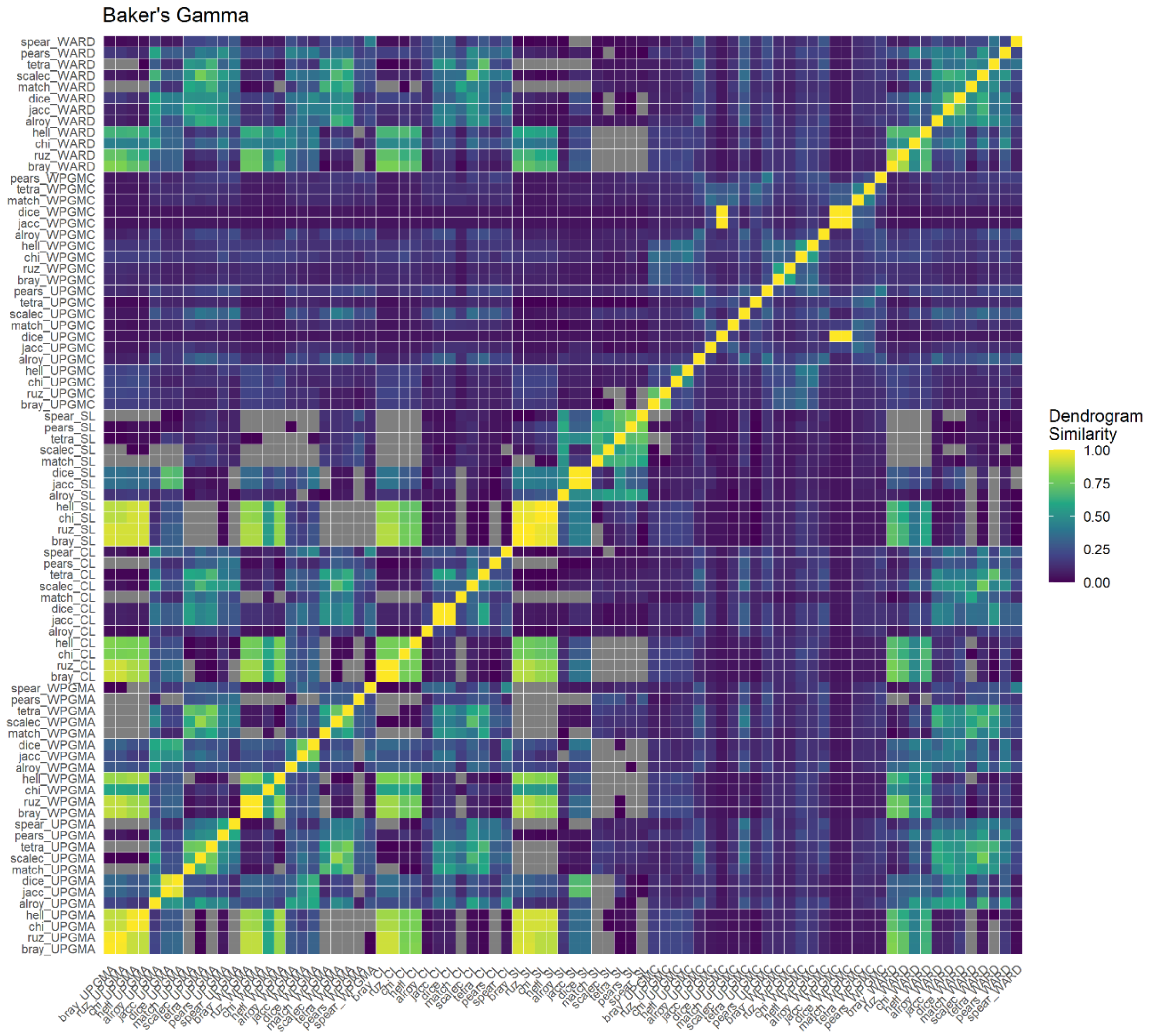


**Figure S9.** Correlation plot indicating dendrogram similarities based on Baker's Gamma (-1 to 1). For visual clarity, only values from 0 to 1 were shown, negative values were negligibly low. Some dendrograms may be missing from the plot because the presence of reversals may have prevented statistically comprehensible comparisons.

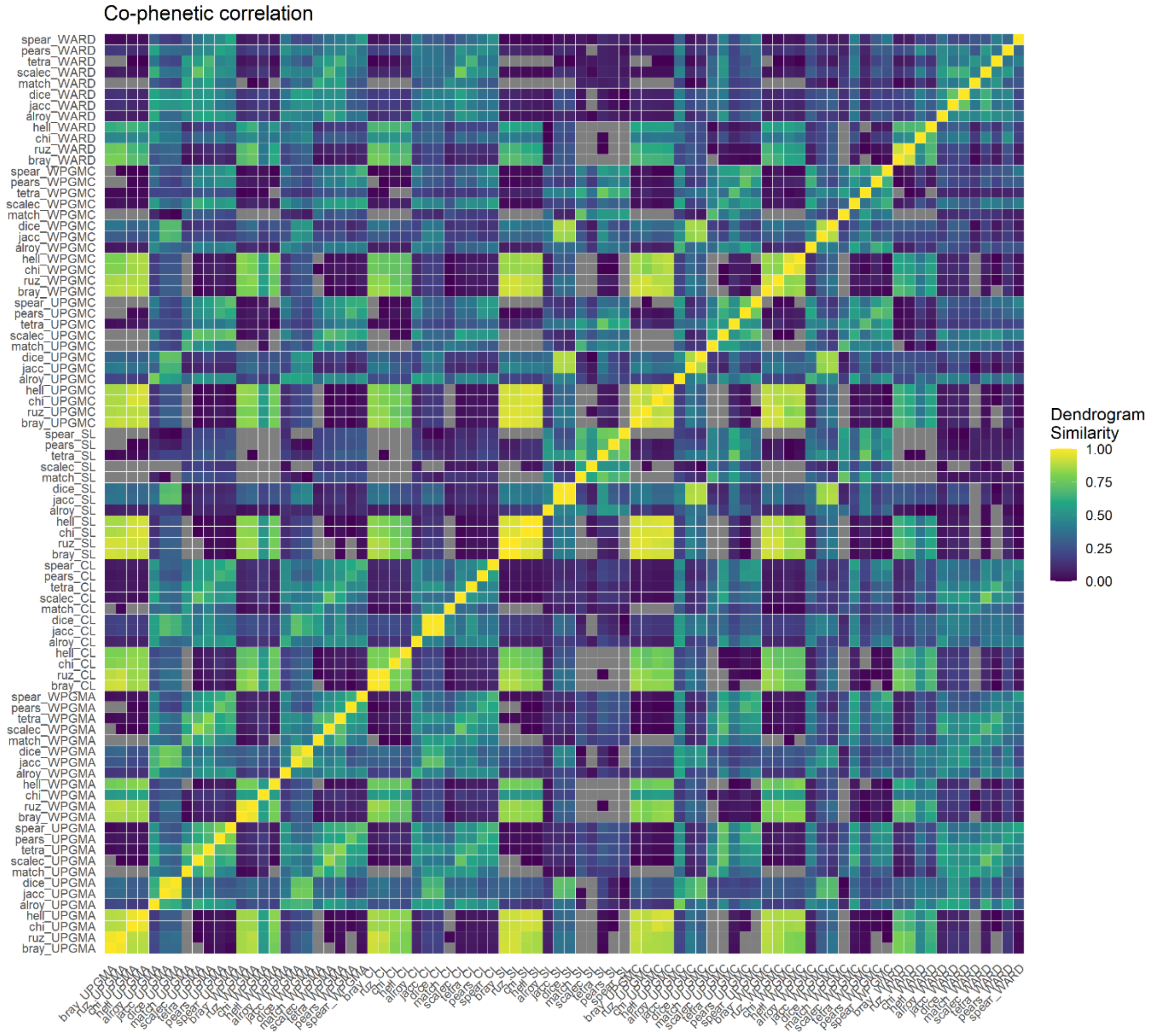


**Figure S10.** Correlation plot indicating dendrogram similarities based on Co-phenetic correlation (-1 to 1). For visual clarity, only values from 0 to 1 were shown, negative values were negligibly low. Some dendrograms may be missing from the plot because the presence of reversals may have prevented statistically comprehensible comparisons.

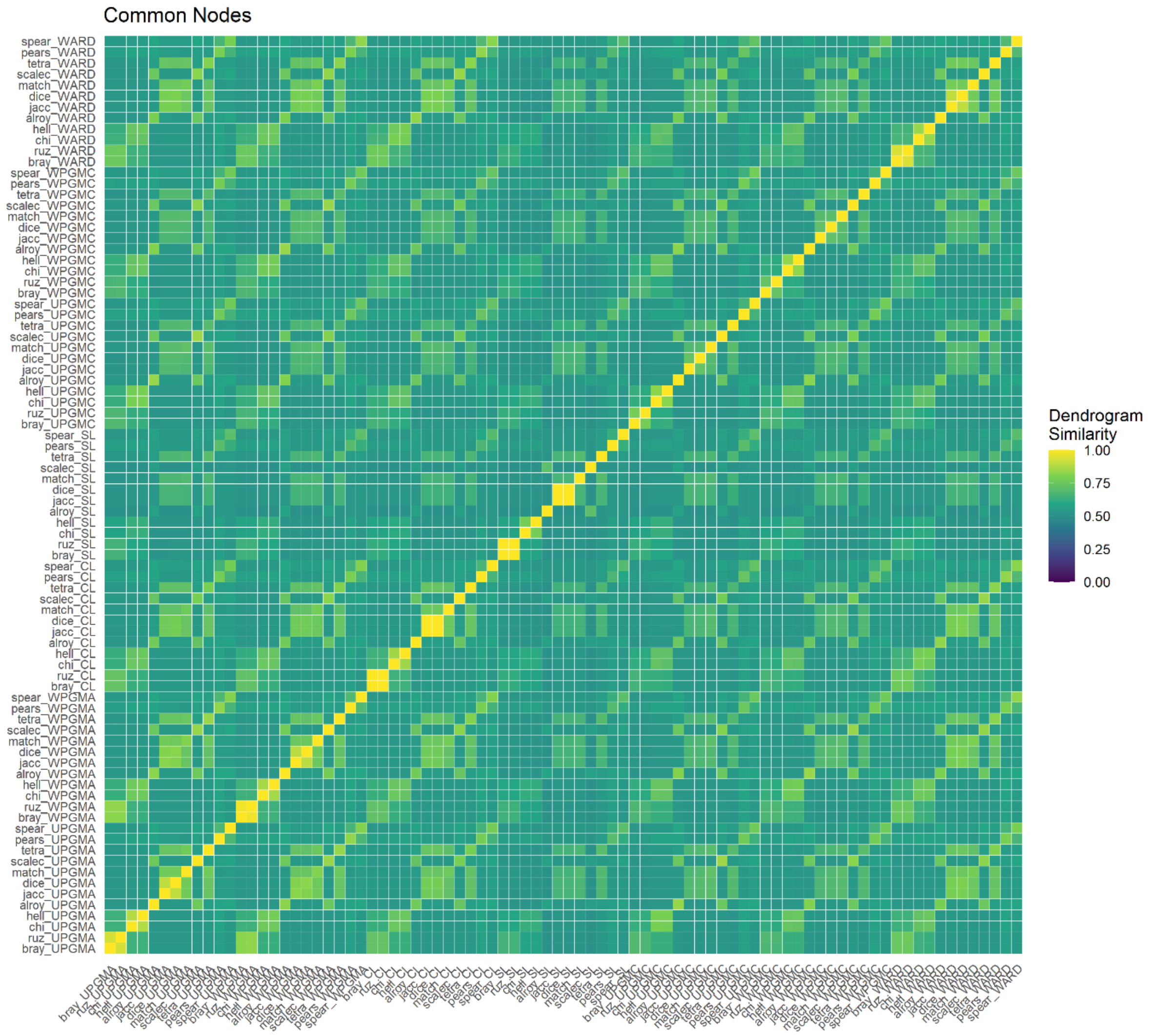


**Figure S11.** Correlation plot indicating dendrogram similarities based on Common Nodes (0 to 1). Some dendrograms may be missing from the plot because the presence of reversals may have prevented statistically comprehensible comparisons.

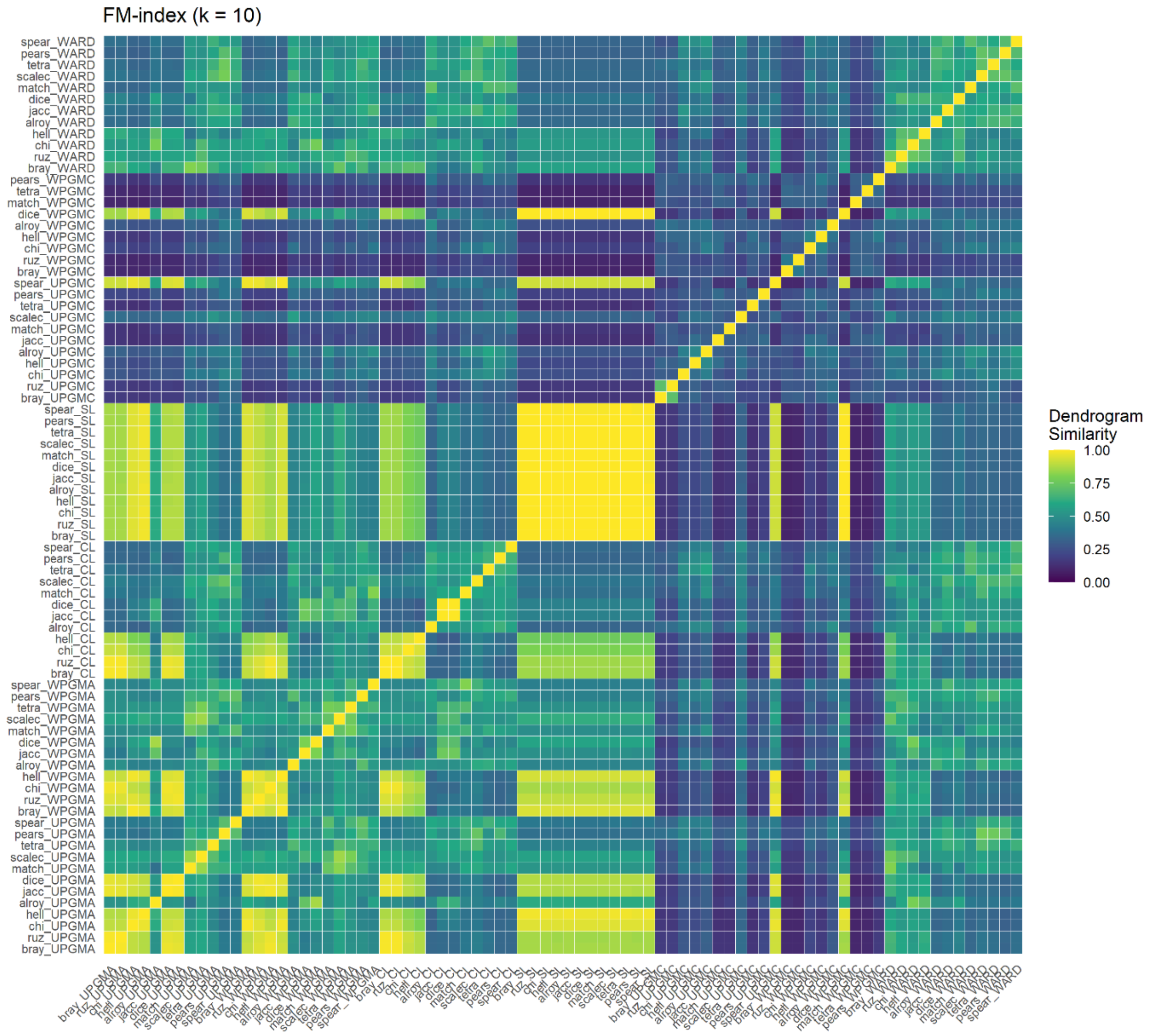


**Figure S12.** Correlation plot indicating dendrogram similarities based on FM-index (*k* = 10) (0 to 1). Some dendrograms may be missing from the plot because the presence of reversals may have prevented statistically comprehensible comparisons.

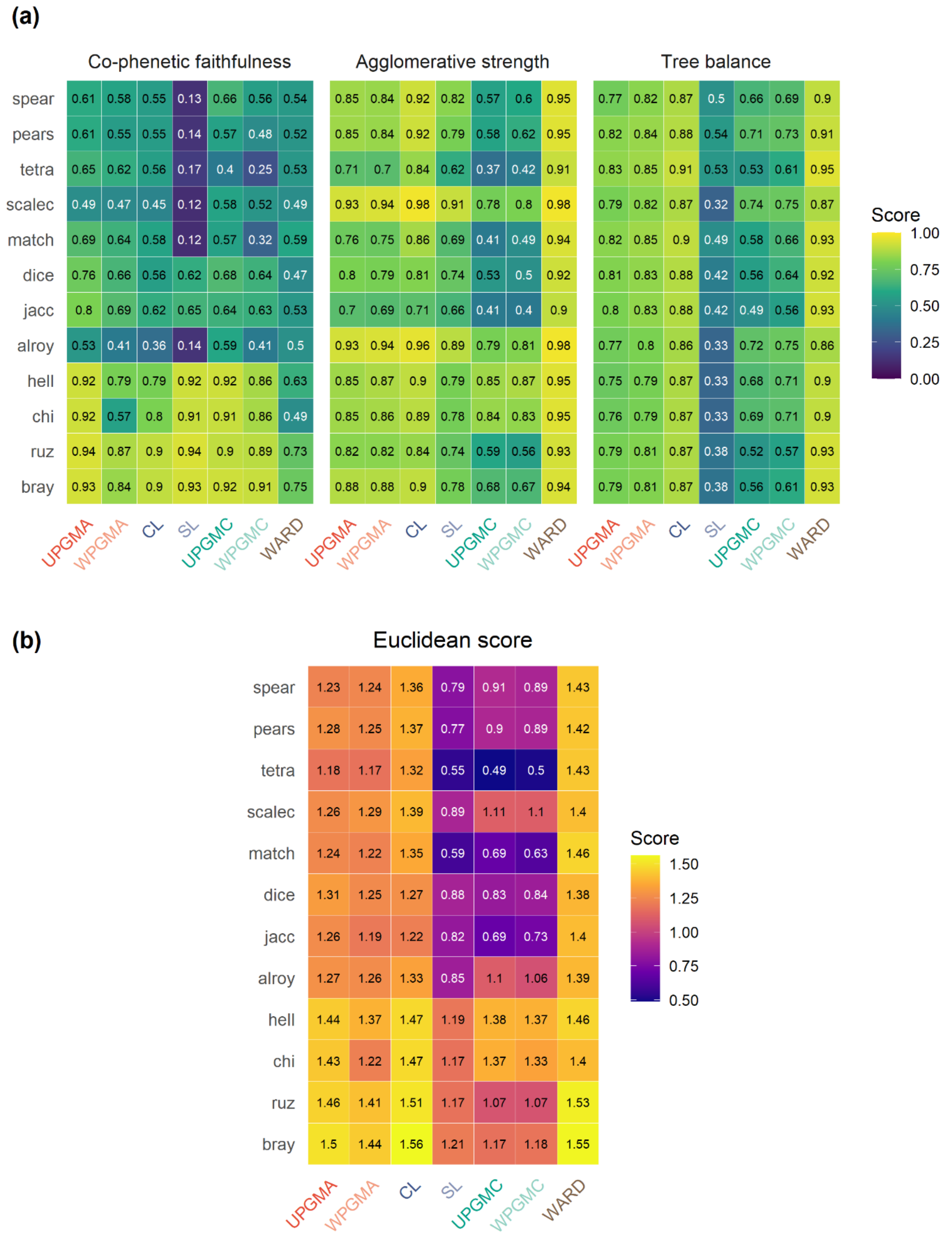


**Figure S13.** The performance of all candidate dendrograms. (a) Dendrogram performance based on the metrics co-phenetic faithfulness, agglomerative coefficient, and tree balance. (b) Dendrogram overall performance as the Euclidean distance (score) across the three metrics after applying a min-max scaling.

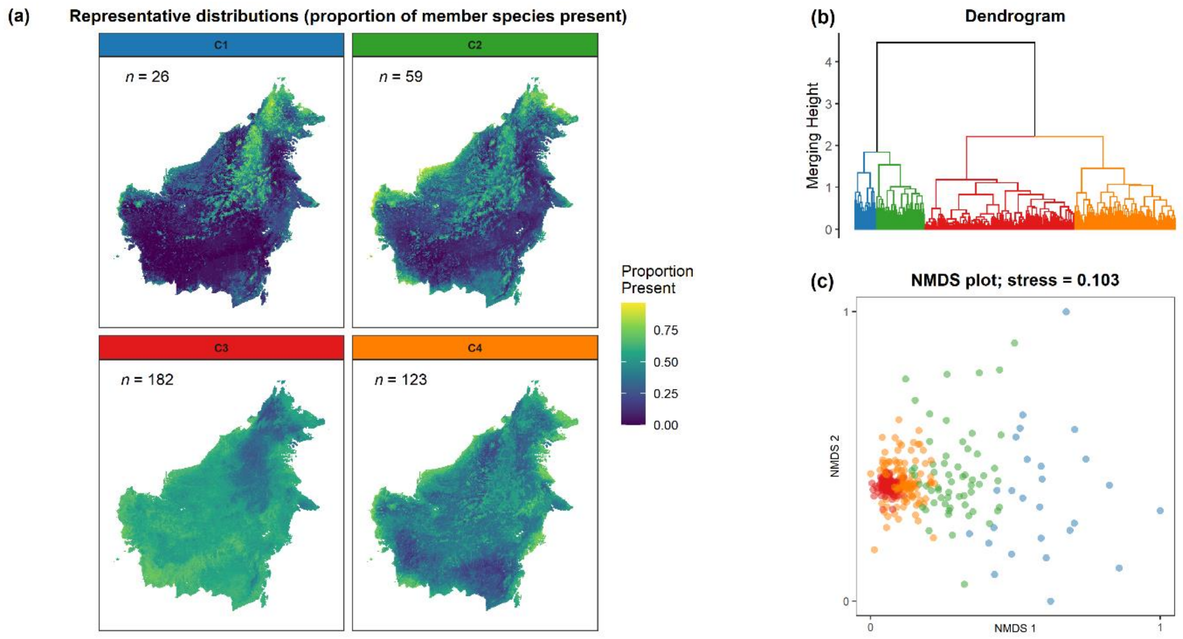


**Figure S14.** The visualisation of clusters at *k* = 4 for our final clustering outcome (Bray-Curtis — WARD). (a) The representative distribution of each cluster, where *n* equals the number of member species. (b) The dendrogram of the clustering outcome and (c) its underlying dissimilarity matrix visualised as an NMDS plot. Clusters were differentiated by colour, which were consistent across panels.

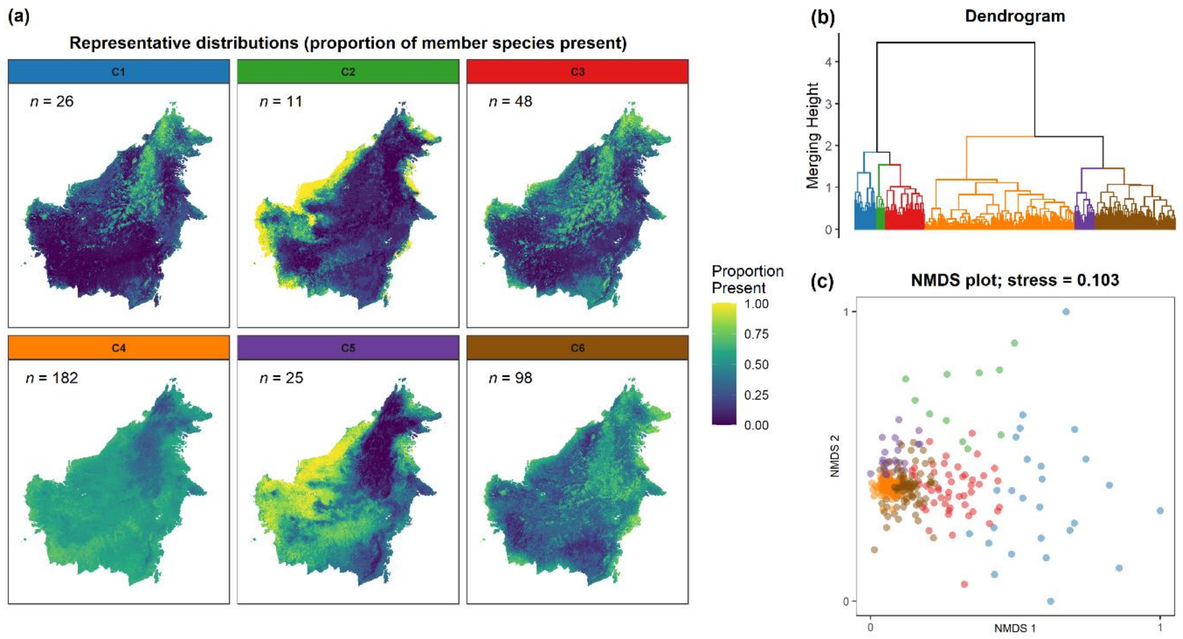


**Figure S15.** The visualisation of clusters at *k* = 6 for our final clustering outcome (Bray-Curtis — WARD). (a) The representative distribution of each cluster, where *n* equals the number of member species. (b) The dendrogram of the clustering outcome and (c) its underlying dissimilarity matrix visualised as an NMDS plot. Clusters were differentiated by colour, which were consistent across panels.

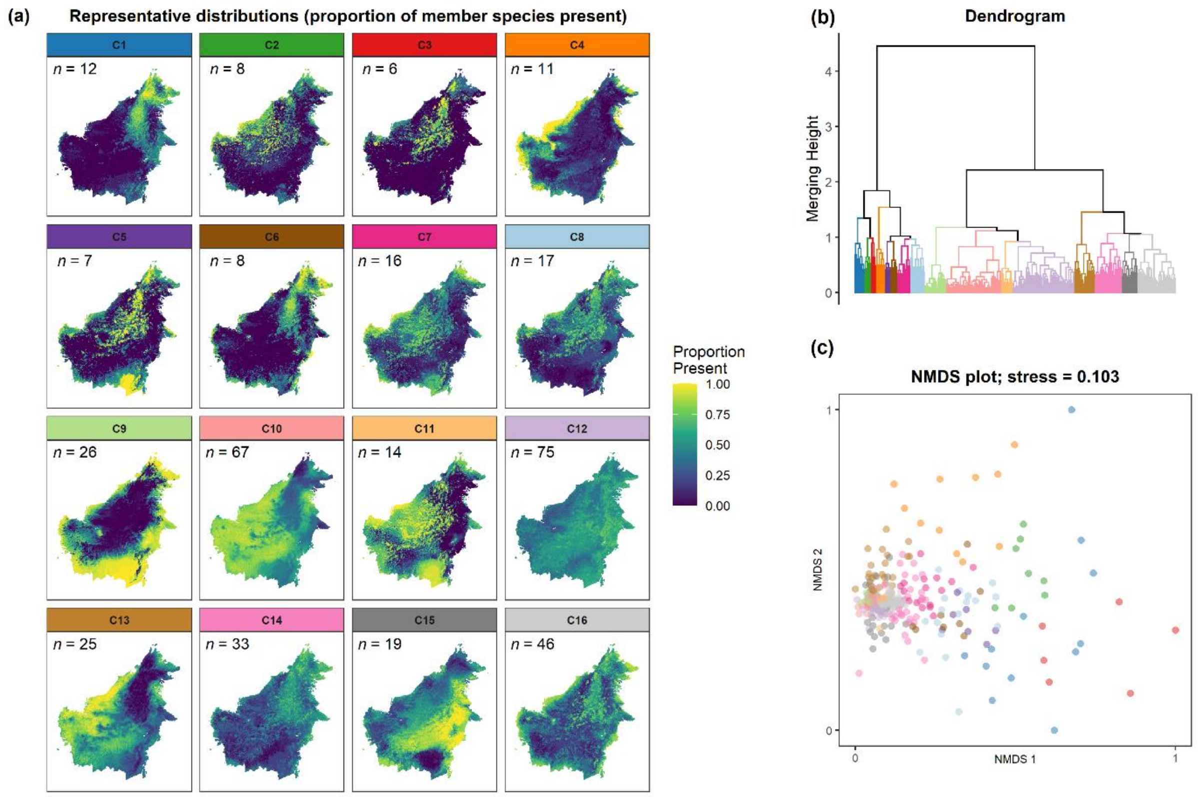


**Figure S16.** The visualisation of clusters at *k* = 16 for our final clustering outcome (Bray-Curtis — WARD); *k* =16 was derived by using the bifurcation paired T-test at a significance level of 0.01 (instead of 0.05) when aggregated distributions (centroids) were used as cluster centres. (a) The representative distribution of each cluster, where *n* equals the number of member species. (b) The dendrogram of the clustering outcome and (c) its underlying dissimilarity matrix visualised as an NMDS plot. Clusters were differentiated by colour, which were consistent across panels.

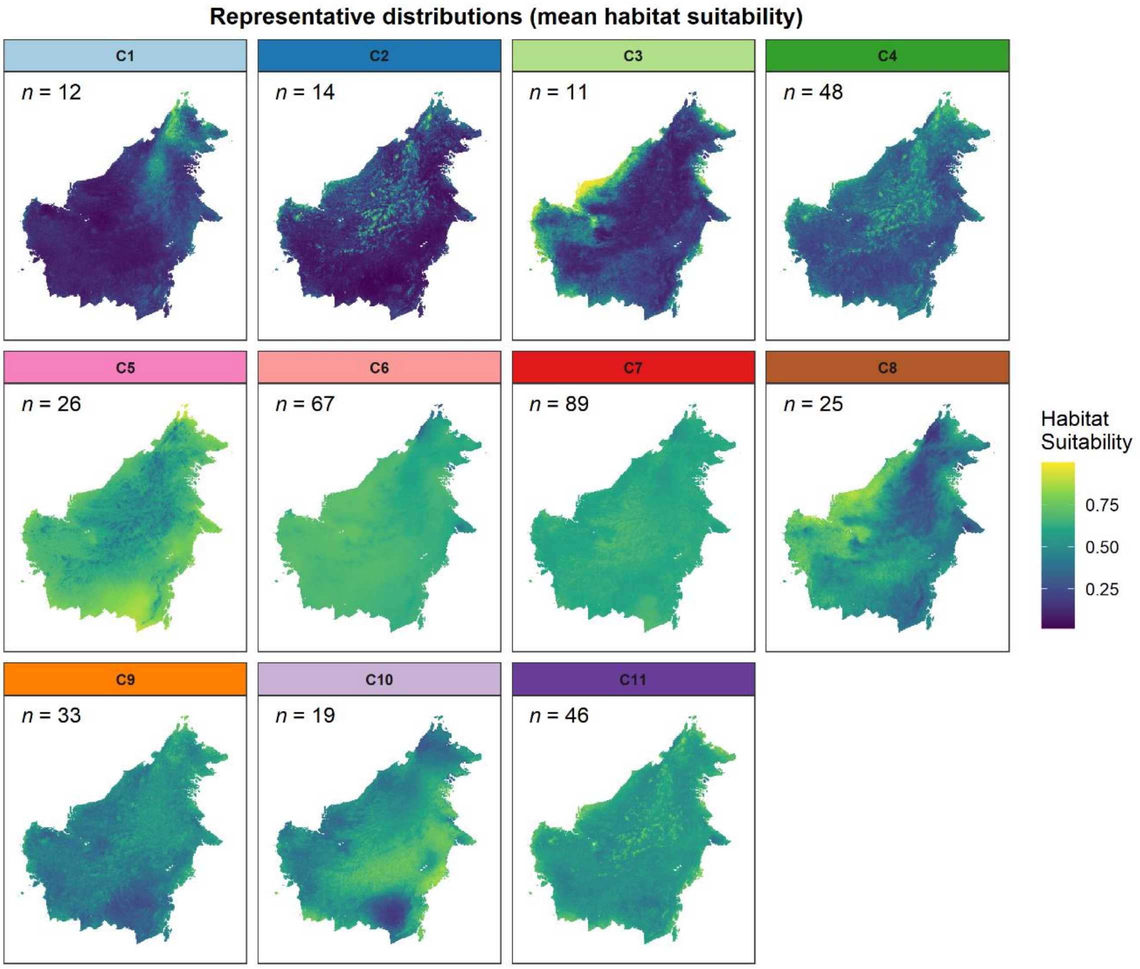


**Figure S17.** The aggregated distributions (centroid or mean habitat suitability) of the chosen clustering outcome (Bray-Curtis — WARD) for number of clusters $k$ = 11.

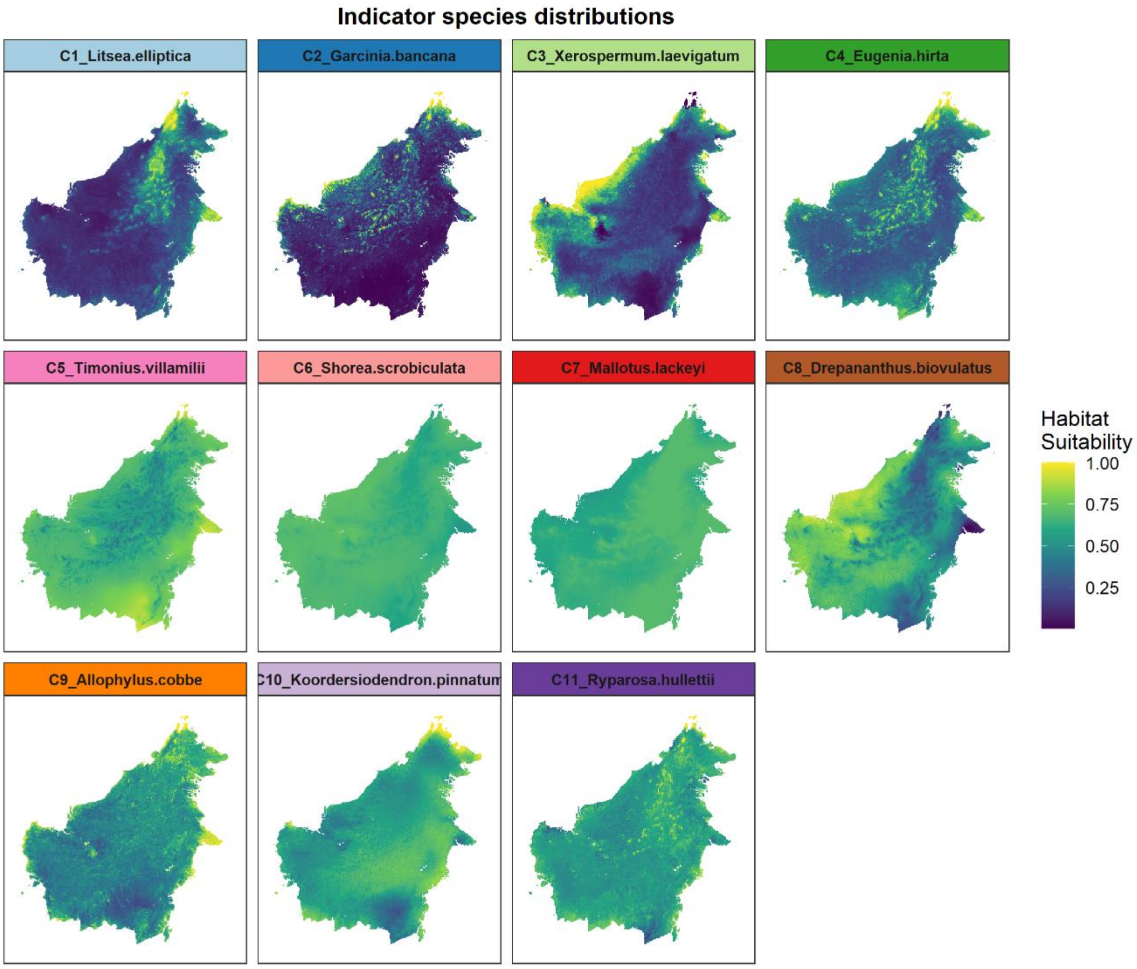


**Figure S18.** The indicator distributions (medoid or least dissimilar habitat suitability) of the chosen clustering outcome (Bray-Curtis — WARD) for number of clusters *k* = 11.

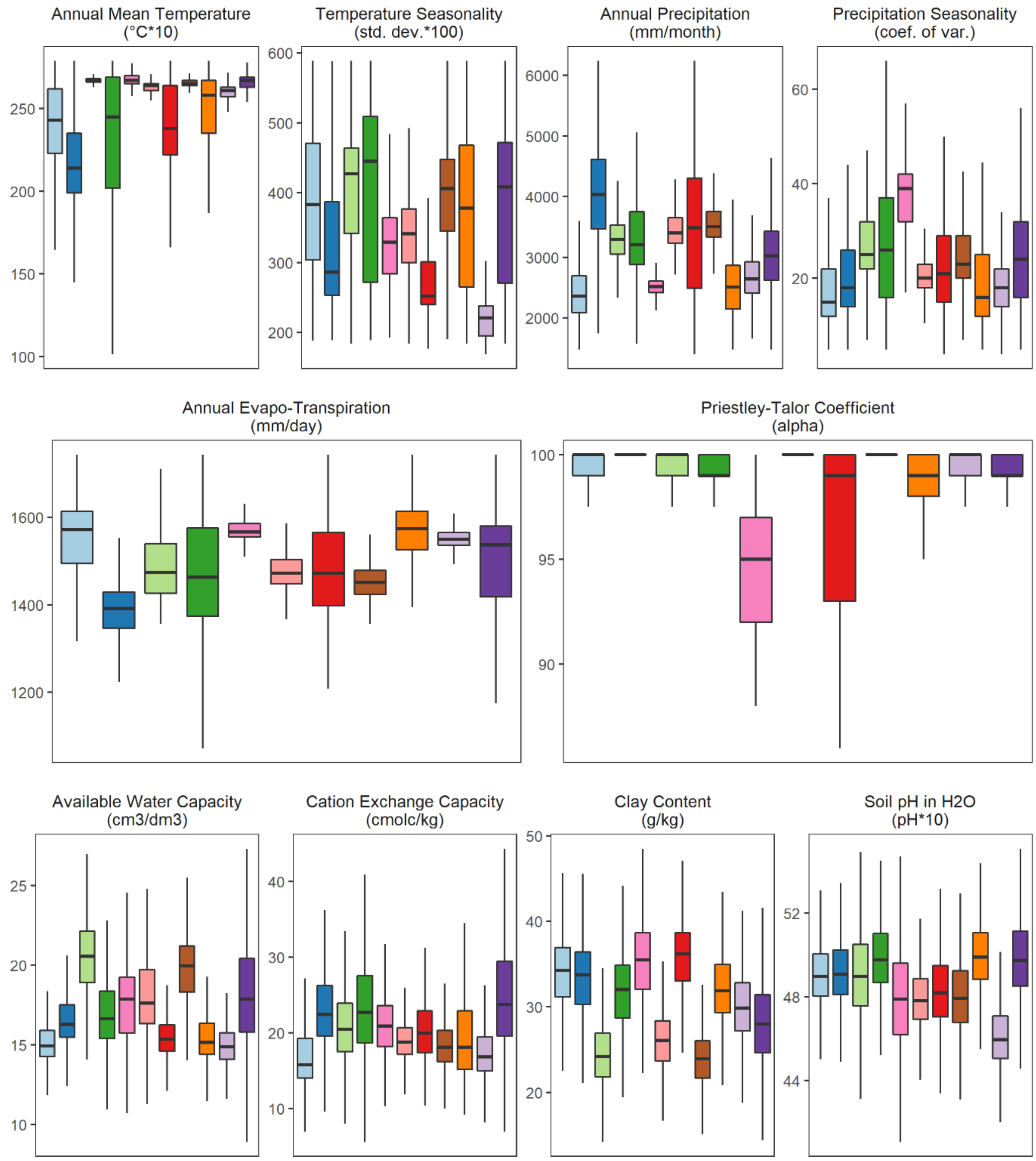


**Figure S19.** The environmental conditions that characterised each representative distribution. For each cluster's representative distribution, environmental conditions from the 95$^{th}$ percentile of pixels were extracted and further weighted based on the pixel's value (i.e., proportion of member species present) to derive the boxplots shown. Hence, the environmental characteristic of each cluster's representative distribution here represents the conditions of only the most suitable of sites (or where more of each cluster's member species is present).

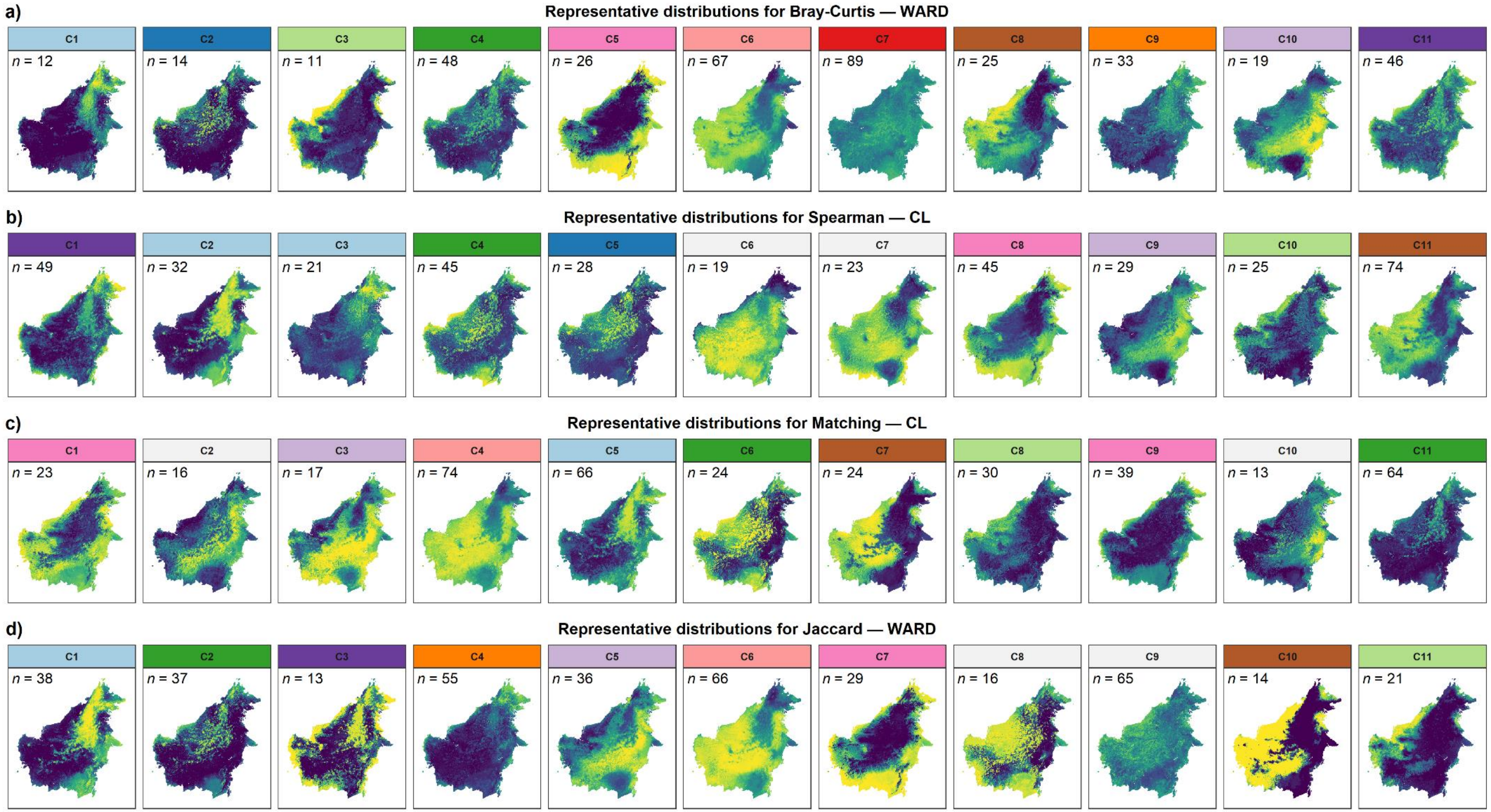


**Figure S20.** The representative distributions of our selected (a) and three other (b, c, d) well-performing clustering outcomes for $k$ = 11 (matching for our selected for easier comparison). Colours in (a) followed those in the main text while colours in (b, c, d) indicated the representative distribution in (a) that it was most similar to. Panels without colours (light grey) indicated representative distributions that were unique from those in (a). A combination of visual assessment and Pearson correlation was used to match distributions in (b, c, d) to those in (a).
